# The Landscape of Prostate Tumour Methylation

**DOI:** 10.1101/2025.02.07.637178

**Authors:** Jaron Arbet, Takafumi N. Yamaguchi, Yu-Jia Shiah, Rupert Hugh-White, Adriana Wiggins, Jieun Oh, Nicole Zeltser, Tsumugi Gebo, Adrien Foucal, Robert Lesurf, Chol-Hee Jung, Rachel M.A. Dang, Raag Agrawal, Julie Livingstone, Adriana Salcedo, Cindy Q. Yao, Shadrielle Melijah G. Espiritu, Kathleen E. Houlahan, Fouad Yousif, Lawrence E. Heisler, Anthony T. Papenfuss, Michael Fraser, Bernard Pope, Amar Kishan, Alejandro Berlin, Melvin L.K. Chua, Niall M. Corcoran, Theodorus van der Kwast, Christopher M. Hovens, Robert G. Bristow, Paul C. Boutros

## Abstract

Prostate cancer is characterized by profound clinical and molecular heterogeneity. While its genomic heterogeneity is well-characterized, its epigenomic heterogeneity remains less understood. We therefore created a compendium of 3,001 multi-ancestry prostate methylomes spanning normal tissue through localized disease of all grades to poly-metastatic disease. A subset of 884 samples had multi-omic DNA and/or RNA characterization. We identify four epigenomic subtypes that risk-stratify patients and reflect distinct evolutionary trajectories. We demonstrate extensive regulatory interplay between DNA copy number and methylation, with transcriptional consequences that vary across genes and disease stages. We define epigenetic dysregulation signatures for 15 important clinico-molecular features, creating predictive models for each. For example, we identify specific epigenetic features that predict patient outcome and are synergistic with clinical prognostic features. These results define a complex interplay between tumour genetics and epigenetics that converges to modify gene-expression programs and clinical presentation, in part through modulation of epigenetic aging.

## Introduction

Over 280,000 men were diagnosed with prostate cancer in the United States in 2023; it was the second leading cause of cancer mortality amongst men (1). Prostate cancer is highly heritable, with ∼60% of variability in incidence being explained by rare and common germline genetics (2). While the majority of new diagnoses occur when the disease is localized and thus potentially curable, treatment outcomes vary widely (3). Current clinical prognostic factors only partially explain the heterogeneity in clinical outcomes to first-line local and systemic therapies (4). Most localized disease follows a relatively indolent clinical course, but once prostate cancer has metastasized and become resistant to anti-androgen therapies, it is incurable (5).

This clinical heterogeneity is matched by extensive genomic heterogeneity. Localized disease in particular exhibits extensive intra- and inter-tumoural variability, which is associated with different clinical outcomes (6–10). Somatic genetic heterogeneity is observed alongside extensive epigenomic and gene-expression variability (7,11–16). Aberrant methylation of <u>CpG</u> Islands (CGI) in gene promoter regions is associated with transcriptional inactivation of tumour suppressor genes, oncogene activation and genome instability (17–20). A hypermethylated subtype has been identified in numerous cancers including prostate cancer (21–23). Methylation of hundreds of loci is associated with adverse clinical outcomes (8,24–27) and is additive or synergistic with clinically used biomarkers (28). RNA methylation drives aberrant gene-expression and prostate cancer progression (29). While informative, these studies have been limited in sample size and clinical data, and do not cover the full natural history of the disease. As a result, we have little understanding of how DNA methylation interacts with tumour genetics to shape clinico-molecular phenotypes across disease stages.

To fill this gap, we created a compendium of DNA methylation from 3,001 prostate specimens: 2,355 from cancers and 646 from non-cancerous prostate tissue. The tumours represent the full range of risk groups within both localized and metastatic disease, and arise in individuals with a range of ancestries. We identify distinct DNA methylation subtypes associated with disease stage and evolution. This allows us to quantify how DNA methylation and copy number jointly regulate gene-expression, and how this regulation evolves during progression. We comprehensively define the epigenomic and transcriptomic consequences of somatic driver mutations, and demonstrate that these accurately predict clinical outcomes. Since tumour progression reflects not only genomic alterations but also pathway states, we systematically examined pathway activity across methylation subtypes, including in tumours with recurrent driver events such as *PTEN* loss. In particular, we focused on hypoxia and related programs such as angiogenesis and glycolysis because these processes are closely linked to tumour progression and treatment resistance. Together, these analyses show that the interplay between genetic and epigenetic variability rationalizes the clinical heterogeneity characteristic of prostate cancer.

## Results

### Methylation subtypes of prostate cancer

We created a cohort of 3,001 prostate tissue methylomes from 14 published and one unpublished cohorts (7,8,13,15,25,30–38). This compendium represents the full spectrum from normal tissue (22%; n = 646) to localized cancer (70%; n = 2,102) to castrate-resistant metastatic disease (8%; n = 253). Localized cancers represent all five ISUP grade groups (GG) with 117 to 403 patients (**Supplementary Figures 1a-e**; **Supplementary Tables 1-3**).

We first derived comprehensive methylation subtypes in the largest single cohort, The Cancer Genome Atlas Prostate Adenocarcinoma (TCGA-PRAD), comprising 498 tumours and 50 normal prostate samples, then verified them in the remaining 14 cohorts. Expanding on previous smaller subtyping studies (7,32,39,40), we define four <u>m</u>ethylation <u>s</u>ubtypes, termed MS-1 through MS-4 (**Figure 1a**; **Supplementary Figure 1f**). To allow these subtypes to be readily applied to new samples, we identified 5,486 subtype-defining CpGs and created a shrunken centroid classifier for single-sample subtype-assignment (**Supplementary Figure 1g**; **Supplementary Figure 2; Supplementary Table 4**). We implemented this in a new open-source R package called *PrCaMethy* to facilitate application and extension of all methods and signatures reported here.

**Figure 1:**
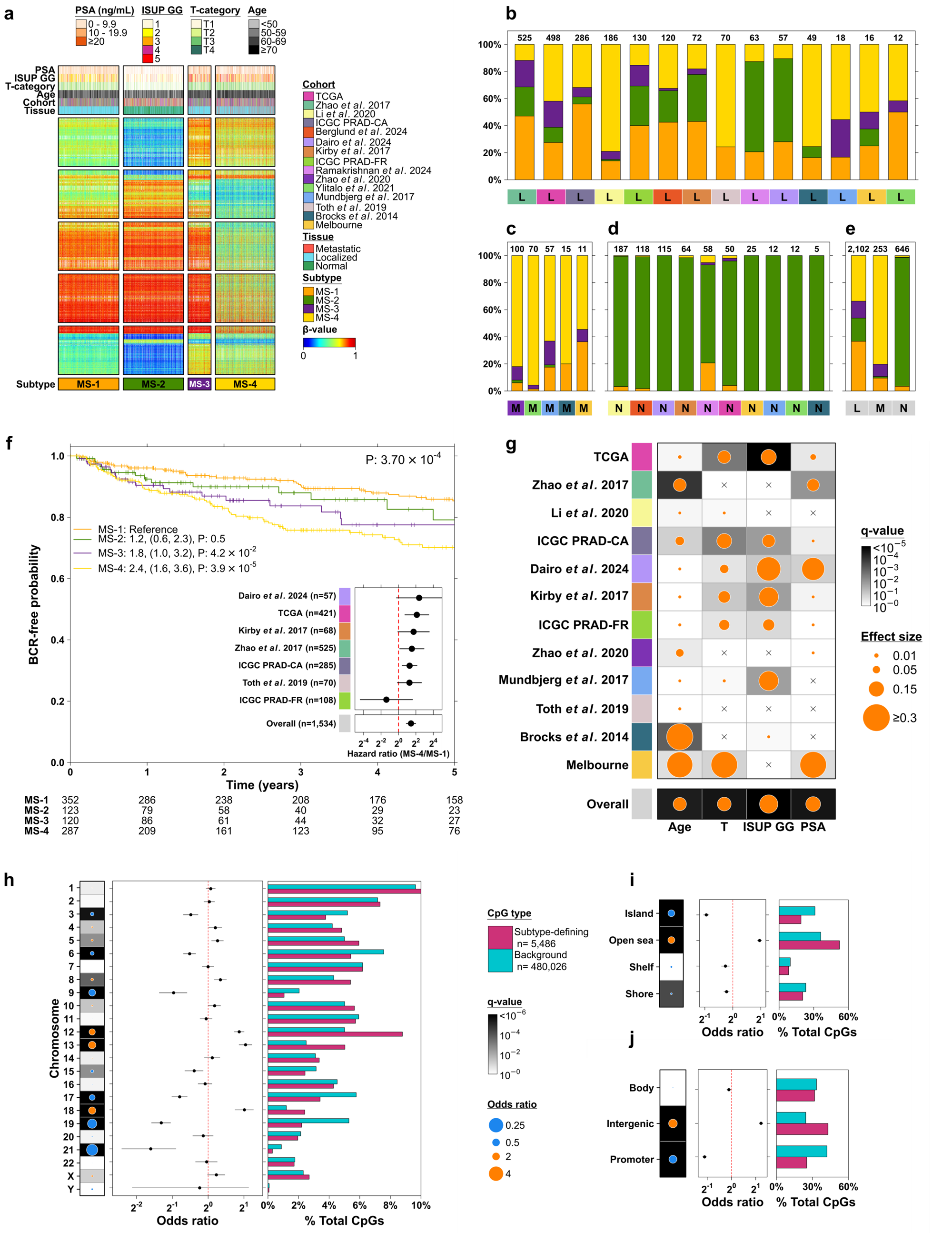
The epigenomic landscape of prostate cancer. **a.** Subtype methylation profiles of 2,355 prostate tumours and 646 normal samples from 15 cohorts, including clinical pre-treatment variables. The heatmap includes 5,486 CpGs which were selected by IMPACC as important for deriving the subtypes. CpGs were sorted using hierarchical clustering with complete linkage and Euclidean distance, then split into five equally sized groups for visualization. Missing values are colored in white and clinical variables are not shown for normal samples (other than age). **b-e.** For each cohort (vertical bar), the percentage of samples across subtypes is shown stratified by localized prostate tumour samples (**b**), metastatic samples (**c**), normal samples (**d**), and when pooling all cohorts together (**e**). The total sample size within each cohort is given above the bars. L: localized, M: metastatic, N: normal sample. **f.** Kaplan-Meier curves were used to estimate BCR-free probabilities separately in each subtype, pooling all patients with available BCR data (n = 882). The p-value in the top-right corner is from a log-rank test. In the upper-left plot, hazard ratios with 95% confidence intervals and Wald-test p-values were estimated using a Cox proportional hazards model treating MS-1 as the reference group and adjusting for cohort. The table below the plot gives the number of patients at risk from 0-5 years. The bottom-right forest plot is a meta-analysis of 7 cohorts with BCR-related outcomes (see **Methods** for details). The “Overall” row contains the overall meta-analysis hazard ratio for comparing MS-4 to MS-1. Dots represent the estimated hazard ratio with 95% CI given by the adjacent lines. **g.** The dot-map summarizes the overall association of subtypes with four clinical features: age, T-category, ISUP GG and PSA when pooling all patients together (Overall) and separately by cohort. Each of the clinical variables was categorized as in **Supplementary Table 1** and tested for an association with the four methylation subtypes using Fisher’s Exact Test. The size of the dots reflects the bias-corrected Cramér’s V effect size and the background color represents the q-value, which adjusts for multiple testing separately for each clinical variable. An x within a cell means that feature was unavailable for that cohort. **h-j.** Fisher’s exact test and odds ratio were used to compare the rates of subtype-defining and non-subtype-defining “background” CpGs within each chromosome (**h**), within different types of CpGs (island, open sea, shelf, and shore) (**i**) and within gene body, intergenic and gene promoter regions (as defined in Methods) (**j**).

The four methylation subtypes are tightly associated with clinical features of the disease. For example, subtype MS-2 is the “normal-like” subtype. It preferentially occurs in low-grade localized disease (64% ISUP grades 1 or 2; **Figure 1b**), is extremely rare in metastatic disease (1%; **Figure 1c**), and dominates in non-malignant prostatic tissue (95%; **Figure 1d**). Subtype MS-4 showed the inverse pattern, becoming most common in aggressive disease: 1% of non-malignant tissue *vs.* 34% of localized disease *vs.* 80% of metastatic disease. Subtypes were not driven by tumour purity, self-identified race (SIR), nor genetic ancestry (**Supplementary Figure 3a, Supplementary Information**) and differed dramatically between disease stages (p < 10^-45^; **Figure 1e**; **Supplementary Figure 3b**).

For patients with localized disease, subtypes MS-1 and MS-2 had the lowest risk of biochemical recurrence (BCR), while MS-4 had the highest risk (HR = 2.4; p = 3.9 x 10^-5^; **Figure 1f**). MS-4 remained independently prognostic after adjusting for cohort, age, pathologic T-category, ISUP GG, pre-treatment serum prostate-specific antigen (PSA) and total <u>c</u>opy <u>n</u>umber <u>a</u>lteration (CNA) burden (adjusted HR = 2.01, p = 0.0078). This pooled analysis was confirmed by meta-analysis of the seven cohorts with BCR-related outcomes: across 1,534 patients showed MS-4 had 2.7 times greater risk of relapse than MS-1 (**Figure 1f** inset) with no evidence of heterogeneity in effect sizes between cohorts (p = 0.52). Methylation subtypes were associated with pre-treatment prognostic features including tumour size and extent (T-category), ISUP GG and serum PSA abundance, with consistent effect-sizes across cohorts and in the full compendium (q-values < 0.05, **Figure 1g**; **Supplementary Figure 3k-n**).

These MS subtypes showed significant concordance with the previously identified CpG Island Methylator Phenotype (CIMP) in the TCGA-PRAD cohort (23) (Qannari test p < 0.001, Adjusted Rand Index, 95% CI = 0.13-0.15; **Supplementary Figure 4a**). CIMP-samples were predominantly assigned to MS-1 or MS-2 (84%; 78/93), and CIMP+ samples were predominantly assigned to MS-3 or MS-4 (88%; 52/59). We then computed CIMP scores for all 3,001 samples (**Supplementary Table 4**). Across the full cohort CIMP scores significantly varied by MS subtype even after adjustment for platform and cohort (Kruskal-Wallis, p = 9.9 x 10^-277^, ε² = 0.43, 95% CI = 0.40 - 0.46). MS-3 and MS-4 showed elevated CIMP scores and MS-2 had the lowest (pairwise Mann-Whitney U-tests, q < 0.05; **Supplementary Figure 4b)**. Several clinico-molecular features including ISUP GG and T-category were associated with CIMP scores (Kruskal-Wallis q < 0.05; **Supplementary Figure 4c-i**). Several key driver mutations were associated with increased CIMP, but notably *not TMPRSS2*-*ERG* fusion events (**Supplementary Figure 4j**).

The vast majority of subtype-defining CpGs (5,407/5,486; 98.6%) showed significant tumour-normal differences (U-test q-value < 0.05; **Supplementary Figure 5a**). Subtype-defining CpGs were modestly more likely to differ between tumour and normal samples than non-subtype-defining CpGs (relative risk = 1.16; p = 7.2 x 10^-259^). Similarly, 96.3% of subtype-defining CpGs showed significant differences between localized and metastatic samples, compared to 80.8% of non-subtype-defining CpGs (relative risk = 1.19; p = 9.0 x 10^-257^; **Supplementary Figure 5b**). Across the four methylation subtypes, these CpGs also showed distinct β-value distributions, with all pairwise comparisons significant when MS-2 was represented using normal samples (**Supplementary Figure 5c**). Subtype-defining CpGs were significantly enriched on five chromosomes and depleted on seven (**Figure 1h**), and preferentially occurred in open sea rather than islands or shores (**Figure 1i**) and in intergenic regions rather than promoters or gene bodies (**Figure 1j**).

### Tumour CNAs drive regional methylation & gene-expression

These positional biases in the genomic locations of subtype-defining CpGs suggested an association between copy number state and methylation status (8,41–43). We therefore considered the 870 tumours in our compendium with matched CNA profiling. Methylation was predominantly inversely correlated with copy number states, with 86% of genes showing 3 correlations (**Figures 2a-b**). Given the inability of array-based technologies to quantify allele-specific methylation status, it remains unclear if the average methylation per allele differs as a function of the number of copies. Methylation of promoter CpG islands was strongly associated with decreased transcript abundance, with negative correlations for 92% of genes (**Figure 2c**).

**Figure 2:**
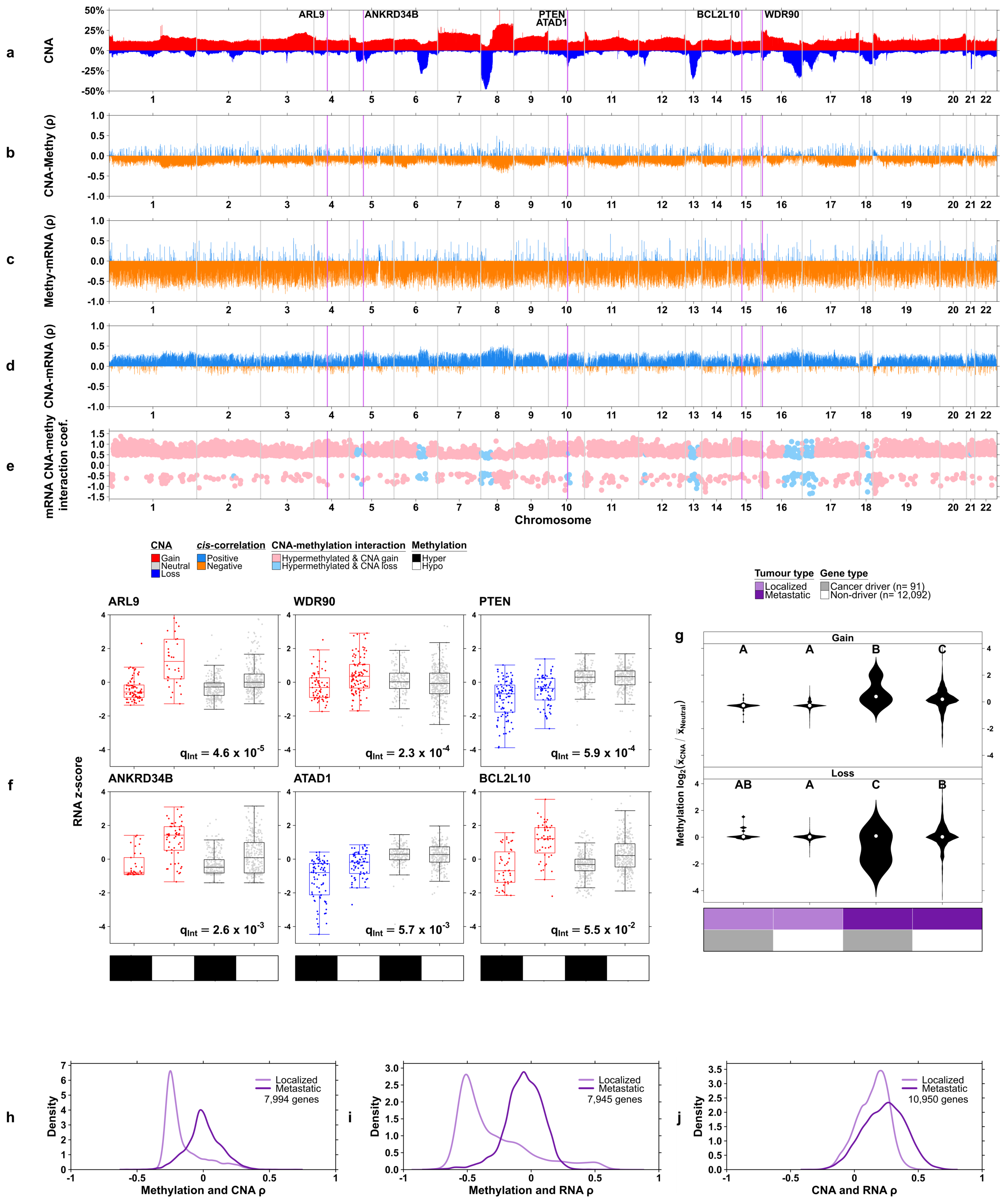
The regulatory landscape of prostate cancer. **a.** The y-axis indicates the percent of patients with copy number gain (red bar) or loss (blue bar) within a gene at that location of the genome (x-axis). The vertical magenta bars indicate genes later shown in (**f**). **b-d.** Spearman’s correlation of methylation and CNA (**b**), methylation and mRNA (**c**), and CNA and mRNA (**d**) within each gene. Positive correlations are shown in blue and negative correlations in orange. Gene-level methylation was measured for each patient by the median β-value among CpG islands in the gene promoter region. **e.** Linear regression interaction coefficient of binary methylation (hyper *vs.* hypomethylation; median dichotomized) and binary CNA status (loss *vs.* neutral if loss was the more common mutation, otherwise gain *vs.* neutral). The dependent variable is RNA abundance (z-score transformed log_2_ TPM). Genes with significant interaction effects (q < 0.05) are shown as well as gene *BCL2L10* (q = 0.055). **f.** Examples of methylation-CNA interactions on mRNA abundance for six genes. The q-value for testing the interaction effect is indicated by q_int_. Gene-level methylation was measured for each patient by the median β-value among CpG islands in the gene promoter region and then median dichotomized into hyper *vs.* hypomethylation. **g.** For each gene, the predominant CNA status was used when CNA status varied across samples (the more prevalent of gain or loss). Then the median methylation of CpG islands in the gene promoter region was compared between patients with the CNA *vs*. patients with neutral copy number (log_2_ fold change; y-axis). Results were compared by tumour type and for known cancer driver genes *vs*. other genes (x-axis). Values greater than 0 indicate genes where methylation is higher in patients with altered copy number compared to patients with neutral copy number; values less than 0 indicate genes with less methylation in the CNA group. Values near 0 indicate little impact of CNA on methylation for those genes. The letters above the violins indicate pairwise differences between groups: two groups sharing the same letter do not significantly differ (Wilcoxon rank sum test q > 0.05) while two groups with different letters differ (q < 0.05). The letters should be interpreted separately for the Gain (top) and Loss (bottom) plots. **h-j.** The distributions of Spearman’s correlation for within-gene methylation, CNA and RNA in localized and metastatic samples. Gene methylation was calculated as the median β-value among CpG islands in the gene-promoter region.

Given that CNAs are common and early driver events in prostate cancer, we hypothesized copy number changes may interact with methylation to jointly regulate RNA abundance. We first verified that CNA status was positively associated with mRNA abundance in most genes (86%; **Figure 2d**). Next, we asked if the effects of CNAs on mRNA were modulated by the methylation status of the locus. We quantified these CNA-methylation effects on mRNA as the interaction term in linear regressions, focusing on the 725 tumours with all three data-types available. Almost half of all genes exhibited significant CNA-methylation interactions (4,950 of 10,919 = 45.3%; q < 0.05; **Figure 2e**). Thus, for about half the genome, fully understanding the gene-expression consequences of tumour CNA changes requires quantifying the methylation status of that genomic region. These CNA-methylation interactions were an even mix of synergy and antagonism, so in some cases methylation changes countered the effects of the CNA and in other cases they enhanced it. Six examples of notable interactions are given in **Figure 2f**, including the driver gene *PTEN*. In each case, the difference in RNA abundance between hyper-and hypo-methylated groups is much greater in patients with CNA gain or loss compared to patients with neutral copy number. The *PTEN* interaction was independent of both T-category and ISUP grade-group (**Supplementary Figure 5d,e**).

To determine if these CNA-methylation changes preferentially influence cancer driver genes, we considered 91 recurrently identified prostate cancer driver genes in the prior studies (8,44–46) (**Supplementary Table 5**). We stratified our analyses by CNA status and disease stage (localized *vs.* metastatic). We first examined 14 well-established recurrent driver mutation events (**Supplementary Figure 6**). In metastatic samples, methylation-CNA distributions were similar regardless of driver-event status, while localized samples typically show shifts in the center and spread of correlation distributions between patients with and without the driver mutation. We next evaluated the overall association between CNA and methylation across the 91 driver events. In localized disease methylation changes in driver and passenger genes were indistinguishable on average: driver genes showed similar amounts of differential methylation as non-driver genes: this was true for both losses and gains (**Figure 2g**). By contrast, in metastatic disease CNA changes were associated with much larger methylation changes overall, and these were significantly enhanced in driver genes (**Figure 2g**). This reflected a general pattern: increased copy number was strongly associated with reduced methylation in localized, but not metastatic disease (**Figure 2h, Supplementary Figure 7a**). Similarly, increased mRNA was strongly associated with reduced methylation in localized, but not metastatic disease (**Figure 2i, Supplementary Figure 7b**). By contrast, the association of copy-number and mRNA abundance was largely indistinguishable between localized and metastatic disease (**Figure 2j**; **Supplementary Figure 7c**). These data suggest methylation is more central to gene regulation in localized than in metastatic prostate cancer (**Supplementary Figure 7d**). To confirm this result, we assessed the mutual information between pairs of data types, which can capture nonlinear relationships. Consistent with our simple correlation analyses, mutual information showed substantial differences between localized and metastatic prostate cancer, particularly in methylation-RNA and methylation-CNA associations. By contrast, the influence of CNAs on RNA abundance was relatively consistent across both localized and metastatic disease (**Supplementary Figure 8**).

Given the inter-tumour and stage-dependent heterogeneity in CNA-methylation interactions, we hypothesized that genes with highly variable methylation status across tumours could identify loci with prognostic associations. Of the top 500 genes with <u>h</u>ighest <u>v</u>ariation in <u>m</u>ethylation (“HVM genes”), 30 were significantly associated with BCR (q < 0.10; **Supplementary Figure 9a**). The HVM genes showed greater variation in size and direction of their correlation with **<u>p</u>**ercentages of the **<u>g</u>**enome with a CN**<u>A</u>** (PGA) compared to non-HVM genes, with 30% of HVM genes being positively correlated with PGA, compared to only 9% of non-HVM genes (p = 4.5 x 10^-78^; **Supplementary Figure 9b**). A similar pattern was observed for within-gene methylation-mRNA correlation (p = 0.01; **Supplementary Figure 9c**). The HVM genes were not associated with different rates of CNA compared to non-HVM genes (p > 0.05) (**Supplementary Figure 9d**), and did not show enrichment based on chromosomal location (**Supplementary Figure 9e**), and were not enriched for any particular biological pathways. Rather, the 30 HVM genes that were significantly associated with BCR showed strong CNA-methylation correlations indicating that they reflect individual outlier events (**Supplementary Figure 9f-g**).

These data reveal a complex interplay between copy number changes, methylation and downstream gene-expression in prostate cancer. Copy number changes have stage-specific influences on methylation, allowing a single somatic mutation to influence gene-expression in different ways as the disease evolves. The unexpected inter-tumoural heterogeneity in the epigenomic consequences of copy number changes directly affects driver gene expression.

### Methylation subtypes reflect tumour genetics and microenvironments

The complex association between CNA, methylation and mRNA abundance led us to explore the genomic hallmarks of the methylation subtypes. We compared the CNA frequency for each CpG using ordinal logistic regression, adjusting for PGA and cohort. CNA frequency differed significantly across methylation subtypes for ∼20% of genes containing subtype-defining CpGs (**Figure 3a**). CNA gains were more prevalent in MS-3 and MS-4 across the genome, while MS-4 had a higher percent of samples with CNA losses in chromosomes 2, 5, 6 and 13 (**Figure 3b**; **Supplementary Figure 10**). The subtypes differed significantly in their overall CNA burden (Kruskal-Wallis p = 2.02 x 10^-55^; **Figure 3c**), chromosome-level CNA burden (**Supplementary Figure 11**), and driver CNA burden (**Supplementary Figure 12**). Tumours in MS-2 had the fewest CNAs, consistent with the observation that most normal samples (95%) had this subtype. Taken together, these data show that subtype-defining CpGs discriminate between tumour *vs.* normal tissue, localized *vs.* metastatic tissue, and have CNA profiles that differ across subtypes. MS-3 and MS-4 had significantly larger PGA, consistent with their higher risk of BCR.

**Figure 3:**
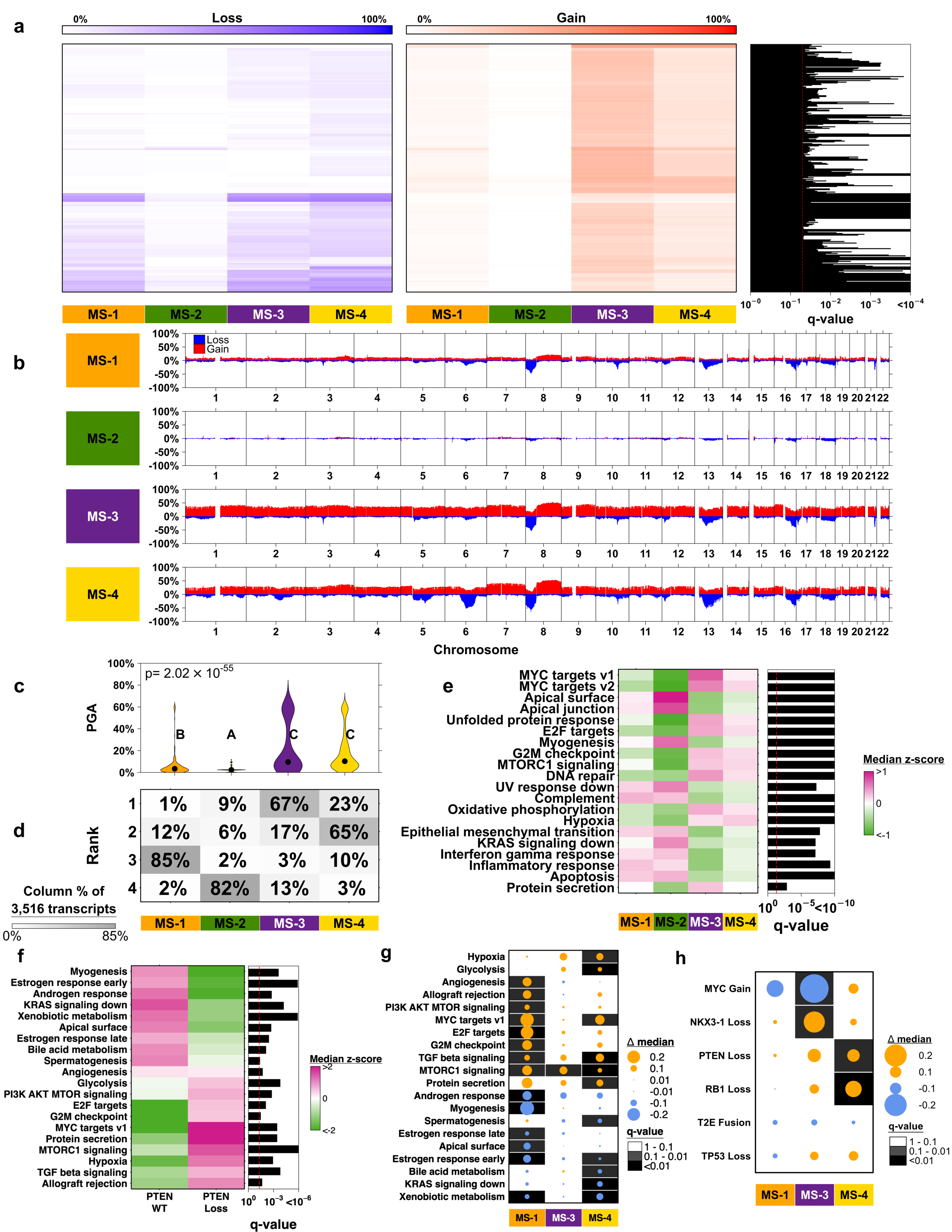
Methylation subtypes have different CNA and RNA profiles. **a.** CNA of 1653 genes containing the subtype-defining CpGs. Each gene (row of heatmap) was summarized by the percent of patients with copy number gain and loss within each subtype (columns). Rows were sorted by hierarchical clustering with Euclidean distance and complete linkage. Ordinal logistic regression was used to model a gene’s copy number status (loss, neutral, gain) as the dependent variable, with subtype, PGA and cohort as the independent variables. A likelihood ratio test p-value was calculated for subtype, and multiple testing-adjusted q-values are displayed in the right barplot. **b.** The y-axis indicates the percent of patients with copy number gain (red bar) or loss (blue bar) at that location of the genome (x-axis). Results are stratified by methylation subtype (row). **c.** Percent genome alteration (PGA) was compared between the methylation subtypes with Kruskal-Wallis rank sum test p-value. Groups sharing the same letter do not significantly differ (Mann-Whitney U-test q > 0.05), while groups with different letters have q < 0.05. **d.** For 3,516 mRNA transcripts (adjusted for PGA and cohort, see **Methods**) that significantly differed between subtypes (Kruskal Wallis rank-sum test q < 0.05 and standard deviation (SD) of subtype-medians > 0.4), the subtype medians were ranked from 1-4 (smallest to largest). Column-wise percentages show the percent of time a given subtype had each rank. **e.** Patient-specific pathway scores were calculated for 50 hallmark pathways using GSVA adjusted for PGA and cohort (Methods). For each pathway, the subtype median adjusted mRNA abundance z-score is displayed in the heatmap. Kruskal Wallis test q-values are shown in the right barplot, and pathways were sorted in descending order by the standard deviation of the subtype medians. The top 20 most significant pathways are shown. For the hypoxia response, the Buffa gene-set was used. **f.** For each pathway, the heatmap shows the median adjusted GSVA pathway z-score in *PTEN* wild-type (WT) and *PTEN*-loss tumours. The right-hand bar plot reports FDR-adjusted q-values from Wilcoxon rank-sum tests comparing *PTEN* WT *vs*. *PTEN* loss. Only pathways with q < 0.05 are shown. For the hypoxia pathway, the Buffa hypoxia gene-set was used. **g.** Associations between *PTEN* CNA status and GSVA pathway scores within each methylation subtype. Dots indicate effect sizes, defined as the median difference in GSVA pathway score between tumours with *vs*. without *PTEN* CNA loss within a subtype. Background colour indicates FDR-adjusted q-values. **h.** Associations between recurrent CNA driver events and GSVA hypoxia scores within each methylation subtype. Dots indicate effect sizes, defined as the median difference in hypoxia score between tumours with *vs*. without the alteration within a subtype. Background colour indicates FDR-adjusted q-values.

To determine if the genome-epigenome interactions were associated with downstream gene-expression, we next considered 738 prostate tumours with matched methylation and RNA profiling. When assessing subtype differences, 40,496 of 53,078 (76%) mRNA transcripts significantly differed between subtypes (q < 0.05; adjusting for PGA and cohort differences). We chose to focus on a subset of 3,516 significant transcripts with large effect sizes (**Supplementary Table 6**). These transcripts showed distinct expression profiles between methylation subtypes (**Supplementary Figure 13a**) with MS-1 and MS-2 tending to have higher abundance (**Figure 3d**). To contextualize these abundance differences, we next summarized hallmark pathway activity using GSVA. Because GSVA produces rank-based enrichment scores, we interpret these values as relative pathway activity across samples rather than absolute levels (47). At the pathway level, these differences reflected a broad range of cancer hallmark pathways (**Figure 3e**), with *MYC* targets showing the largest differences between subtypes (48,49). Given the significant CNA-methylation interaction observed in *PTEN*-deficient tumours (**Figure 2f**), we next assessed subtype-specific pathway activity within *PTEN*-loss cases. Angiogenesis pathway activity was elevated in MS-1, with no significant change in MS-3/4. In contrast, hypoxia and glycolysis scores were significantly increased in MS-4 (with a consistent, non-significant trend in MS-3) and were not significantly changed in MS-1 (**Figure 3f,g**).

Given the prominent subtype differences across pathways, we next focused on hypoxia, which is closely linked to prostate cancer progression and evolutionary dynamics and has well-established associations with prognosis and treatment response (43,50–52). We compared a validated hypoxia RNA signature (53) between the four methylation subtypes. MS-2 (the normal-like subtype) was significantly less hypoxic than the other subtypes (**Supplementary Figure 13b**). Subtype-defining CpGs had 1.6 times greater odds of being correlated with hypoxia compared to non-subtype-defining CpGs (95% CI 1.4-1.8, p = 1.48 x 10^-15^; **Supplementary Figure 13c**). CpGs throughout the genome were correlated with hypoxia, mostly inversely (69%; q < 0.05; **Supplementary Figure 13d**). Methylation of CpG islands in gene promoters was significantly correlated with hypoxia score in 1,046 genes (**Supplementary Figure 13e; Supplementary Table 6**). Finally, we examined associations between hypoxia scores and recurrent driver alterations (present in ≥ 20% of tumour samples) within each subtype. In MS-3, tumours with *MYC* gain showed significantly lower hypoxia scores, whereas tumours with *NKX3-1* loss showed significantly higher scores. In MS-4, hypoxia scores were significantly elevated with *RB1* and *PTEN* loss (**Figure 3h**). Collectively, pathway activities differ markedly across methylation subtypes and the associations between recurrent driver alterations and hypoxia vary by subtype, suggestive of subtype-specific evolutionary contexts.

### Somatic driver mutations shape the prostate cancer epigenome

Prostate cancer is a C-class tumour (8,54), with few recurrent driver single nucleotide variants (SNVs), but many driver CNAs and <u>g</u>enomic rearrangements (GRs). We evaluated the association of global methylation with 28 recurrent driver aberrations (8), as well as the mutation burden of SNVs, CNAs and GRs in 870 prostate tumours (**Figure 4a**). Consistent with our gene-expression analyses, PGA had the largest effect on methylation levels (87.5% significant CpGs, q < 0.05), followed by global SNV burden, *MYC* copy number gain and GR burden (**Figure 4b**). These CpG changes were distributed across the genome and affected both CpG Island and non-CpG Island sites (**Supplementary Figures 14a,b**), and influenced downstream mRNA abundance (**Supplementary Figure 14c**).

**Figure 4:**
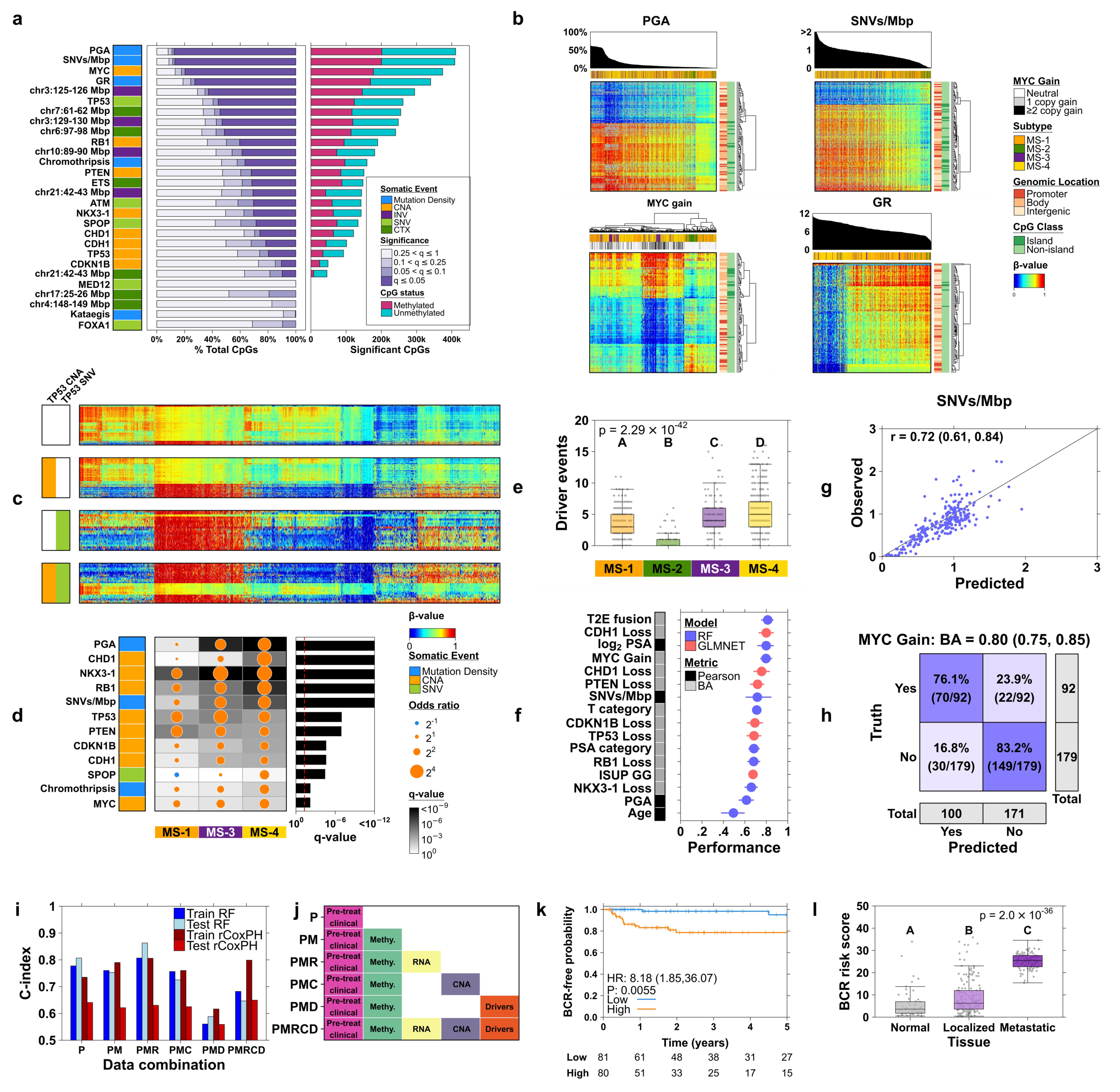
Somatic driver mutations impact the prostate cancer epigenome; and epigenetic signatures of clinico-molecular features. **a.** CpG β-values were tested for an association with 28 somatic events and clinical features. Somatic features were divided into five categories: mutational burden (mutation density), CNAs, INVs, SNVs, and CTXs including ETS fusions. Barplot on the left shows different q-value category percentages, resulting from Spearman’s ρ and Kruskal-Wallis tests for continuous and discrete features, respectively. Barplot on the right shows the number of significantly differentially methylated CpGs associated with each feature (q < 0.05), with methylated *vs.* unmethylated status (median β-value threshold = 0.61). **b.** Methylation profiles of the top 4 driver events with the most number of significant CpG associations. For each driver event, 100 CpGs are shown where q < 0.05, correlation > 0.6 for continuous features and epsilon-squared (ε²) > 0.2 for binary *MYC* Gain, and then select the CpGs with the 100 largest standard deviations. Rows (CpGs) are sorted using hierarchical clustering, complete linkage and Euclidean distance. Columns/samples for continuous features are sorted in descending order (PGA, SNVs/Mbp, GR count) while columns for *MYC* gain were sorted using the aforementioned hierarchical clustering. **c.** Heatmap of CpG methylation (columns) of patients (rows) stratified by *TP53* CNA loss and SNV mutation status (left covariate bars). The top 1000 CpGs that were significantly positively associated with *TP53* CNA loss (q < 0.05) and top 1000 inversely associated were combined with the top 1000 positively associated with *TP53* SNV, and top 1000 inversely associated. Columns and rows were sorted using hierarchical clustering with complete linkage and Euclidean distance. **d.** Logistic regression was used to test the association between driver events and subtypes while adjusting for cohort and PGA. The odds of having a given mutation (row) are shown for each subtype (column) when treating MS-2 as the reference group. PGA and SNVs/Mbp were median-dichotomized; 12 drivers with q < 0.05 are shown. **e.** Boxplots of the total number of driver events by methylation subtype, with Kruskal-Wallis p-value. Pairwise comparisons between groups were conducted using the Mann-Whitney U-test, such that groups with different letters significantly differ (q < 0.05). **f.** Performance of the final selected models for predicting features (y-axis labels). Dots indicate the estimated performance on the test dataset with adjacent lines giving 95% bootstrap confidence intervals. Pearson’s correlation (r) and balanced accuracy (BA) were used for numeric and categorical features, respectively. **g.** Test dataset performance for predicting log_2_ SNVs/Mbp. Pearson’s r is reported with 95% bootstrap confidence interval in parentheses. **h.** Test dataset confusion matrix for predicting *MYC* copy number gain. Balanced accuracy (BA) is reported with 95% bootstrap confidence interval in parentheses. Each cell is shaded by the percentage of the row total. **i.** A random forest (RF, blue) and regularized Cox proportional hazards model (rCoxPH, red) were trained for predicting time-until-BCR using different combinations of data as predictors (x-axis labels; see (**j**) for definitions). The height of the bars indicates the estimated C-index in the training and test datasets. Higher C-index indicates better performance. **j.** The combinations of predictors used in figure (**i**). For a given row, the y-axis label names the data combination and the boxes indicate which data types are included. **k.** The best model from (**i**) was used to predict BCR risk score in the test dataset of 161 patients and median-dichotomized into High *vs.* Low risk. The Kaplan-Meier plot shows the BCR-free probabilities for the High and Low risk groups, with hazard ratio, 95% confidence interval, and Wald-test p-value estimated from a Cox proportional hazards model. The table below the plot gives the number of patients at risk from 0-5 years. **l.** The best model from (**i**) was used to predict the BCR risk score for 52 normal samples, 161 localized prostate cancer samples from the test dataset, and 99 metastatic samples. The risk score was compared between the three tissue groups using Kruskal-Wallis rank sum test p-value. Groups with different letters have significantly different risk score profiles (pairwise Wilcoxon test q < 0.05).

To determine if different somatic mutations on a single gene induce similar methylation effects, we focused on the tumour suppressor *TP53*. We compared *TP53* inactivation across three groups: SNV only (6% of patients), copy number loss only (24%) and both SNV and copy number loss (8%). The *TP53* SNV-only category was associated with substantially larger methylome changes than *TP53* copy number loss (56% of CpGs *vs.* 20%; **Figure 4c**). The specific CpGs affected only partially overlapped (**Supplementary Figure 14d**), indicating that different mutation types induce different methylation changes.

Consistent with these results, the methylation subtypes were significantly associated with 12 of 28 driver mutations (q < 0.05, logistic regression adjusting for cohort and PGA), with MS-3 and MS-4 having greater odds of each mutation relative to the normal subtype MS-2 (**Figure 4d**; **Supplementary Figure 14e,f**). Similarly, MS-4 had more total driver events and MS-2 fewer (**Figure 4e**; **Supplementary Figure 14g; Supplementary Table 7**), even after controlling for the large differences in mutation burden and driver mutation frequency between localized and metastatic cancers (**Supplementary Figure 15**).

It is relatively uncommon to have sufficient resources and tissue to simultaneously profile tumour DNA and methylation. Given the strong associations between methylation and the mutational composition of a cancer, we attempted to create prediction models that infer features of a cancer’s somatic genome from its epigenome. We randomly split 1,436 patients into 70% for training (n = 1,005) and 30% for testing (n = 431). We focused on 15 important clinico-molecular features with sufficient data for model training: patient age and serum PSA (both continuous and discretized), tumour grade and T-category, mutation burden (SNVs and CNAs), seven tumour suppressors and one oncogene. We fit random forest and glmnet models in the training dataset, with standard feature selection, hyper-parameter optimization and assessment of over-fitting (**Methods**; **Supplementary Figures 16,17; Supplementary Table 7**).

The final models performed well on the final held-out test dataset, with all except age achieving performances > 0.60 (**Figure 4f**; **Supplementary Figure 18**). For example, tumour SNV mutational burden was very well-predicted by our methylation model, with a Pearson’s correlation of 0.72 between our predictions and actual DNA sequencing (**Figure 4g**). Similarly, our *MYC* gain prediction model accurately predicted 83% of MYC neutral cases and 76% of *MYC* gain cases despite only 34% of cancers having a *MYC* gain (**Figure 4h**). Only one model showed cohort-specific bias: PSA, reflecting the strong clinical bias between localized and metastatic disease (**Supplementary Figure 19**). All prediction models are available in the PrCaMethy R package (**Supplementary Figure 20a**) and we define a subset of ∼7,000 CpGs that overlap across prediction models for targeted assays (**Supplementary Figure 20b**).

### A multi-omic signature predicts disease recurrence

To extend these clinico-molecular methylation signatures to patient risk-stratification, we focused on 774 localized prostate cancer patients with time to BCR and multi-omic data available. We again split these randomly into 70% for training and 30% for testing, and employed standard model- and feature-selection (**Methods**; **Supplementary Figure 21a,b**). The optimal model in the training dataset was carried forward to the validation dataset, and included 2 clinical features (PSA, ISUP GG), methylation subtype, 10 promoter-methylation events and RNA abundance of 7 genes (**Supplementary Figure 21c**). Risk scores from this model predicted outcome in the test dataset with a C-index of 0.86 (95% CI 0.77-0.91), compared with 0.81 for a clinical-only model (95% CI for the difference: 0.0011 to 0.11) (**Supplementary Figure 21d**). Median-dichotomized risk-scores identified a group of patients with 8.2-fold greater risk of BCR in the held-out test dataset (95% CI 1.9-36.1; p = 0.0055; **Figure 4k**). Model risk scores differed significantly between non-malignant, localized and metastatic prostate samples (**Figure 4l**), and remained prognostic even within the subset of held-out testing patients classified as intermediate risk (HR = 16.23, p = 0.013; **Supplementary Figure 21e**). Overall, these results suggest the prostate cancer epigenome and its interactions with the transcriptome may serve as very accurate predictors of clinical outcomes.

### Inverse associations between DNA methylation age and driver mutations

The single clinico-molecular feature we were unable to accurately model from tumour methylation was patient age (**Figure 4f**). This suggested to us that prostate tumour epigenetic age might differ significantly from patient chronological age. We estimated the DNA methylation (DNAm) age of 102 normal samples using ten widely available methylation clocks (55). The Horvath clock achieved the highest correlation (Pearson’s r = 0.48, 95% CI 0.31 – 0.61, q-value = 5.18 x 10^-6^; **Figure 5a-b**), consistent with the original study of that clock (56,57). We then analyzed paired normal and tumour samples using the Horvath clock. Tumour samples exhibited decelerated DNAm ages compared to their normal counterparts, suggesting that the aging pathways in tumours differ from those in adjacent normal tissues (**Figure 5c**).

**Figure 5:**
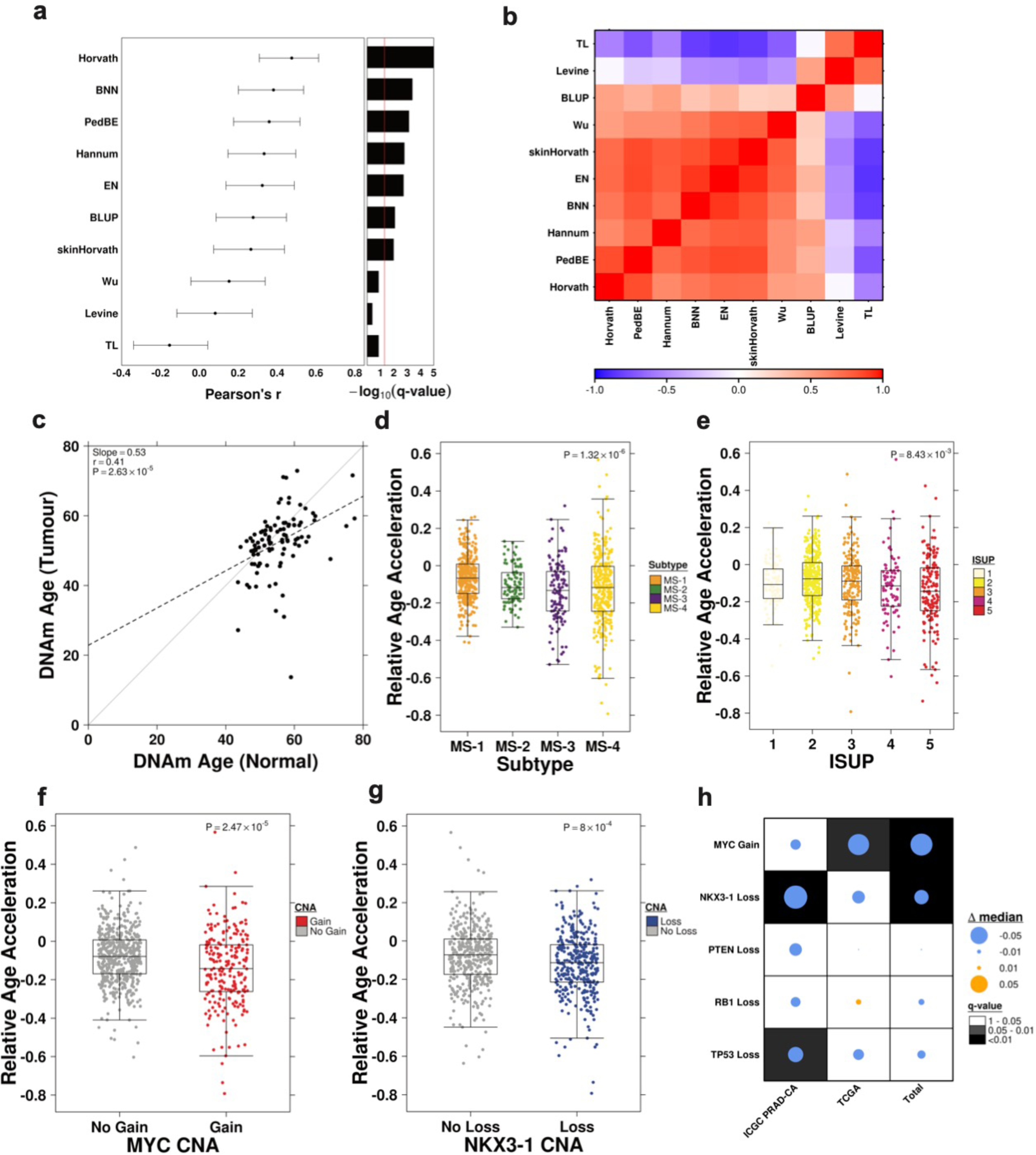
Somatic driver mutations influence prostate cancer aging. **a.** Performance of the ten DNA methylation clocks. Pearson’s correlation coefficient (r) is reported with a 95% confidence interval. False discovery rate (FDR)-adjusted q-values are shown in the right bar plot. **b.** Pairwise correlations (Pearson’s r) among the ten DNAm clocks. **c.** Pairwise comparison of DNAm age between normal and tumour samples. The dotted line represents the fitted linear model (slope = 0.53, intercept = 23.1) **d, e.** Relative age acceleration was compared among methylation subtypes (**d**) and ISUP groups (**e**) using the Kruskal-Wallis rank sum test on residuals adjusted for chronological age. **f, g.** Relative age acceleration was compared based on *MYC* CNA gain (**f**) and *NKX3-1* CNA loss (**g**) using the Mann-Whitney U test on residuals adjusted for chronological age. **h.** Summary of associations between CNA drivers and relative age acceleration (adjusting for chronological age) in ICGC PRAD-CA, TCGA, and the combined cohorts. Dots represent effect sizes, with background colour indicating q-values.

Next, we investigated the associations between clinico-molecular features and epigenetic age acceleration, adjusting for chronological age (**Methods**). Both the methylation subtypes (p = 1.32 x 10^-6^) and ISUP grade (p = 8.43 x 10^-3^) demonstrated significant associations with relative age acceleration (**Figure 5d-e**). Global mutation burden showed an inverse correlation with relative DNAm acceleration (SNVs/Mbp: Spearman’s ρ = -0.30, p = 2.82 x 10^-16^; PGA: Spearman’s ρ = -0.26, p = 2.53 x 10^-13^), mirroring the previously observed association between SNV burden and DNAm age in other cancers (56,57). Two specific drivers were associated with significant DNAm age deceleration: *MYC* copy number gain (p = 2.47 x 10^-5^) and *NKX3-1* loss (p = 8.00 x 10^-4^; **Figure 5f-h**). Taken together, these data show that prostate tumours exhibit the hallmarks of reduced age, and this is associated with clinical phenotypes, global mutation burden and with specific somatic driver mutations.

## Discussion

Cancer is a disease of continual genomic evolution. The epigenome serves as the critical mediator between somatic mutations and their gene-expression consequences, which ultimately drive clinical phenotypes like diagnosis, prognosis and response to therapy. A range of evidence has shown that various forms of epigenomic (dys)regulation, such as chromatin modifications, can drive prostate cancer phenotypes (58–60). Here we focus on DNA methylation to maximize sample-size and infer general principles of how it changes and influences phenotypes across the course of disease evolution.

We show that prostate cancers fit into four general methylation subtypes. Unexpectedly, each subtype can be identified at all stages from normal prostate through to metastatic disease, although at different frequencies (**Figure 6**). This suggests that methylation subtypes tend to evolve over time, but at least in part reflect the early tumour-initiation state. Consistent with that, the subtypes are associated with CIMP, specific somatic-mutational, transcriptional and clinical profiles.

**Figure 6:**
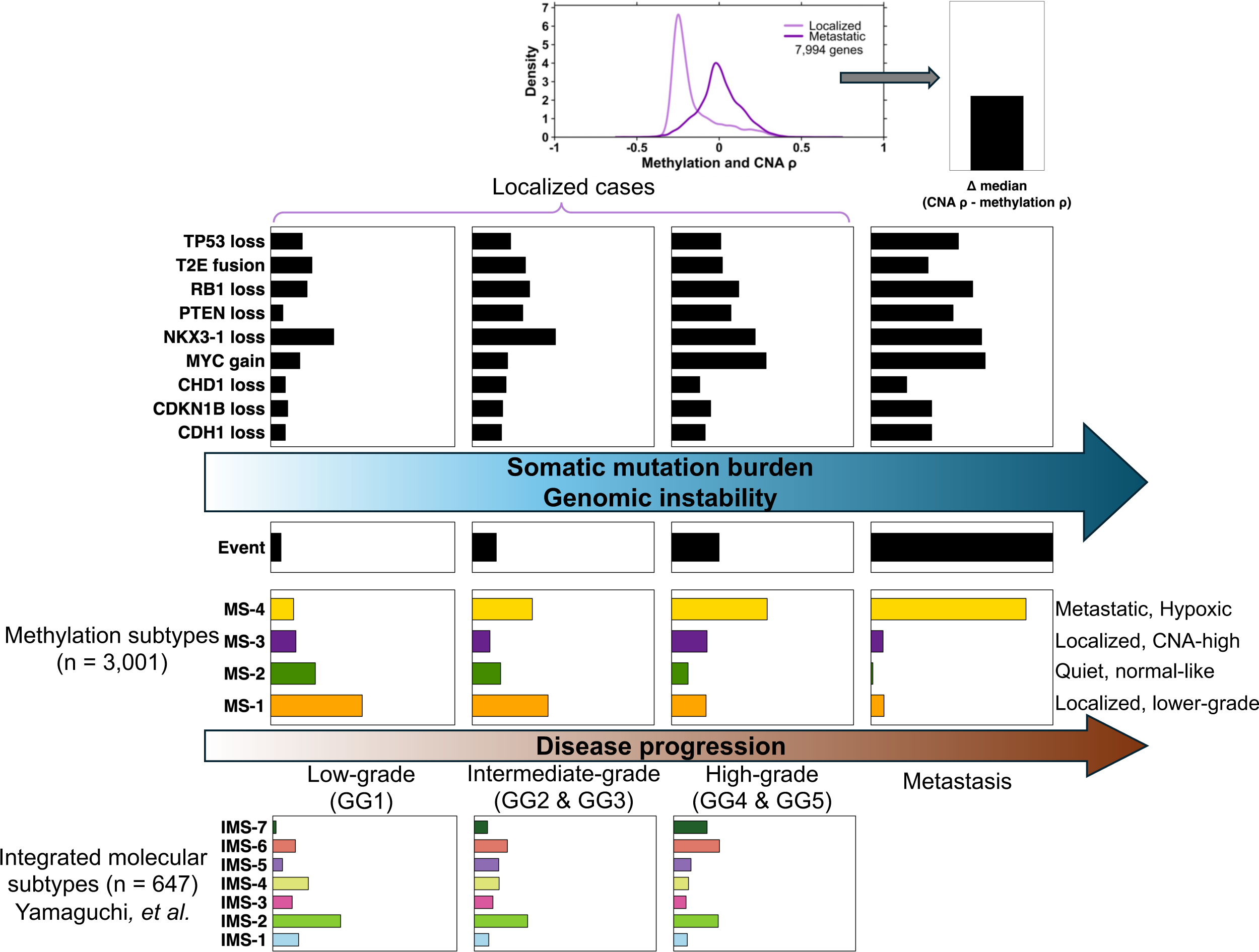
Methylation subtype distributions differ across disease states. Schematic overview of methylation subtype, clinical event, and recurrent driver event distributions across disease progression. Primary tumours were grouped as low-grade (GG1), intermediate-grade (GG2-GG3), and high-grade (GG4-GG5) according to ISUP GG, with metastatic tumours shown separately. For each disease state, proportions of MS subtypes, 5-year BCR rate, and recurrent driver events present in at least 10% of tumours are shown. Metastatic tumours are shown as uniformly event-positive. The overall MS subtype distributions for normal, localized, and metastatic cases are shown in Figure 1e. For comparison, proportions of integrated molecular subtypes (IMS-1 to IMS-7) from our previous study are shown at the bottom for the primary disease states. The upper inset summarizes the shift in DNA methylation-CNA association between metastatic and localized tumours, illustrated as Δ median ρ (metastatic − localized; ∼ 0.22).

Although WGS enables inference of tumour subclonal structure (10,16), extending methylation-based subtyping to the subclone level remains challenging. Bulk methylation profiles derived from Illumina 450K/850K arrays represent mixtures of epigenetic states, and while whole genome bisulfite sequencing (WGBS) provides read-level information, its typical coverage in tumour samples limits reliable resolution of low-frequency subclones. Accordingly, understanding the single-cell heterogeneity of the prostate cancer methylome (and of its subtypes) is a major opportunity to further elucidate how DNA methylation shapes cancer evolution.

The best-known drivers of prostate tumour evolution are changes in CNA, which are closely linked to methylation changes (8,13,61). We demonstrate that these changes vary significantly across individuals and across disease stages. The *PTEN* tumour suppressor serves as an excellent example of this, where both hypermethylation and hemizygous deletion can (but do not always) result in decreased gene expression, with characteristic DNA methylation profiles (61) that may lead to functional changes from *PTEN* haploinsufficiency (62–64).

We identified 30 HVM genes that were significantly associated with BCR. These BCR-associated HVM genes did not show pathway enrichment, but did show strong CNA-methylation correlations, highlighting the complexity and heterogeneity of methylation influences on gene-regulation and clinical outcomes. Prior work in one cohort showed DNA methylation heterogeneity in prostate cancer can parallel copy-number evolution, and CNA-methylation relationships are highly context-dependent (13). Methylation at distal *cis*-regulatory elements can influence gene expression over long genomic distances, such that simple promoter-level CpG assignment may dilute coherent pathway-level signal (65,66). Amongst the BCR-associated HVM genes, several have known links to prostate cancer biology, including *HNF1B* (67,68), *LDHB* (69), and *MT1E* (70), suggesting this approach may be valuable in other cancer types.

Using GSVA to summarize hallmark pathway activity (rank-based enrichment, reflecting relative rather than absolute levels), we found that pathway activity differs markedly across methylation subtypes within *PTEN*-deficient tumours. Angiogenesis pathway activity was most prominent in MS-1, consistent with evidence that angiogenic programs can be induced early in prostate tumorigenesis and may be accentuated by *PI3K*/*AKT* signaling in *PTEN*-loss settings (71–73). In contrast, glycolysis activity was significantly elevated in MS-4, aligning with reports that glycolytic metabolism and lactate production are more pronounced in advanced prostate cancer and can be promoted by *PI3K*/*AKT* signaling (74,75).

We also observed MS subtype-specific associations between recurrent driver alterations and Buffa hypoxia-signature activity. Prior work from our group reported an association between *MYC* gain and higher hypoxia levels using a median-binarized hypoxia score (top half = +1, bottom half = -1 per gene, summed across genes) (50). However, in the rank-based framework, *MYC* gain was associated with lower relative hypoxia signaling within MS-3, suggesting a shift in the balance of transcriptional programs that is specific to this methylation context. Together, these results highlight subtype-specific interplay between recurrent driver alterations and pathway-signature activity, consistent with subtype-specific evolutionary and microenvironmental constraints.

We extend previous correlative studies (76) to define precise epigenomic signatures of key prostate cancer driver events, and demonstrate that these are highly predictable. These findings support the view that methylation captures stable tumour states that reflect lineage programs and are shaped by genomic disruption and microenvironmental pressures. As methylation profiling of circulating tumour DNA becomes increasingly feasible (77,78), these provide an avenue to develop non-invasive techniques to estimate both individual driver events and global mutational features longitudinally. Moreover, because driver-pathway coupling differs across methylation subtypes, integrating methylation profiling with predictions of clinico-molecular features may support subtype-aware therapeutic hypotheses and stratification strategies. Finally, we anticipate that the next stage of validation will be most informative in unfavourable intermediate-risk and high-risk localized prostate cancer rather than low-risk disease managed with active surveillance, where improved molecular stratification could support optimal treatment strategies and guide study design.

Our DNAm age analysis recapitulated the decelerated DNAm age (56,57) and identified associations with clinico-molecular features, including ISUP grade and recurrent driver events, suggesting that “younger” DNAm age is not purely stochastic. The Horvath clock was trained on non-malignant tissues, and its clock CpGs may be directly perturbed in malignant methylomes. Thus, the apparent clock reversal may reflect tumour-driven remodeling of these loci rather than literal rejuvenation. Mechanistically, this is consistent with impaired cellular differentiation (*NKX3-1* copy number loss), and/or induction of an embryonic stem cell-like state (*MYC* copy number gain) (48,79).

While this study provides novel insights into the prostate tumour methylome landscape, the findings should be interpreted in light of several limitations. First, two cohorts (Li et al. 2020; Zhao et al. 2020) were profiled by WGBS, whereas most others used Illumina methylation arrays (**Methods**; **Supplementary Table 3**). To mitigate platform-related effects, we harmonized data by restricting analyses to CpGs shared with the Illumina 450K array. Importantly, subtype-CIMP concordance using independently defined CIMP scores remained robust after adjustment for cohort and methylation-profiling platform.

Another limitation of our study is that raw IDAT files were not available for all cohorts, preventing uniform intensity-level reprocessing across datasets. The TCGA subtype discovery cohort used SeSAMe-processed data, which can reduce spurious methylation measurements (including artifacts associated with copy number loss regions). Other cohorts were processed using a range of pipelines. To mitigate this, we incorporated a measure of global CNA burden (i.e. PGA) into downstream models to reduce the influence of copy-number-related methylation bias on our findings. However, PGA is a global measure and does not replace locus-specific CNA-aware correction of array intensities.

Multiple methods are available for epigenetic subtype discovery, including discrete clustering, integrative multi-omic models (80,81), and admixture-aware deconvolution approaches (82,83). Here, we applied consensus clustering with a silhouette-based method because it yields reproducible and classifier-ready subtype labels that can be readily extended as new cohorts are added. This class of approach has two general limitations. First, it uses a single modality, whereas integrative subtype frameworks that combine multiple omic layers and molecular data types have been proposed. In practice, there is a trade-off between multi-omic richness and preserving sufficient sample size and cohort diversity to rigorously evaluate generalizability. Second, our analysis relies on bulk tumour profiles, which average across malignant and microenvironmental compartments and compress intratumoral heterogeneity. These considerations motivate future work using single-cell to explicitly model admixture and relate bulk subtype assignments to mixtures of malignant and microenvironmental cell states.

In addition to these technical limitations, our metastatic cohort is smaller and has limited treatment annotation, which constrains our ability to fully characterize the epigenetic subtypes that may emerge under therapeutic pressure. Larger metastatic datasets with detailed treatment histories, ideally sampled longitudinally, will be essential for defining therapy-induced epigenomic states more completely.

Because this compendium integrates cohorts with differing clinical composition and technical characteristics, tumour purity and genetic ancestry represented potential sources of confounding. To assess the robustness of our findings, we performed extensive sensitivity analyses and cohort-adjusted analyses (**Methods**; **Supplementary Information**). Our main conclusions remained unchanged, although further systematic benchmarking of purity estimation algorithms and harmonized genetic ancestry annotation across cohorts would be valuable.

Collectively, our data elucidates central roles for DNA methylation in regulating prostate cancer throughout its lifespan, from early aging deceleration through to extensive gene-regulatory effects in localized disease and strong biomarkers of clinically important events like relapse and metastasis. Despite the size of our compendium, it lacks coverage of untreated and lymph node metastases, and these may help fill a key missing piece in our understanding of the mechanisms of prostate cancer evolution from localized lesions into lethal metastatic ones.

## Materials and Methods

### Patient cohorts

The TCGA-PRAD cohort (n = 498 tumour samples) (7) was used to derive the methylation subtypes which were validated in 14 external prostate cancer cohorts (13 published and one unpublished - Melbourne) (7,8,13,15,25,30–38) (**Figure 1b-e**, **Supplementary Figure 2**, **Supplementary Tables 1-2**). TCGA, ICGC PRAD-CA and Zhao *et al.* 2020 prostate cancer cohorts were included in the methylation, mRNA, CNA, and somatic driver mutation analyses. See **Supplementary Figure 1a** for sample sizes per cohort and data type used in each analysis.

All International Cancer Genome Consortium (ICGC) samples were processed from prostate cancer tissue taken from patients with localized disease, following ethics approval from the University Health Network Research Ethics Board (REB)–approved study protocols (UHN 06-0822-CE, UHN 11-0024-CE, CHUQ 2012-913:H12-03-192) as previously described (8).

### Pathology and Tissue Processing

ICGC PRAD-CA fresh-frozen tissue specimens were audited for tumour content, Gleason/International Society of Urological Pathologists (ISUP) grade, and tumour cellularity by two independent genitourinary pathologists, as previously reported (8). Specimens of at least 70% tumour cellularity were manually macro-dissected and digested for nucleic acid extraction, as previously reported (8,9).

### Microarray data generation

ICGC PRAD-CA methylation, SNP and mRNA microarray data were generated as previously described (8). Briefly, Illumina Infinium HumanMethylation450 BeadChip kits were used to assess global methylation levels with 500 ng of input genomic DNA at the McGill University and Genome Quebec Innovation Centre (Montreal, QC). SNP microarrays were performed with 200 ng of DNA on Affymetrix OncoScan FFPE Express 2.0 and 3.0 arrays. Where DNA quantities were limiting (17 samples), we used whole-genome amplification (WGA; WGA2, Sigma-Aldrich, St. Louis, MO), and validated that WGA of genomic DNA does not significantly alter CNA profiles (9). Total RNA was extracted from alternating adjacent sections (to reduce potential heterogeneity relative to the DNA specimen), using the mirVana miRNA Isolation Kit (Life Technologies). Extracted total RNA (100-150 ng) was assayed on Affymetrix Human Gene 2.0 ST (HuGene 2.0 ST) array and Human Transcriptome Array 2.0 (HTA 2.0) platforms at The Centre for Applied Genomics (The Hospital for Sick Children, Ontario, Canada) and the London Regional Genomics Centre (Robarts Research Institute, London, Ontario, Canada).

### Methylation

Methylation processing was performed in R statistical environment (v3.4.0). Methylation data for the ICGC PRAD-CA cohort is in the Gene Expression Omnibus (GEO): GSE107298, and was processed using the Dasen method (84,85) as previously described (8). Probes that have detectability below background noise, non-CpG methylation, SNPs with an annotated minor allele frequency of greater than 5% with a maximum distance of two nucleotides to the nearest CpG site, and those known to cross-hybridize to multiple genomic locations were excluded. Probe average intensity levels were calculated across technical replicates. Probes were annotated to gene symbols *via* the IlluminaHumanMethylation450kanno.ilmn12.hg19 annotation package (v0.6.0). Unless stated otherwise, all methylation analyses were performed using the β-value, which represents the ratio of methylated intensity over combined methylated and unmethylated intensity and reflects the methylation level of a specific CpG locus. Tumour purities were estimated using LUMP as previously described (86). Purity estimates by LUMP were high overall (Q1 0.83; median 0.88; Q3 0.94) with no samples below 0.3, and no tumour purity thresholds were applied. Additional purity estimates were profiled using ten algorithms spanning four modalities for sensitivity analysis (86–93) (**Supplementary Information**).

Illumina Infinium Human Methylation 450K or 850K BeadChip CpG processed β-values were obtained for the following cohorts: Kirby *et al*. 2017 (31), Zhao *et al.* 2017 (25), TCGA-PRAD (7,94), ICGC PRAD-FR (32), Ramakrishnan *et al.* 2024 (34), Ylitalo *et al.* 2021 (36) and Mundbjerg *et al.* 2017 (38). For the subtype discovery TCGA-PRAD cohort (498 prostate tumours and 50 normal prostate samples), we used the processed DNA methylation dataset from GDC Data Release 34, generated using the SeSAMe pipeline (95). Additional details about data processing can be found in the corresponding citations. In all analyses, only CpGs from the Illumina 450K array were used.

For Melbourne methylation data, Illumina’s Infinium Human Methylation 450K array was used to profile the methylation status of all Australian prostate cancer samples. Raw signal data were processed and normalised for both within-array and between-array variations using the minfi package (96). Minfi’s implementation of the Illumina’s reference-based normalisation and the SWAN method within the minfi package were used for between-array and within-array (correcting for the probe-type bias) normalisations, respectively. Any methylation measurements with detection p-value greater than 0.01 were omitted from further analysis.

For cohorts Brocks *et al.* 2014 (13), Toth *et al.* 2019 (37) and Dairo *et al.* 2024 (35), raw Illumina methylation IDAT files were processed using the minfi R package (v1.52.1) (96). Methylation values with detection p-values greater than 0.05 were excluded. Probes with detection failure rates exceeding 10% across samples were also removed. Additionally, any sample in which more than 10% of probes failed detection was excluded from further analysis. Normalization was done using the dasen method in the wateRmelon R package (v2.12) (84,85). WGBS methylation data were used in Li *et al.* 2020 and Zhao *et al.* 2020 as detailed in (15,30). The “illumina450k_hg38” dataset from the Rockermeth R package (v.0.1.1) (97) was used to identify locations of the CpGs corresponding to the HumanMethylation450 BeadChip, thus allowing us to obtain the same CpGs used in the other cohorts. In Zhao *et al.* 2020, CpGs with a median number of reads < 10 and patient-level CpG calls with < 10 reads were omitted from analyses; while a cutoff of 5 was used in Li *et al.* 2020 due to lower coverage. For each CpG, the β-value was calculated as the proportion of methylated reads at that CpG site: # methylated reads / (# methylated reads + # unmethylated reads).

### Genetic Ancestry

Publicly available genetic ancestry group classifications were obtained for the ICGC PRAD-CA (16) and TCGA-PRAD (98) cohorts. For the Zhao *et al.* 2020 (15) cohort, we estimated genetic ancestry as follows. Categorical ancestry was inferred in relation to the super-continental ancestral populations of the 1000 Genomes (1KG) reference panel: European (EUR), African (AFR), East Asian (EAS), South Asian (SAS) and Admixed American (AMR) (99). First, genotypes were merged with the 1KG reference cohort (Phase 3 primary release, genome build GRCh38, founders only), and the set of intersecting SNPs was extracted using bcftools. Merged genotypes were converted to PLINK2 (100) format and filtered for a maximum missingness of 0.05 and a minimum allele frequency of 0.01, then pruned for variants in high linkage disequilibrium. Genotype principal components (PCs) were computed using the PLINK2 --pca flag. K-nearest-neighbors (KNN) was applied to the first five PCs to assign samples to 1KG-annotated ancestry clusters.

Continuous ancestry was defined as proportions of admixture from 1KG populations. The ADMIXTURE (101) tool was used in unsupervised mode to estimate admixture proportions in the merged, intersecting SNP-set for each sample. The hyper-parameter K, indicating the number of expected populations, was set to 5, mirroring the five 1KG super-populations. Inferred clusters were annotated with 1KG population labels based on which cluster contributed the greatest admixture proportion in each known reference sample set.

Five patients where KNN-PC based ancestry classifications disagreed with ADMIXTURE based classifications were considered “admixed” and omitted from further analyses involving genetic ancestry.

The associations between SIR and genetic ancestry with key clinico-molecular features were quantified using Cramér’s V and epsilon-squared (ε²) effect sizes for categorical and continuous features, respectively.

### Copy number alterations (CNAs)

ICGC PRAD-CA CNA data was obtained as previously described (8). The output from the Affymetrix OncoScan FFPE Express 3.0 SNP probe assay was aligned to a custom reference (8) using the OncoScan Console v1.1. Copy number segments were subsequently called using the BioDiscovery’s Nexus Express for OncoScan software using the SNP-FASST2 algorithm. Parameters were identical to the default one except for the minimum number of probes per segment which was set to 20. Diploid re-centering was performed when needed using the Nexus Express software. Using BEDTools the gene-level CNAs were identified with the RefGene annotation (2014-07-15) (102). Percent genome alteration was defined as PGA = (base-pair length of all genome regions with gain or loss) / 3.2 billion bases * 100. Zhao *et al.* 2020 CNA data was obtained from (103) and TCGA CNA data was downloaded from the TCGAbiolinks R package v2.25.3 (7,94) with copy number calls of 2 treated as “neutral”, less than 2 as “loss” and greater than 2 as “gain”.

### mRNA

ICGC PRAD-CA pre-processed mRNA data was downloaded from GSE107299 and processed in R (v3.4.3). Briefly, the RMA algorithm implemented in the affy (v1.48.0) package (104) was used with modified Entrez Gene CDFs to pre-process the raw intensity data. Probes were mapped to Entrez gene ID using custom CDF files (v18) for HTA 2.0 and HuGene 2.0 ST array (105). The sva package (v3.24.4) was used to correct for batch effects between different arrays (106). mRNA abundance levels from HuGene 2.0 ST and HTA 2.0 were combined based on Entrez Gene IDs. The mRNA abundance levels were averaged amongst duplicated Entrez Gene IDs. Entrez gene IDs were then converted into gene symbols and chromosome locations based on the human reference genome GRCh37 from UCSC table browser (download date: 2018/01/30). TCGA processed mRNA abundance levels were downloaded from the TCGAbiolinks R package v2.25.3 (7,94). Zhao *et al.* 2020 FASTQ files were aligned to GRCh38 with GENCODE v36 using STAR (v2.7.6a) (107) and mRNA abundance was calculated using RSEM (v1.3.3) (108). All mRNA abundance levels were analyzed as log_2_ transcripts per million (log_2_ TPM), unless specified otherwise.

### Whole-genome sequencing

Data processing and variant calling for the ICGC PRAD-CA cohort is described in (16). Chromothripsis scores were generated using ShatterProof (v0.14, default settings) (109). Kataegis was quantified using a fixed-width sliding window to test deviation of observed SNV trinucleotide content and inter-mutational distance from expected using an exact binomial test (110).

Corresponding whole-genome sequencing data for the Zhao *et al.* 2020 cohort (15,103) was downloaded as BAM files (See **Data Availability** Section). The BAM files were converted to FASTQs and realigned, and variants were called using Metapipeline-DNA (v5.0.0-rc.4 - 5.3.1) (111). A consensus SNV call set was created by retaining SNVs that were present in two or more callers using BCFtools (v1.17) (112). Consensus SNVs were then filtered using allow-and deny-lists. Deny-lists that were used included AccuSNP, Complete Genomics, dbSNP (v156) (113), duplicate gene (v68) (114), Encode DAC and Duke (115), gnomAD (v3.1.2, variants with ‘PASS’ flag) (116) and 1000 Genome project (99,117). Allow-lists involved Catalogue of Somatic Mutations in Cancer (COSMIC) (v92) (118) coding and non-coding variants. Somatic structural variants were called using Manta (v1.6.0) (119) and annotated using the StructuralVariantAnnotation R package (v1.18.0) (120).

### General statistical considerations

The Benjamini-Hochberg method was used to calculate multiple testing adjusted p-values to control the false discovery rate (FDR) at level 0.05 (referred to as ‘q’ or ‘q-value’ throughout) (121). When using Fisher’s Exact Test on larger than 2x2 contingency tables, the stats R package fisher.test function was used with 100,000 simulation replicates.

### Methylation subtypes

Methylation subtypes were identified in the TCGA-PRAD cohort (n = 498) using Interpretable MiniPatch Adaptive Consensus Clustering (IMPACC v0.1.0) (122) with default settings, 10,000 clustering partitions and CpG β-values. CpGs were filtered to keep the top 50,000 highest variance CpGs for clustering. The silhouette method (123) was used to choose the number of subtypes that maximized the median silhouette score, resulting in 4 subtypes. Specifically, the total number of subtypes was varied from 2 to 6, and each time the median silhouette score was calculated for each cluster, with the median of the cluster-wise scores used as an overall measure of clustering accuracy. A nearest shrunken centroid classifier (124) was trained on the TCGA subtypes to classify samples from seven cohorts, with missing values imputed by k-nearest neighbors imputation (impute R package v1.74.1, default settings) (125). As additional cohorts were incorporated into the project, several were profiled using the Illumina 850K methylation array, which lacks approximately 9% of the CpG sites present on the Illumina 450K array used to derive the subtypes. To address the resulting missing data, a random forest classifier (randomForestSRC R package v3.3.3) was trained on the TCGA subtypes. This method was chosen for its built-in imputation capabilities, enabling robust classification of samples from the remaining seven cohorts despite incomplete CpG coverage (126).

### Subtypes *vs.* clinical features

Categorical pre-treatment clinical features were summarized in **Supplementary Table 1** using counts and percentages. Race was self-reported, except for Chinese patients from Li *et al.* 2020 (30), whom we classified as Asian. The subtypes were tested for association with pre-treatment clinical features of age, T-category, ISUP GG and PSA. Each of the clinical variables was categorized as in **Supplementary Table 1** and tested for an association with the methylation subtypes using Fisher’s Exact Test and the bias-corrected Cramér’s V effect size (127). The subtypes were then tested for associations with the following outcomes: time until biochemical recurrence (logrank test; TCGA, ICGC PRAD-CA, Kirby *et al*. 2017, ICGC PRAD-FR), time until death (logrank test; Zhao *et al*. 2020) and binary outcomes of metastasis or death during follow-up (Zhao *et al.* 2017), BCR within 5 years (Dairo *et al.* 2024) and BCR within 3 years (Toth *et al.* 2019). Fisher’s exact test was used for binary outcomes. A permutation logrank test was used in cohorts Kirby *et al*. 2017 and Zhao *et al*. 2020, due to small and uneven group sample sizes (10,000 permutations) (128). Furthermore, the association between subtypes and BCR was assessed when pooling all patients with available BCR data (n = 882) where hazard ratios with 95% confidence intervals and Wald-test p-values were estimated using a Cox proportional hazards model treating MS-1 as the reference group, and adjusting for cohort differences. BCR was defined as two consecutive post radical prostatectomy (RP) PSA measurements of > 0.2 ng/mL, within 6 months. Patients who underwent successful salvage radiotherapy following RP (*i.e.* post-RP PSA below 0.2 ng/mL) were classified as non-failures; those that underwent unsuccessful salvage radiotherapy (*i.e.* two post-RP PSA readings above 0.2 ng/mL) were classified as failures, backdated to the original date of post-RP PSA rise. Patients who underwent other salvage treatments (*e.g.* hormones, chemotherapy) to control rising PSA were considered as having BCR.

Lastly, meta-analysis was performed for seven cohorts of localized disease with BCR-related outcomes. Four cohorts had right-censored time until BCR (TCGA, ICGC PRAD-CA, Kirby *et al.* 2017, ICGC PRAD-FR) and three cohorts had binary BCR-related outcomes: BCR within 5 years (Dairo *et al.* 2024), BCR within 3 years (Toth *et al.* 2019) and metastasis or death during follow-up (Zhao *et al.* 2017). A Cox proportional hazards model was used to estimate hazard ratios for time until BCR comparing subtype MS-4 to MS-1. For binary outcomes, the relative risk of MS-4 versus MS-1 was converted to a log hazard ratio and standard error using the approach of Watkins and Bennett (129). All log hazard ratios and standard errors were input to a random effects meta analysis using the metagen() function from the meta R package v8.0.2 with the DerSimonian-Laird estimator of between-study variance (130,131). The heterogeneity Q-test was used to assess heterogeneity in effect sizes between cohorts.

### CIMP classification

CpG Island Methylator Phenotype (CIMP) classification for TCGA-PRAD samples was defined and obtained from Sánchez-Vega *et al.* (23). Briefly, CIMP subtypes were derived by *k*-means clustering based on the average β-values of 450 CpG methylation probes provided in Sánchez-Vega *et al.* (23) for TCGA-PRAD samples. A clustering parameter of *k* = 3 was used to define three subtypes: CIMP-, CIMP intermediate, and CIMP+. The correspondence between CIMP classification and MS groups was evaluated using the Adjusted Rand Index (ARI) and the Qannari test. CIMP scores were computed for all samples as the average β-value across the same 450 CpG methylation probes. To account for differences in methylation technology and cohort, linear regression was used to regress CIMP scores on technology platform and cohort, and the resulting residuals were used for downstream comparisons with MS subtypes and other features. CIMP scores were compared across MS subtypes and other key clinical or molecular features using the Kruskal–Wallis rank-sum test and the epsilon-squared (ε²) effect size (132), followed by pairwise Mann–Whitney U tests for within-category comparisons. CIMP-low and CIMP-high groups were defined as samples within the 0 - 33rd and 66 - 100th percentiles of CIMP scores, respectively. Each feature was tested for an association between CIMP-low and CIMP-high groups using the χ² test or Fisher’s exact test when any cell counts were < 5, along with Cramér’s V (127). Driver mutations present in ≥20% of tumour samples were analyzed in relation to CIMP scores using the Mann-Whitney U test. FDR-adjusted q-values were computed.

### Subtype-defining CpGs

IMPACC intrinsically performs variable selection and identified 5,486 CpGs that were important for deriving the subtypes (referred to as ‘subtype-defining CpGs’ throughout). The subtype-defining CpGs were compared between subtypes using Kruskal-Wallis rank sum test and the epsilon-squared (ε²) effect size (132), while further compared between tumour and normal samples, and localized *vs.* metastatic tumour samples using the Mann-Whitney U-test. Chromosome position, CpG type (island, open sea, shelf, shore: https://www.illumina.com/content/dam/illumina-marketing/documents/products/other/field_guide_methylation.pdf), and gene region (gene body, intergenic, or gene promoter) were compared between the subtype-defining CpGs and non-subtype-defining CpGs using Fisher’s Exact Test and odds ratio. The gene regions were defined as follows: CpGs were annotated using the IlluminaHumanMethylation450kanno.ilmn12.hg19 bioconductor R package (v0.6.0) where CpGs in gene promoter regions were defined as within 200 or 1500 base pairs from a transcription start site (TSS). CpGs that were not mapped to any gene symbol were defined as “intergenic regions”, and remaining CpGs were defined as occurring within the gene body.

### Methylation interaction with mRNA and CNAs

Adjusted mRNA abundance was calculated by fitting a linear regression model to each mRNA transcript (log_2_ TPM) with cohort (TCGA, ICGC PRAD-CA, Zhao *et al.* 2020) and PGA as predictors, and using the standardized residuals (z-scores) in subsequent analyses. The adjusted mRNA abundance was compared between methylation subtypes using Kruskal-Wallis rank-sum test. Pathway analysis was performed using 50 hallmark pathways (49) by first calculating patient-specific pathway scores using adjusted mRNA abundance with the GSVA R package (v.1.50.0, default settings) (47) and comparing the pathway scores between methylation subtypes using Kruskal-Wallis rank-sum test.

An mRNA signature of hypoxia was calculated separately for the TCGA and ICGC PRAD-CA cohorts using a panel of 51 genes (53). The GSVA based hypoxia score was then z-score transformed (separately by cohort), and compared between methylation subtypes using Kruskal Wallis rank sum test and pairwise Mann-Whitney U-tests. Spearman’s correlation was used to assess the association between each CpG and hypoxia score, with Fisher’s exact test and odds ratio to assess whether subtype-defining CpGs were more likely to be correlated with hypoxia compared to non-subtype-defining CpGs. Lastly, Spearman’s correlation was used to assess the association of hypoxia score with gene-level methylation, where the latter was calculated for each patient as the median β-value of CpG islands in the gene promoter region.

To test the association of prostate tumour methylation levels with CNAs and mRNA abundance level, we first examined copy number effect on CpG methylation β-values. For each patient, we generated segmented CNA profiles and called each segment as: homozygous deletion, hemizygous deletion, neutral, gain or high gain. The β-values for each CpG from all patients within any given segmented CNA state were then used to examine CNA effect on methylation levels.

Spearman’s correlation was used to test the within*-*gene associations of CNA and methylation of CpG island in gene promoters (CGIp) and CGIp methylation with mRNA abundance levels, when pooling TCGA, ICGC PRAD-CA and Zhao *et al.* 2020 cohorts. Furthermore, linear regression of z-score transformed log_2_ TPM was used to test the interaction between CNA and methylation. This model considered binary methylation (hyper *vs.* hypomethylation; median dichotomized), binary CNA status (loss *vs.* neutral if loss was the more common mutation, otherwise gain *vs.* neutral), adjusted for cohort (TCGA, ICGC PRAD-CA, Zhao *et al.* 2020) as a covariate, and removed genes with < 5 CNA events in either methylation group.

The CNA status for genes containing the subtype-defining CpGs was compared between the four methylation subtypes using ordinal logistic regression from the polr function in the MASS R Package (133). A given gene’s copy number status (loss, neutral, gain) was used as the dependent variable, with subtype, PGA and cohort as the independent variables, and a likelihood ratio test was calculated for subtype. Percent genome alteration was defined as PGA = (length of all genome regions with gain or loss) / 3.2 billion bases * 100 and was compared between the four subtypes using Kruskal-Wallis rank sum test and pairwise Mann-Whitney U-tests.

The top 500 genes with highest variation in methylation (“HVM genes”) were tested for associations with time-until-BCR using a Cox proportional hazards model Wald test. Spearman’s correlation was used to test the association between each HVM gene with PGA and compared with the distribution of PGA correlations for non-HVM genes. The percent of patients with CNA loss or gain was calculated separately for HVM and non-HVM genes and compared using Mann-Whitney U-test.

Lastly, we investigated whether the associations between methylation, CNA and RNA within each gene differed between localized and metastatic prostate cancers. Two approaches were taken: 1) visually comparing the distribution of Spearman’s correlations for each pair of data types, and 2) comparing the distribution of mutual information for each pair of data types. Unlike Spearman or Pearson’s correlation, mutual information can capture arbitrary linear or nonlinear relationships. The mutual information was calculated as follows: for each pairwise data type comparison, data from each gene were discretized into N^1/3^ bins, where N is the intersection of the samples with data available for each pair of datatypes (134). Mutual information (MI) between two data types was defined as:

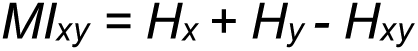

where *H_x_* and *H_y_* are the marginal entropies of data types *x* and *y* and *H_xy_* is the joint entropy, calculated using the R package Entropy (v1.2.1) (135). To reduce bias in MI estimates, the James-Stein shrinkage estimator (136) was applied, as implemented via the entropy.shrinkage function of the entropy R package (135). MI was normalized over the mean entropy of the two input vectors. To assess statistical significance of normalized MI values, permutation testing was performed. Gene-level data were permuted 1000 times to generate a null MI distribution, and p-values were calculated as the proportion of null MI values greater than or equal to the true observed MI. Multiple testing adjusted q < 0.05 was regarded as statistically significant.

### Driver mutations

Within the TCGA, ICGC PRAD-CA and Zhao *et al.* 2020 cohorts, CpG methylation levels were compared with each of 28 previously established somatic events (8). Spearman’s correlation test was used for continuous events and Kruskal-Wallis rank-sum test for discrete events. Somatic features were divided into six categories: mutational burden (chromothripsis, kataegis, number of GRs, number of SNVs and PGA). SNVs, CNAs (losses of *CDH1*, *CDKN1B*, *CHD1*, *NKX3-1*, *PTEN*, *RB1, TP53* and gain of *MYC)*, *ETS* fusion, CTXs (translocations with break-ends within chr4:148, chr6:97, chr7:61, chr17:25 and chr21:42 Mbp blocks) and INVs (inversions with break-ends within chr3:125, chr3:129, chr10:89 and chr21:42 Mbp blocks). CpG β-values were tested for an association with the 28 somatic events and clinical features, using Spearman’s ρ and Kruskal-Wallis tests for continuous and discrete features, respectively. Driver events that were significantly associated with at least one CpG (q < 0.05), were summed to calculate the total number of significant driver events per patient, and then compared between subtypes using the Kruskal-Wallis test and pairwise Mann-Whitney U-tests.

Lastly, the distribution of each driver mutation was compared between localized and metastatic prostate cancers. Mann-Whitney U-test p-value was calculated for continuous drivers. Fisher’s exact test q-value and odds ratio were calculated for categorical drivers, as well as logistic regression odds ratio adjusting for total mutation burden (CNA drivers adjusted for PGA, SNVs adjusted for log_2_ SNVs/Mbp, and genomic rearrangements adjusted for log_2_ GR count).

### Epigenetic predictors of clinical features and driver mutations

A total of 1,436 patients were randomly split into a 70% train (n = 1,005) and 30% test (n = 431) dataset. Two machine learning models were compared for predicting clinical features and driver mutations: random forest and a regularized general linear model (glmnet) (137,138). These two models were considered because both random forests and regularized linear models have been shown to perform well across a wide variety of applications (139,140). Random forest can automatically capture potential interactions or non-linear effects (141), but is harder to interpret compared to glmnet. In contrast, glmnet assumes linear and additive effects (no interactions) by default, and has a similar interpretation as traditional logistic or multinomial classification models (estimates log odds ratios for each predictor) but performs automatic feature selection by shrinking coefficients towards zero, attempting to remove unimportant features. Random forest was fit using the randomForest R package (v4.7.1.1) with the mtry parameter tuned by the caret R package (v6.0.94) (142) with 5-fold cross-validation to select the best value. Glmnet was fit with the glmnet R package (v4.1.8) (137) setting the α parameter to 0.5 (*i.e.* “elastic net” penalty that is half way between ridge and LASSO (143,144)) with the main penalty parameter lambda chosen by 5-fold cross validation. Missing values were imputed by k-nearest neighbors imputation (impute R package v1.74.1, default settings) (125).

Prediction accuracy was assessed by Pearson’s correlation for continuous features and balanced accuracy (BA = (sensitivity + specificity) / 2) for categorical features. For categorical features with three or more categories, BA was calculated separately for each category using a “one-versus-all” approach (*i.e.* a given category compared to all other categories combined) and then averaged. The predictors consisted of prostate cancer gene-level methylation for 13,603 genes, defined as the median β-value among CpG islands in the gene promoter region. Feature selection methods were performed in the training dataset in an attempt to improve performance and identify a smaller subset of important predictors. Specifically, the univariate association of each gene methylation predictor with each clinical feature and driver mutation was assessed using Kruskal-Wallis rank-sum test for categorical features and Spearman’s correlation for continuous features. To account for cohort differences, linear regression was used to regress each gene against cohort, and the residuals were used in the aforementioned univariate test. Then models were fit using only the top 10, 50, 100, 1000 or 5000 most significant genes, as well as a model that considered all 13,603 genes. To encourage a more parsimonious model, the final model for predicting a given feature was chosen as the model with the smallest number of predictors that was within 0.02 of the best Pearson’s correlation or balanced accuracy for numeric and categorical features, respectively. Lastly, using the test-dataset, patients were resampled with replacement and 95% bootstrap BCa confidence intervals were calculated for performance metrics using the boot R package (v1.3.28.1) (145).

### Predicting time-until-BCR using multi-omic data

A total of 774 localized prostate cancer patients with time-until biochemical recurrence (BCR) and multi-omic data (methylation, RNA, CNA) were randomly split into a 70% train and 30% test dataset. Two machine learning models were compared for predicting time-until-BCR: random survival forest (146) and a regularized Cox proportional hazards model (147), with prediction accuracy assessed using the C-index (148). The random forest was fit using the randomForestSRC R package (v3.2.30) (146) with 1000 trees and default settings, while the regularized Cox model was fit with the glmnet R package (v4.1.8) (147) setting the α parameter to 0.5 (*i.e.* “elastic net” penalty that is half way between ridge and LASSO (143,144)) with the main penalty parameter lambda chosen by using 10-fold cross validation to maximize the C-index in the training data. Missing values were imputed by k-nearest neighbors imputation (impute R package v1.74.1, default settings) (125).

Several combinations of data types were considered for model training (**Supplementary Figure 17a**). Given the large number of potential features, feature selection was performed in the training dataset to identify a smaller subset of potential predictors. To perform feature selection, a separate Cox model was fit to each feature treating it as a predictor of time-until-BCR, while adjusting for the pre-treatment clinical features. Then only features with Wald-test p-value < α were retained. Several values for α were considered (0.001, 0.005, 0.01, 0.05, and 1) with the final value of α = 0.005 chosen that maximized the random forest C-index in the training and test datasets (**Supplementary Figure 17d**).

The best machine learning model (highest C-index on test-dataset) was compared to the test C-index from a model that only considered pre-treatment clinical features. Using the test-dataset, patients were resampled with replacement and a 95% bootstrap BCa confidence interval was calculated for the difference in C-indices between the two models, using 50,000 bootstrap replications with the boot R package (v1.3.28.1) (145). Similarly, a 95% bootstrap BCa CI was calculated for the test C-index of the best performing model. Lastly, the predicted BCR risk score from the best model was compared between 52 non-malignant TCGA samples, 161 localized prostate cancer samples from the test dataset, and 99 metastatic samples. Given that the best model included PSA and ISUP grade as predictors, and these were not measured for the non-malignant samples (ISUP grade was not measured for metastatic samples), their values were imputed using the randomForestSRC impute.rfsrc R function (default settings) (126).

### DNA Methylation Age Correlations with Clinical and Genomic Features

DNA methylation (DNAm) age was computed using the methylclock R package (v1.4.0) (55) for cohorts with available continuous chronological age data (TCGA, ICGC PRAD-CA, Kirby *et al*. 2017). Whole-genome bisulfite sequencing datasets were excluded, as methylation clocks were trained on microarray data. Relative DNAm age acceleration was defined to capture proportional age-related changes under the assumption of linear aging in senior adult cancer patients:

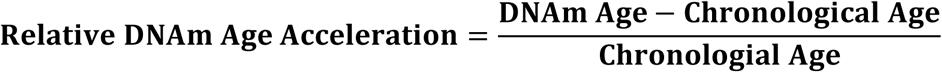

Relative DNAm age acceleration was compared across age categories, methylation subtypes, and ISUP grades using the Kruskal-Wallis rank sum test with epsilon-squared (ε²) effect size. Driver mutations present in ≥ 20% of tumour samples were analyzed in relation to relative DNAm age acceleration using the Mann-Whitney U test. False discovery rate (FDR)-adjusted q-values were computed. To account for chronological age, linear regression was applied to regress relative DNAm age acceleration against chronological age, and residuals were used in the aforementioned univariate tests.

## Data Availability

The data download sources for each cohort are given in **Supplementary Table 3**.

## Software Availability

The PrCaMethy R package offers tools for predicting methylation subtype, clinical features and somatic driver mutations using prostate cancer DNA methylation data generated from Illumina 450K and 850K arrays. The R package is publicly available on GitHub at: https://uclahs-cds.github.io/package-PrCaMethy/.

## Data Visualization

Data visualization was performed in the R statistical environment (v4.3.3 - 4.5.0) using the BPG package (v7.1.2)(149). Venn Diagrams were made using the VennDiagram package (v1.7.3) (150).

## Acknowledgments

The authors thank all members of the Boutros lab, especially Veronica Y. Sabelnykova, for helpful suggestions. The results described here are in part based on data generated by the TCGA Research Network. We greatly appreciate the West Coast Dream Team consortium for sharing processed methylation data from metastatic prostate tumour biopsies, and the authors of Kirby *et al.* 2017 for sharing clinical data.

## Author’s Contributions

Pathology analyses: TvdK

Data processing: JA, TNY, JO, TG, YJS, AF, RL, RH-W, CJ, RMAD, RA, JL, AS, CQY, SMGE, KEH, LEH, NZ

Statistical and machine-learning analyses: JA, AW, YJS, TNY, RMAD, KEH, FY

Wrote the first draft of the manuscript: JA, TNY, YJS, MF, PCB

Initiated the project: TvdK, RGB, MF, PCB

Supervised research: AK, AB, MLKC, ATP, MF, BP, NMC, TvdK, CMH, RGB, PCB

Approved the manuscript: all authors

---

**Supplementary Figure 1:**
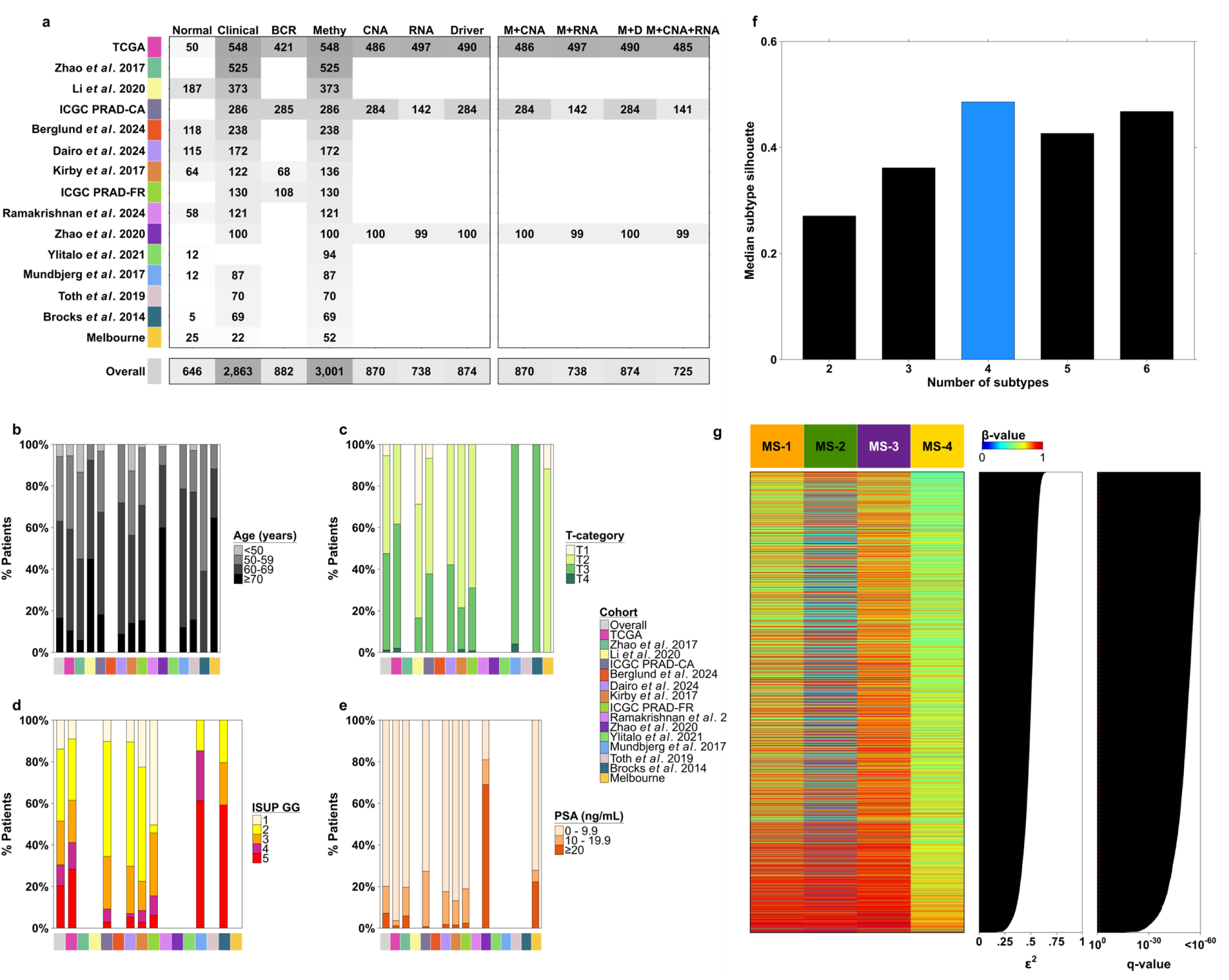
Sample size, pre-treatment clinical features and subtype-defining CpGs. **a.** Sample size by cohort (row) and data type (column). Columns: Clinical stands for pre-treatment clinical features; BCR is time-until biochemical recurrence; Driver is the panel of 28 driver mutations; M = methylation, D = driver. **b-e.** Stacked barplots showing the percent of patients in each category (y-axis) stratified by cohort (x-axis). T-category: tumour category; ISUP GG: International Society of Urologic Pathologists grade group; PSA: prostate-specific antigen. **f.** The silhouette method was used to choose the number of subtypes that maximized the median silhouette score, resulting in 4 subtypes (blue). **g.** Each of 5,486 subtype-defining CpGs (rows) is summarized by the median methylation β-value within each subtype (column). Kruskal-Wallis test was used to compare each CpG between the subtypes, with q-values displayed in the right barplot and epsilon-squared ɛ^2^ effect size in the middle barplot.

**Supplementary Figure 2:**
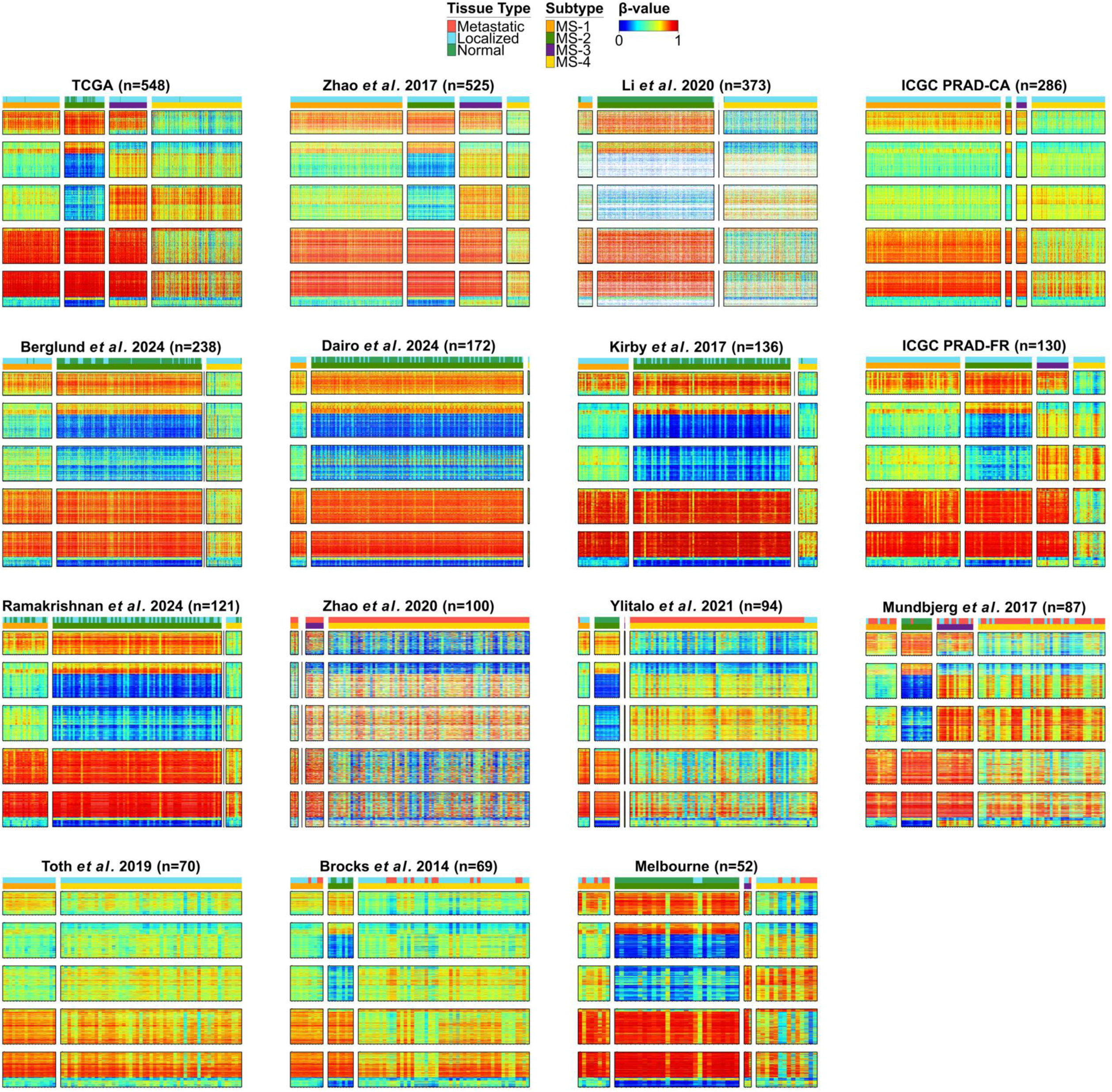
Subtype methylation profiles by cohort, tissue type. Methylation profiles stratified by cohort, subtype and tissue type (covariate bars beneath each cohort’s title). Patient-wise consensus clustering was performed using IMPACC to derive four subtypes within the TCGA cohort. The heatmaps include 5,486 CpGs which were selected by IMPACC as important for deriving the subtypes. A classifier was trained to classify all remaining samples to their respective subtypes identified in the TCGA cohort. CpGs were sorted using hierarchical clustering with complete linkage and Euclidean distance in the TCGA cohort, then split into five equally sized groups for easier visualization. Missing values are colored in white.

**Supplementary Figure 3:**
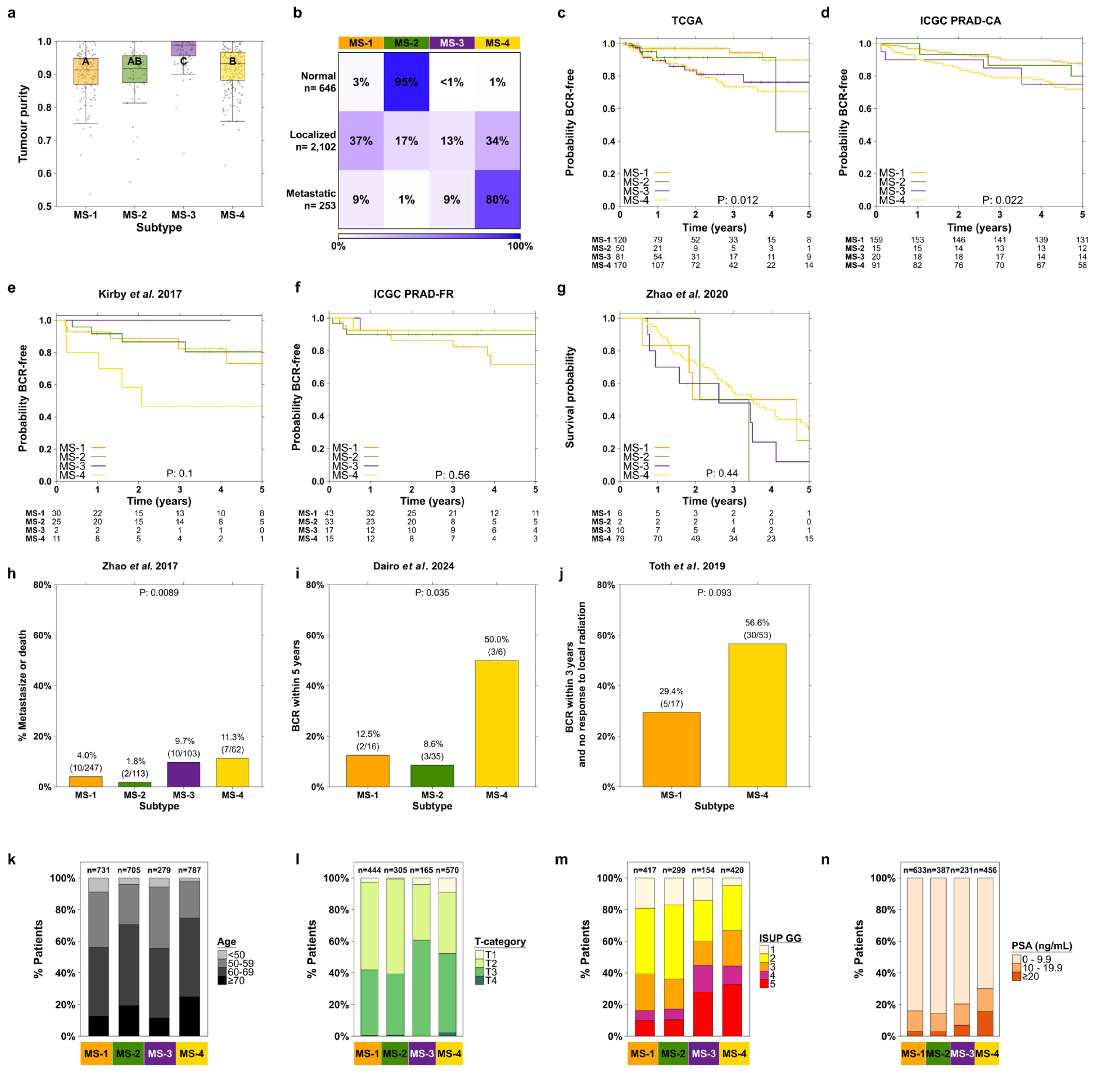
Methylation subtypes *vs.* tumour purity and clinical features. **a.** Tumour purity was compared between subtypes within the TCGA derivation cohort. Groups sharing the same letter do not significantly differ, U-test q > 0.05. **b.** Within each tissue type (row), the percent of samples assigned to each subtype (column) is shown. **c-j.** The subtypes were tested for association with time until biochemical recurrence (BCR) using a logrank test in cohorts (**c,d,f**) and a permutation logrank test in Kirby *et al.* 2017 (**e**), due to small/uneven group sample sizes (10,000 permutations). The permutation logrank test was used for time until death in Zhao *et al.* 2020 (**g**), while Fisher’s Exact Test was used for the binary outcomes of (**h-j**). **k-n.** Stacked barplots showing % of patients in each category (y-axis) stratified by methylation subtype (x-axis). T-category: tumour category; ISUP GG: International Society of Urologic Pathologists grade group; PSA: prostate-specific antigen.

**Supplementary Figure 4:**
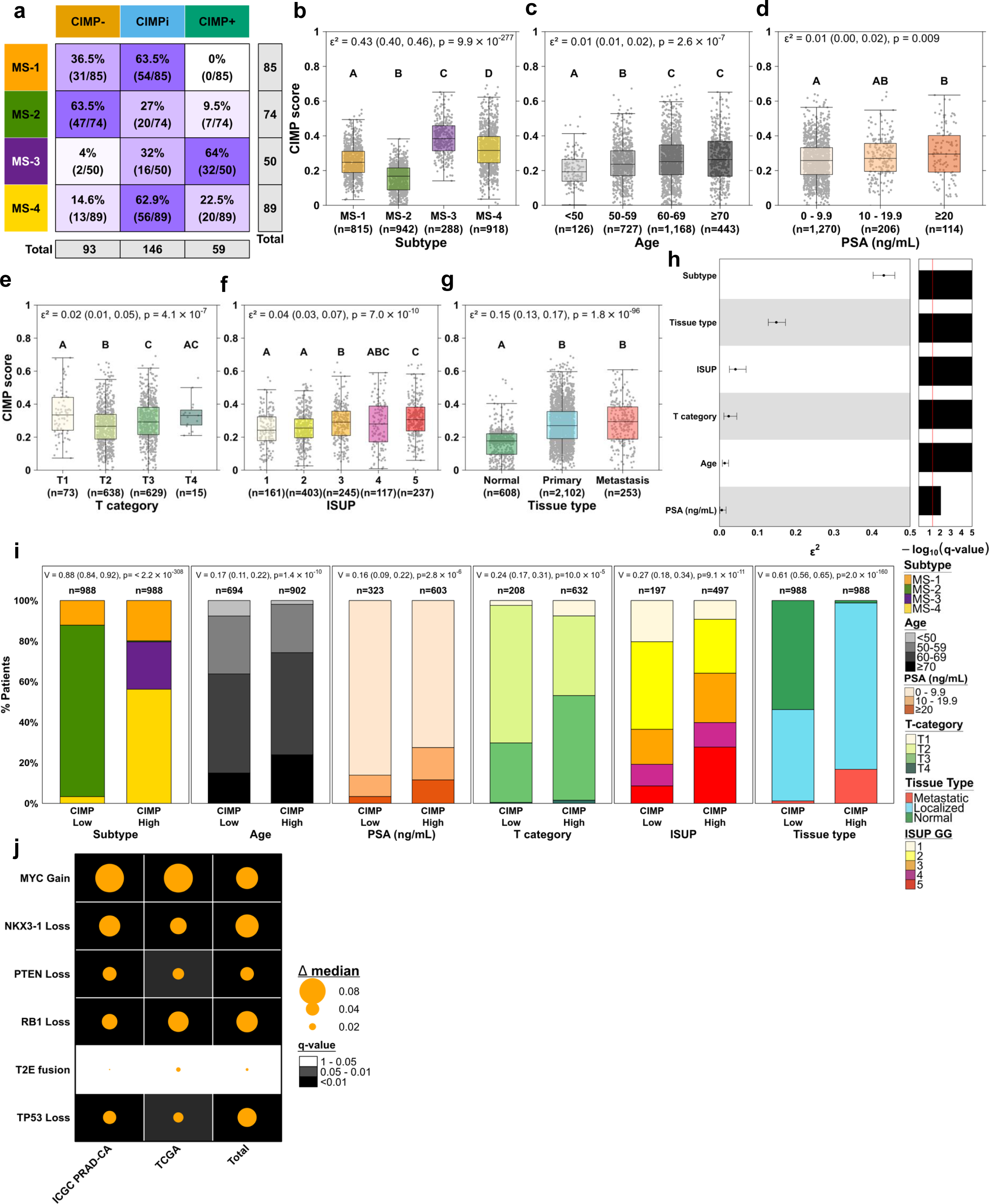
CIMP association patterns across MS subtypes and key features. **a.** Within each MS type (row), the percentage of samples assigned to each CIMP category (column) is shown (CIMP-, CIMP intermediate, CIMP+). **b-g.** CIMP scores were compared within MS subtypes and other key features after adjusting for platform and cohort effects. Groups sharing the same letter do not significantly differ (U-test q > 0.05). **h.** Summary statistics showing the effect sizes and q-values for the features in panels **b-g,** ordered by effect size. **i.** Distribution of feature categories across CIMP-low (0-33rd percentile) and CIMP-high (66-100th percentile) groups, shown as stacked percentages. **j.** Summary of associations between drivers and CIMP score in the ICGC PRAD-CA, TCGA, and combined cohorts. Dots represent effect sizes, with background colour indicating q-values

**Supplementary Figure 5:**
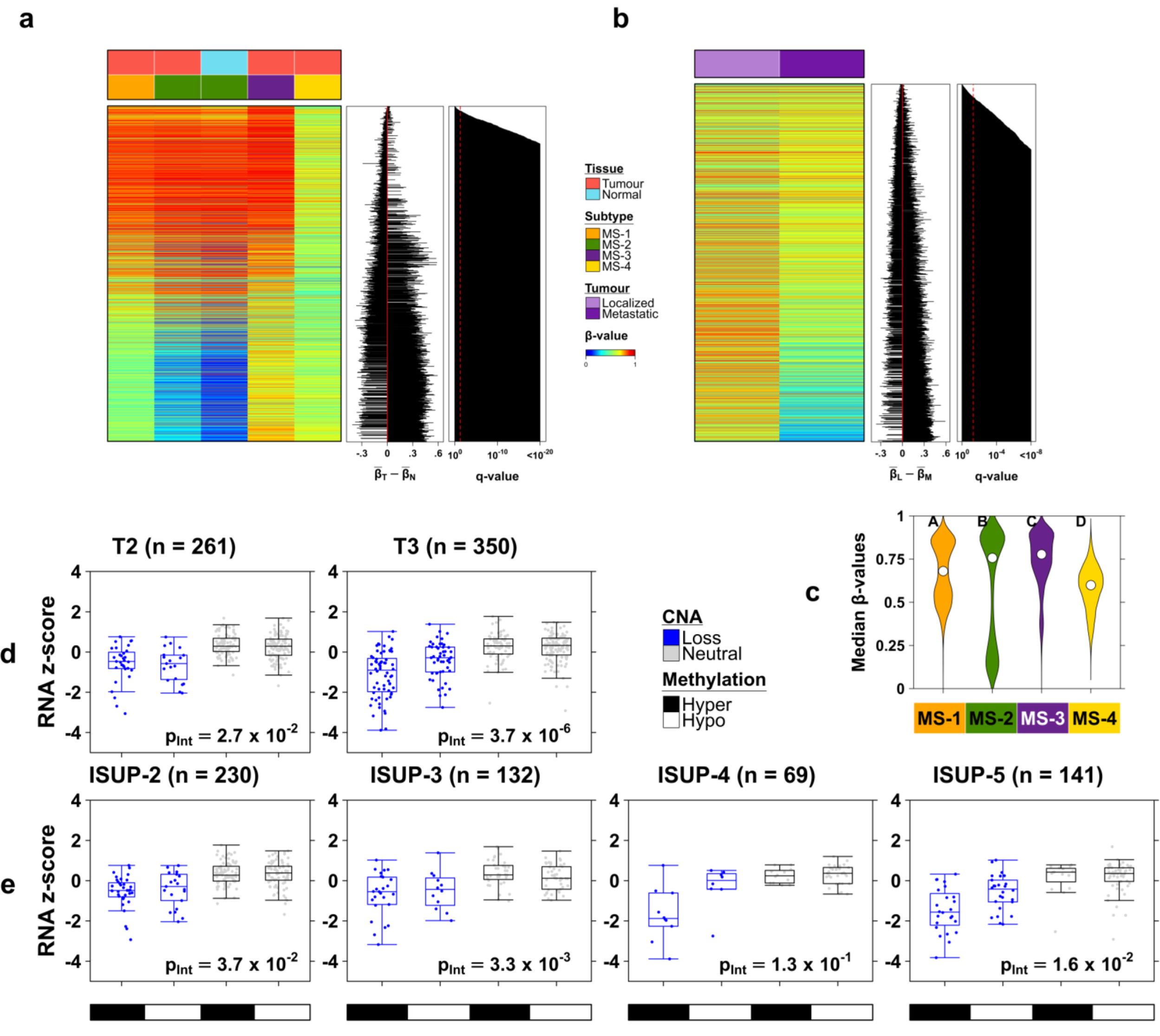
Subtype-defining CpGs *vs.* tissue type; *PTEN* interaction sensitivity analysis. **a. Tumour *vs.* normal samples**: Heatmap of 5,486 subtype-defining CpGs, sorted by |**β̅_T_ − β̅_N_**|, where **β̅_T_** is the median methylation beta-value in tumour samples, and **β̅_N_** is the median beta-value in normal samples. The Mann–Whitney U test was used to test for differences between the tumour and normal distributions of each CpG, with q-values displayed in the right barplot. **b. Lo**calized vs. metastatic samples: Similar to a., except CpGs sorted by |**β̅_L_** − **β̅_M_**|, where **β̅_L_** is the median methylation beta-value in localized prostate cancers, and **β̅_M_** is the median beta-value in metastatic prostate cancers. **c.** Distributions of the 5,486 subtype-defining CpGs (median methylation beta-value per CpG) compared between the four subtypes with only normal samples used for MS-2. The letters at the top of the plot indicate pairwise differences: if two groups have different letters, then they significantly differ (Wilcoxon test q < 0.05). **d,e.** A sensitivity analysis was performed for the *PTEN* methylation-CNA interaction on mRNA abundance (from Figure 2e**,f**) to determine whether the same association held across T-categories (**d**) and ISUP grade groups **(e)**. Linear regression was used to test the interaction between binary methylation (hyper vs. hypomethylation; median dichotomized) and binary CNA status (loss vs. neutral), with the interaction p-value given by p_Int_. The dependent variable is RNA abundance (z-score transformed log_2_ TPM). Boxplots are only shown for tumour categories and ISUP grades where each of the four groups had sample size greater than five.

**Supplementary Figure 6:**
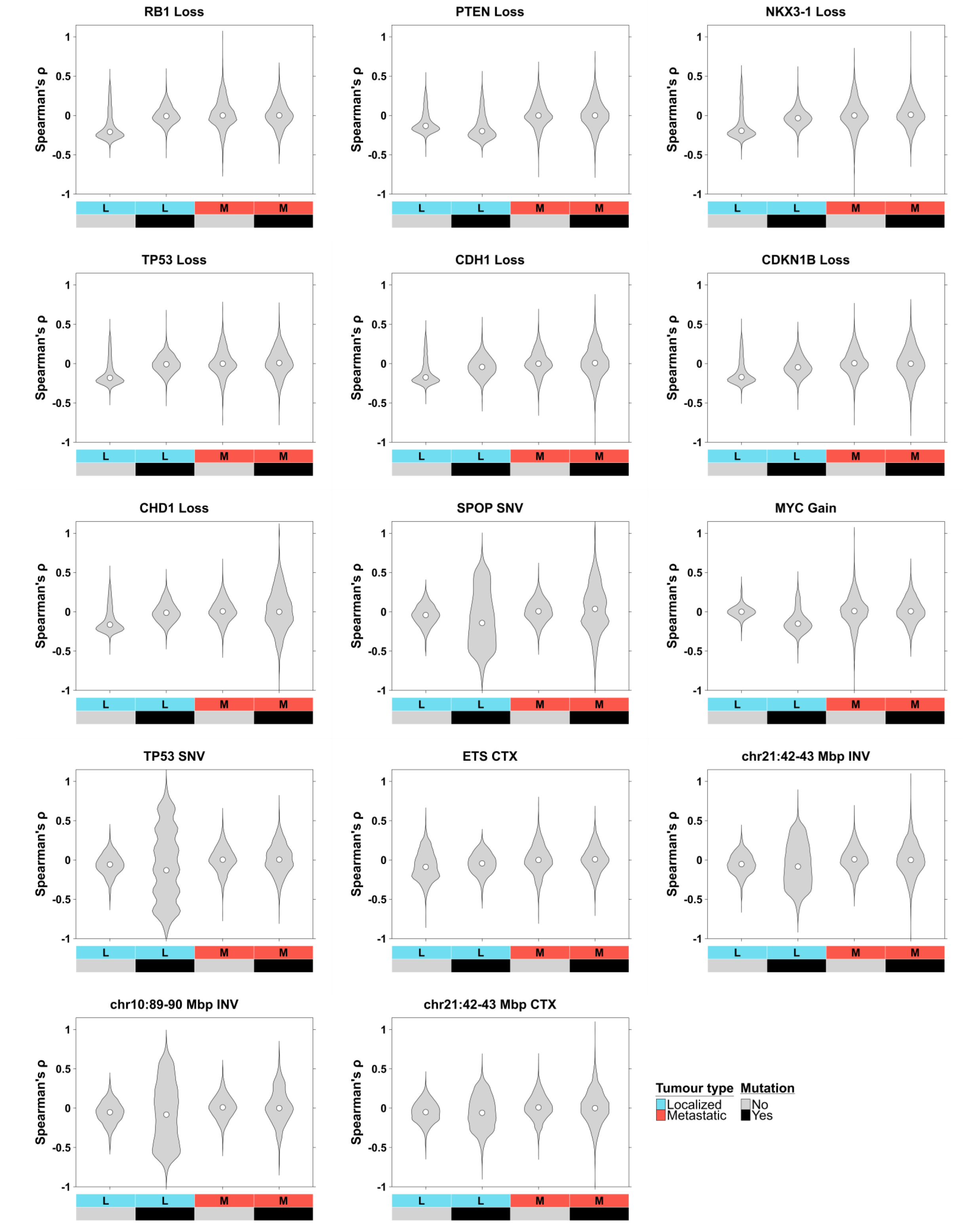
Distribution of methylation-CNA correlations stratified by 14 driver mutation events. For a given driver event, the methylation-CNA correlations are stratified by tumour type (localized, metastatic) and whether the driver mutation is present (yes/no).

**Supplementary Figure 7:**
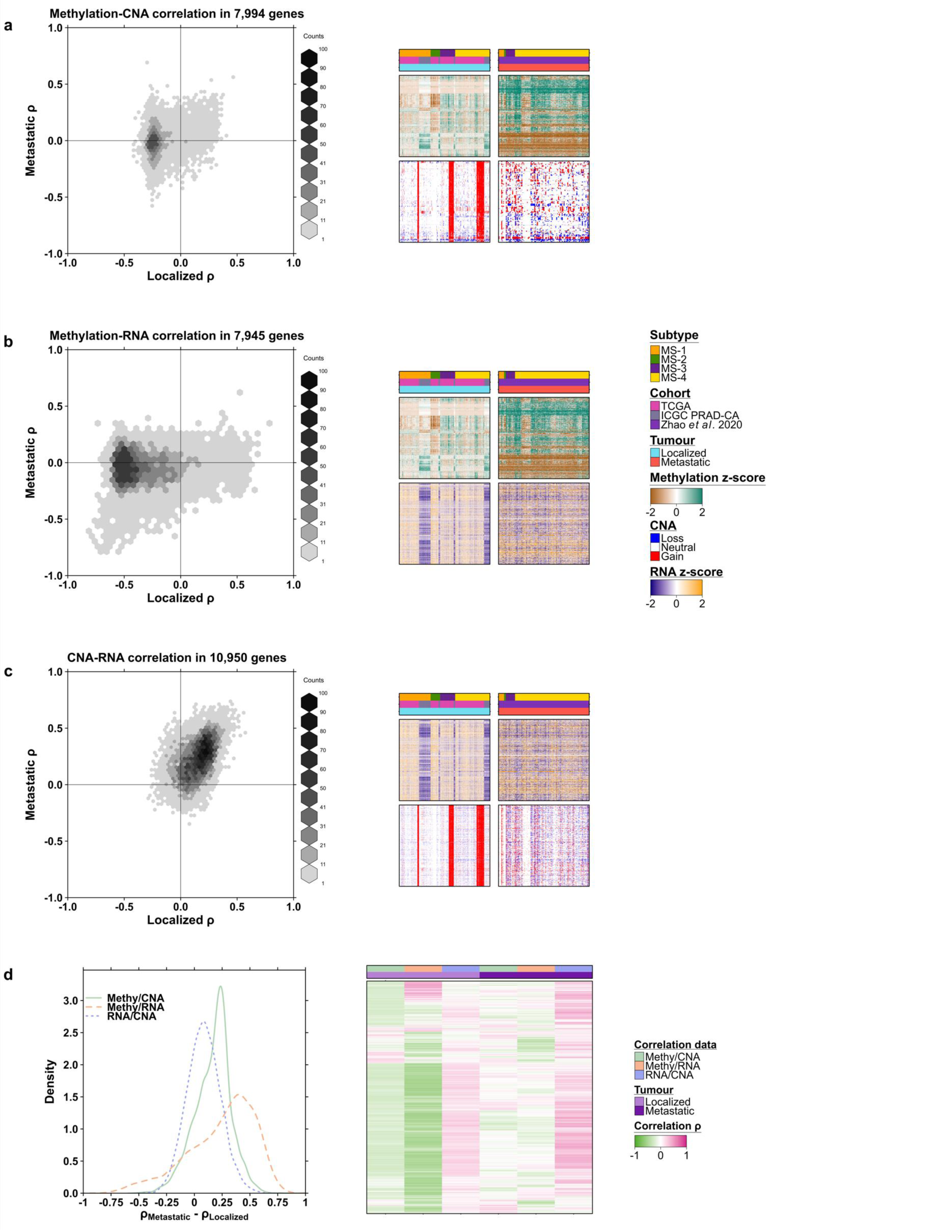
Multi-omic correlations differ between localized and metastatic samples. **a. Left:** Spearman’s correlation of gene-level methylation and CNA compared between localized and metastatic samples. **Right:** Heatmaps of methylation z-scores (top row) and CNA profiles (bottom row) stratified by tumour type. The heatmaps show the top 1,000 genes with the largest difference in median methylation between localized and metastatic samples. **b-c**. Similar to (a) except for methylation-RNA and CNA-RNA correlations. The right-hand heatmaps show the same set of 1,000 genes from the heatmap in (a). **d.** Left: differences in gene-level correlations by tumour type ρ_Metastatic_ - ρ_Localized_ for each pair of data types. Right: heatmap of gene correlations for each pair of data types stratified by localized and metastatic samples.

**Supplementary Figure 8:**
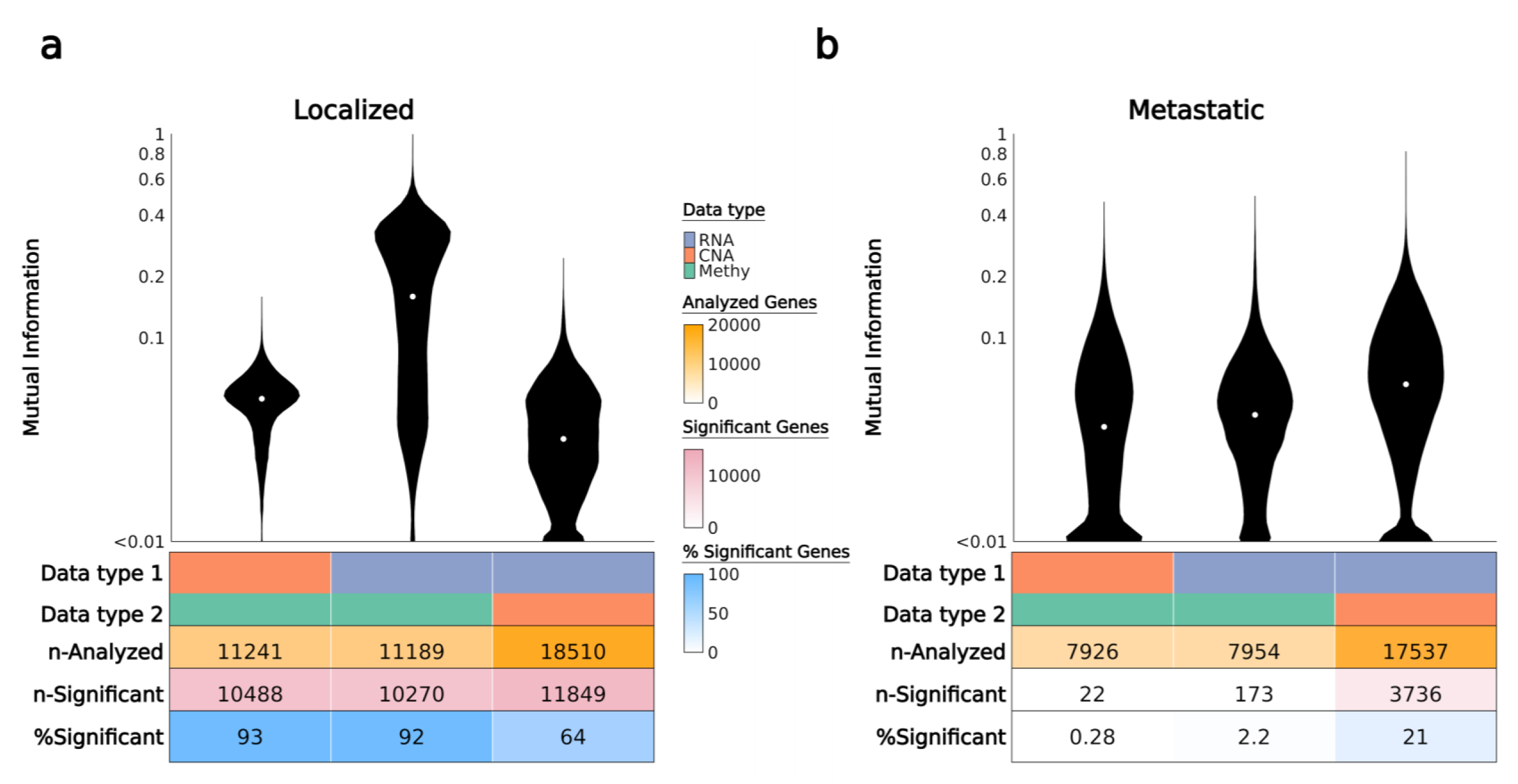
Mutual information of multi-omic data differs in localized and metastatic samples. **a-b.** Mutual information was calculated for all features within each pair of data types (RNA, CNA, methylation) and displayed in a violin plot. A permutation test with 1000 permutations was used to calculate p-values and results with multiple-testing adjusted q < 0.05 were regarded as statistically significant. Results were stratified by localized (**a**) and metastatic (**b**) prostate cancers.

**Supplementary Figure 9:**
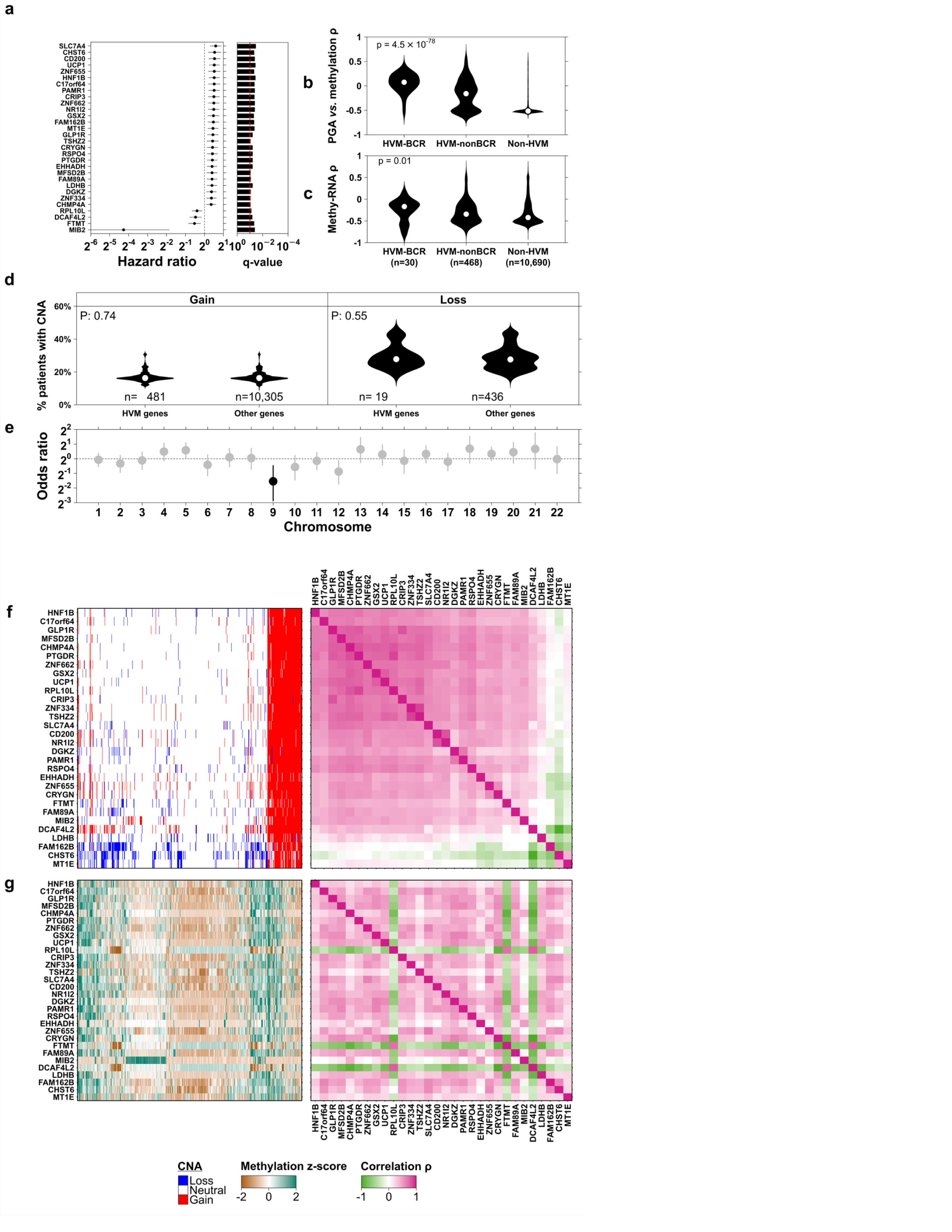
Genes with high variation in methylation (HVM) *vs.* BCR and CNA. **a.** 30/500 HVM genes significantly associated with time-until-BCR (Cox proportional hazards Wald test q < 0.10). Gene methylation measured by median beta-value of CpG islands in the gene promoter region. **b.** Distribution of Spearman correlations of gene methylation vs. PGA stratified by HVM genes that are associated with BCR (HVM-BCR; q < 0.10); not associated with BCR (HVM-nonBCR; q > 0.10) and non-HVM genes. **c.** Distribution of Spearman correlations for within-gene methylation and mRNA abundance, stratified by the same groups as part **b.** **d.** Percent of patients with CNA loss or gain was calculated separately for HVM and non-HVM genes and compared using Mann-Whitney U-test. **e.** The number of HVM and non-HVM genes was compared within each chromosome using Fisher’s exact test and odds ratio. Significant results shown in black, q < 0.10. **f. Left:** CNA profiles for the 30 HVM genes in **a**. Both rows (genes) and columns (patients) were sorted using hierarchical clustering with Euclidean distance and complete linkage. **Right:** Spearman’s correlation matrix for gene CNAs (rows and columns correspond to the same set of 30 genes). **g. Left:** Methylation profiles for the 30 HVM genes in **a**. Rows (genes) and columns (patients) were sorted to match **f**. **Right:** Spearman’s correlation matrix for gene methylation (rows and columns correspond to the same set of 30 genes).

**Supplementary Figure 10:**
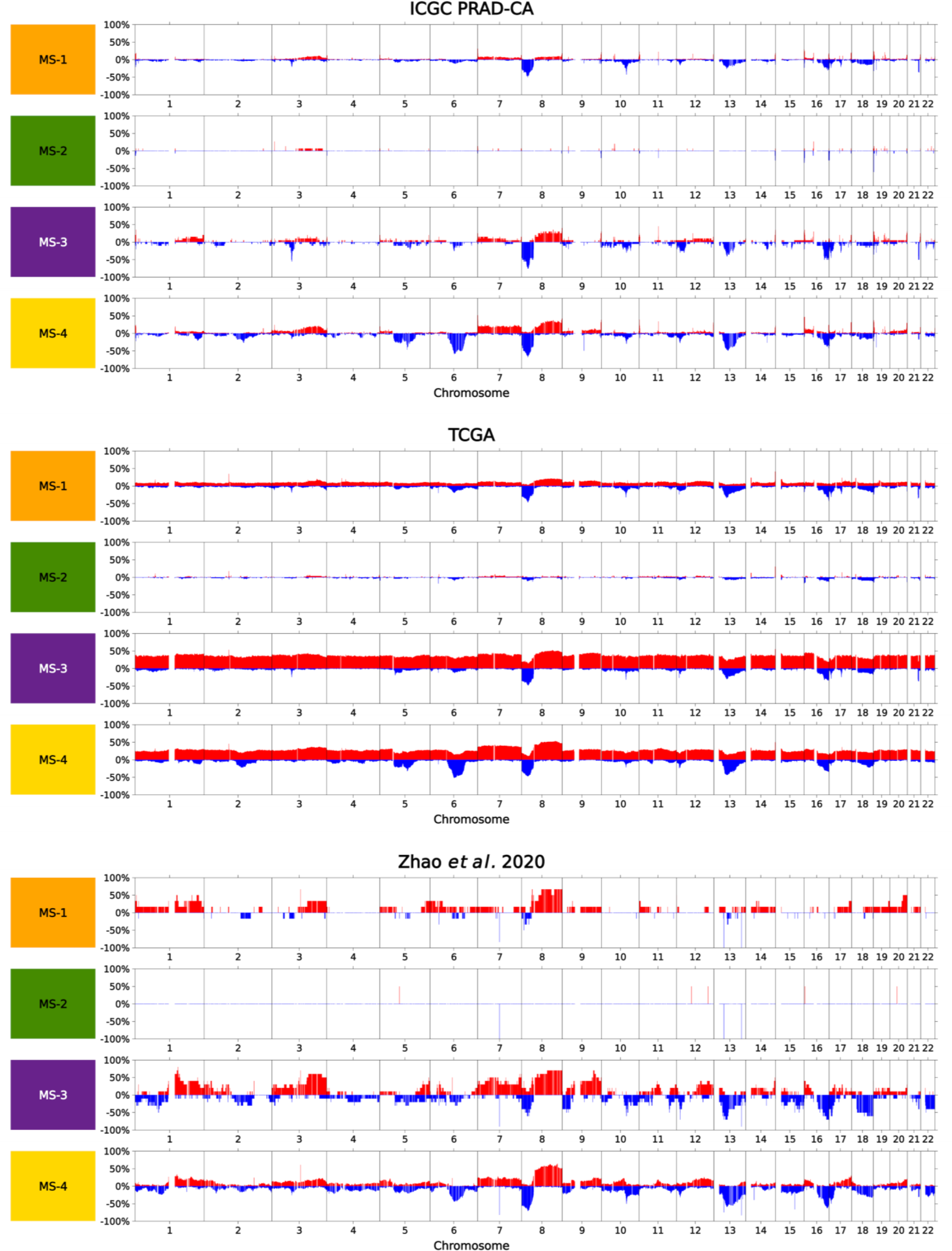
Genome-wide CNA profiles by subtype and cohort. The genome-wide copy number alteration (CNA) profile is shown for each subtype (row), stratified by the ICGC PRAD-CA, TCGA and Zhao *et al.* 2020 cohorts. Bars indicate the percent of patients with copy number gain or loss (y-axis) at each location of the genome (x-axis).

**Supplementary Figure 11:**
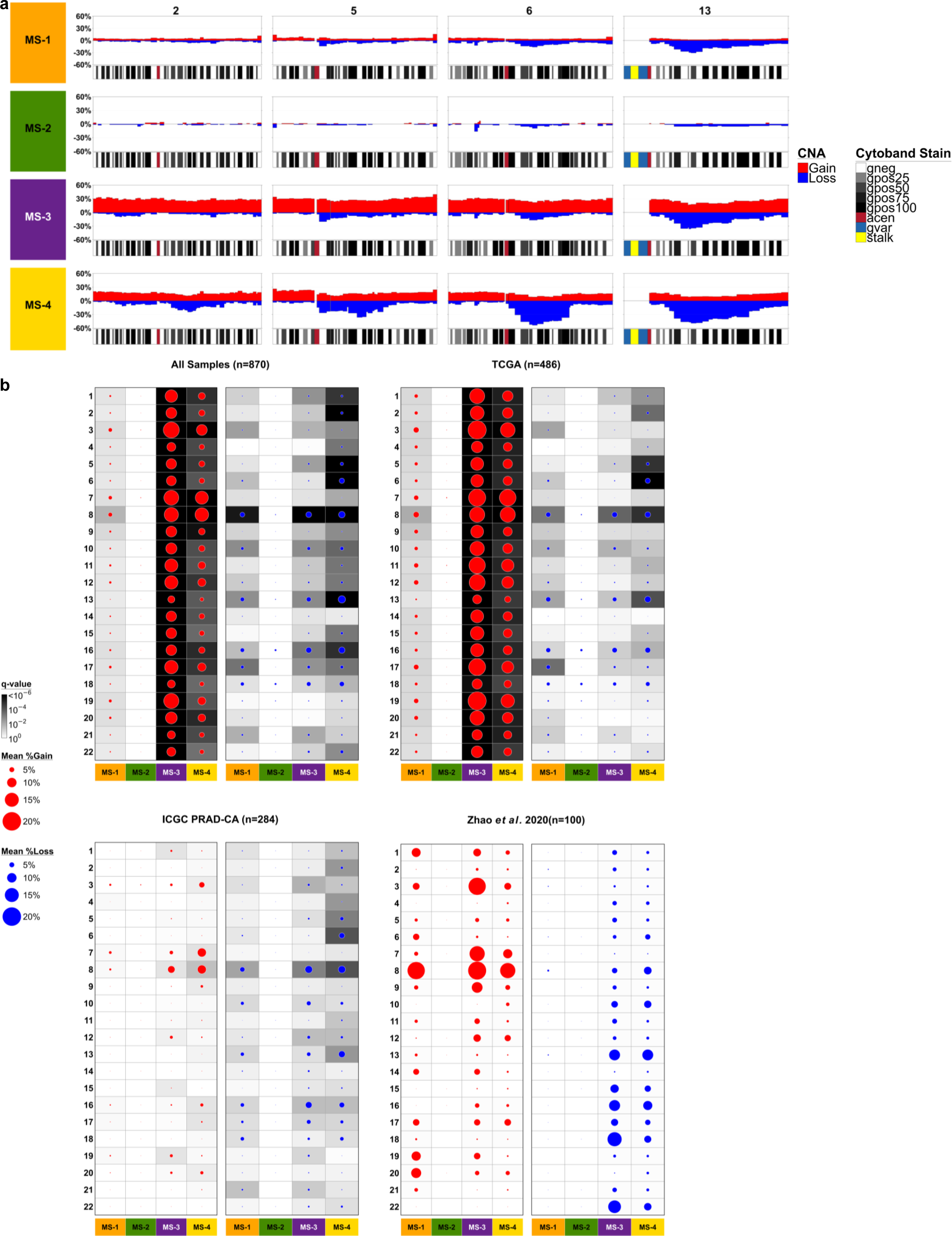
Chromosome-level CNA profiles by subtype and cohort. **a.** The y-axis shows the percent of patients with CNA gain (red) or loss (blue) in that region of the genome (x-axis), stratified by the four methylation subtypes (rows). Four chromosomes (2,5,6, and 13) are shown (columns) for which MS-4 had higher rates of CNA loss. Covariate bars below each plot show cytoband stains. **b.** For each patient, the percent of a given chromosome involved with a CNA loss or gain event was calculated. The mean percent of CNA gain in a given chromosome is shown by a red dot, while the mean percent CNA loss is shown by a blue dot. Results are shown when pooling all samples (top left), or stratified by cohort. The background shading in a cell indicates the q-value for testing whether the mean CNA rate differs from the MS-2 reference group.

**Supplementary Figure 12:**
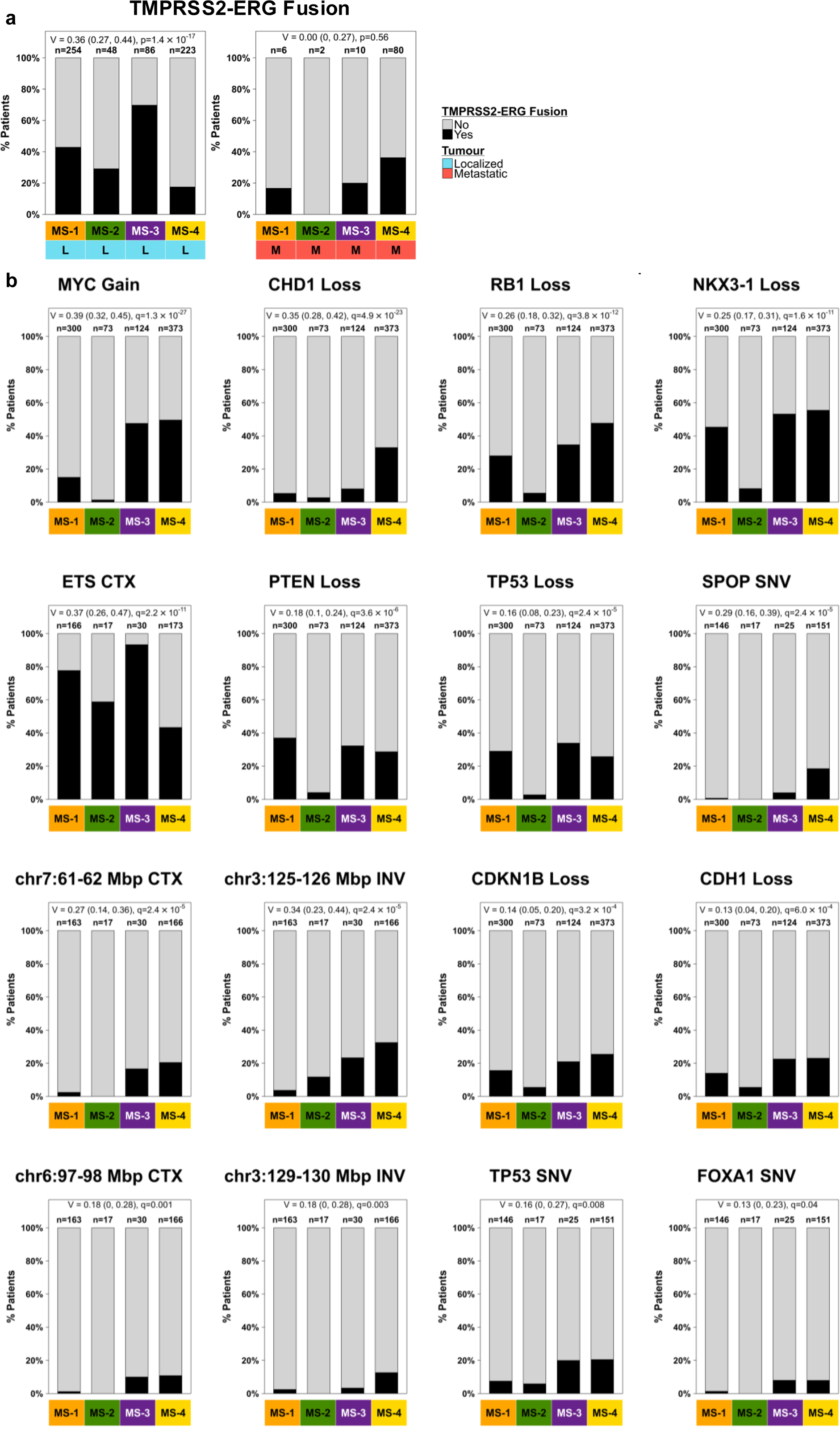
Distributions of categorical driver mutations stratified by the four methylation subtypes. **a.** TMPRSS2-ERG fusion rates (y-axis) are further stratified by localized (left) and metastatic (right) samples. **b.** Absolute distributions across the methylation subtypes for recurrent alterations. Bias-corrected Cramér’s V with 95% confidence interval and Fisher’s exact test q-value displayed at the top of each plot.

**Supplementary Figure 13:**
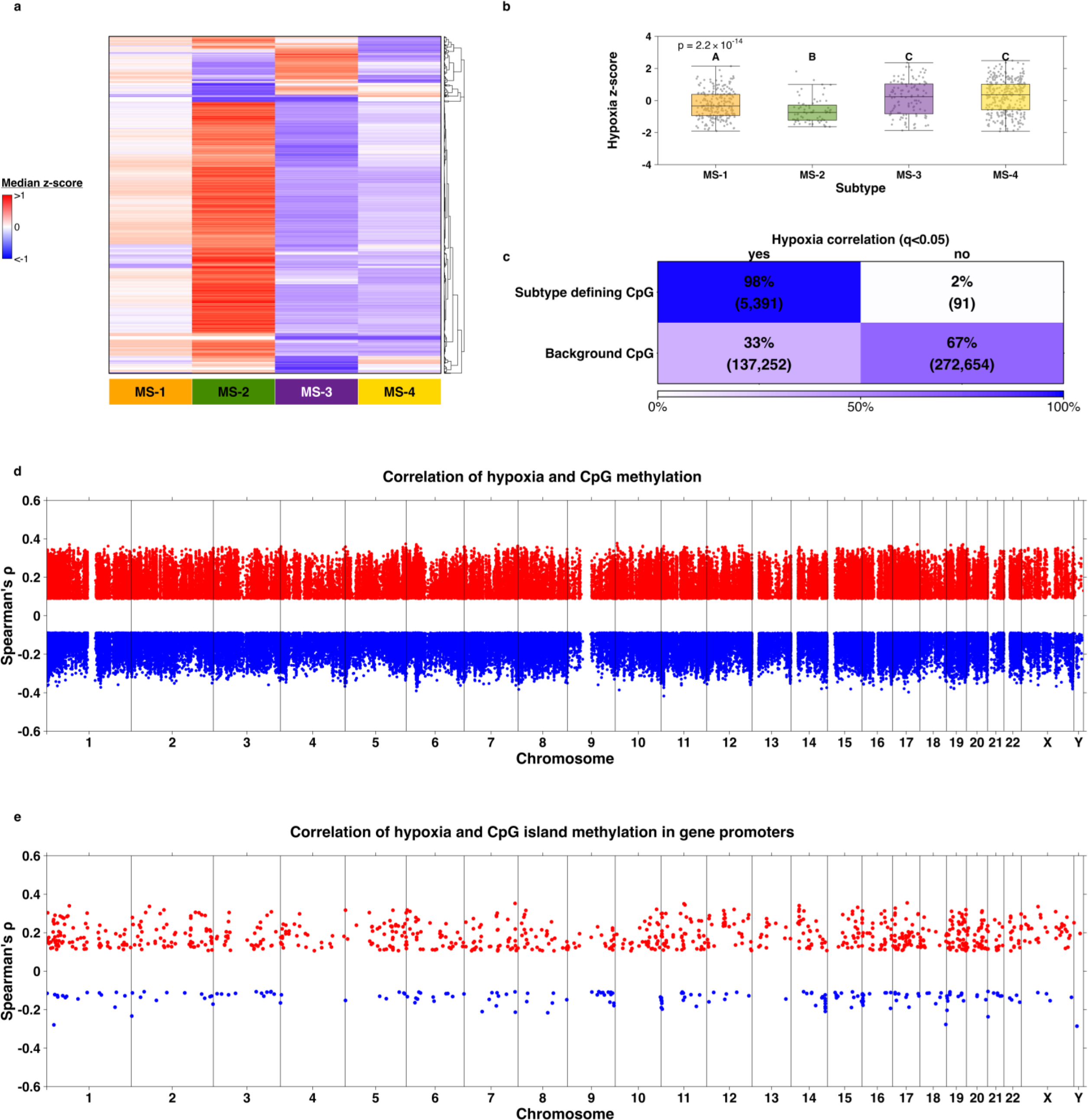
RNA *vs.* subtypes and hypoxia *vs.* methylation. **a.** Heatmap of median adjusted RNA abundance z-score (adjusted for cohort and PGA) by subtype for 3,516 significant transcripts (Kruskal Wallis q < 0.05, standard deviation (SD) of subtype median z-scores > 0.4). **b.** Hypoxia z-score was compared between subtypes using Kruskal-Wallis rank sum test. Pairwise comparisons between groups were conducted using the Mann-Whitney U-test, such that groups with different letters significantly differ (q < 0.05). **c.** Spearman’s correlation was calculated for each CpG with hypoxia score. The table compares the rate of significant CpG-hypoxia correlation (q < 0.05) between subtype-defining and non-subtype-defining background CpGs. **d.** Spearman’s correlation between each CpG and hypoxia score. 142,643 significant CpGs shown (q < 0.05) out of 415,388 tested **e.** Spearman’s corr. of hypoxia and methylation of CpG islands in gene promoters. 1,046 significant genes shown (q < 0.05) out of 13,890 tested.

**Supplementary Figure 14:**
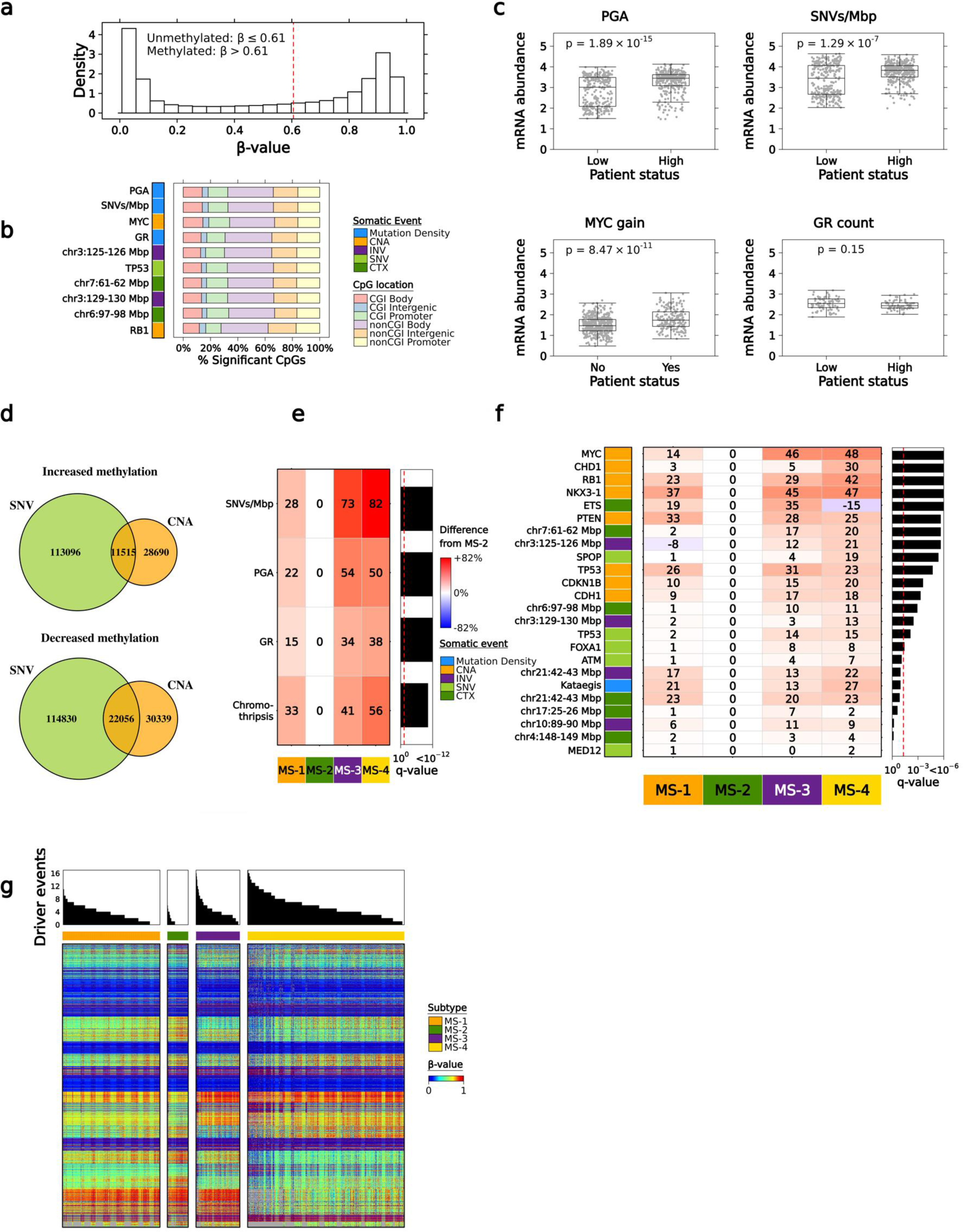
Driver events *vs.* methylation. **a.** The median methylation β-value was calculated for each CpG and displayed as a histogram (498 TCGA patients; 418,822 CpGs). The red dotted line represents a median β-value = 0.61. **b.** Top 10 driver events with the most number of significant CpG associations (q < 0.05). CpG locations were categorized into six groups with a combination of CpG island/non-CpG island and different genomic regions (gene promoter, body and intergenic). Percentages were calculated based on how many significant differentially methylated CpGs in each group. **c.** mRNA abundance levels of the top four driver events with the most number of significant CpG associations. For a given driver event, patient-level mRNA abundance was calculated as the median abundance among all genes that map to the top 100 most significant CpG associations. SNV count and PGA were median-dichotomized into high *vs.* low groups. P-value was generated from the Mann-Whitney U-test. **d.** Compares the overlap of CpGs significantly associated (q < 0.05) with *TP53* SNV (green) and *TP53* CNA loss (orange). Positive and inverse associations are shown in the top and bottom Venn Diagrams, respectively. **e.** Global mutation burden *vs*. subtypes: the median percentile was calculated for each subtype then subtracted from MS-2; Kruskal-Wallis test q-value. **f.** Driver mutation burden *vs*. subtypes: the percent of patients with the driver event was calculated for each subtype then subtracted from MS-2; chi-square test q-value (or Fisher’s exact test if 2x2 table or cell with < 5 expected counts). **g.** Heatmap of union of top 100 significant CpGs from each driver event (2,196 total CpGs). For each patient, the total number of driver events was calculated by summing binary indicator variables. Continuous drivers were median dichotomized into high *vs.* low, with high adding to the total score.

**Supplementary Figure 15:**
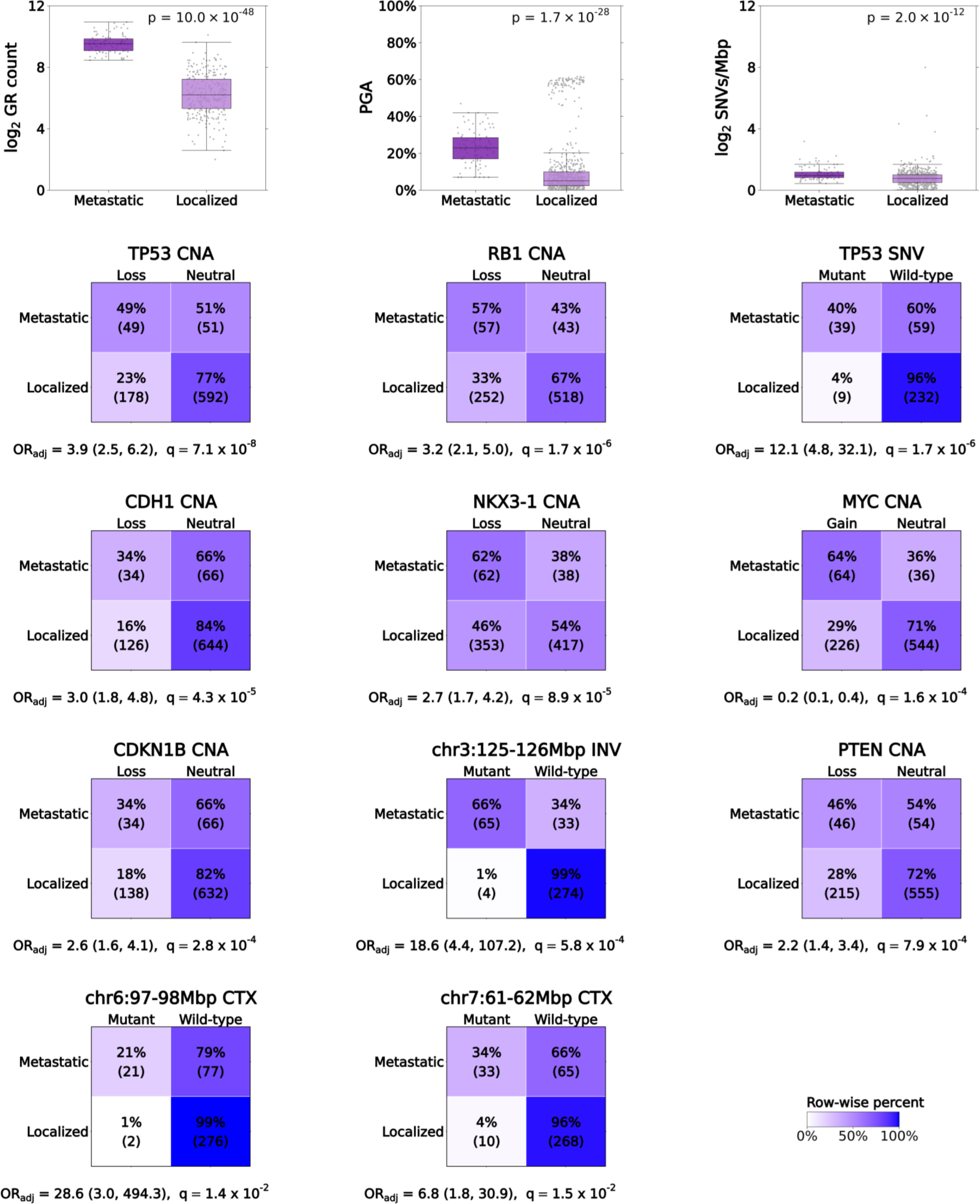
Driver mutations in localized *vs.* metastatic prostate cancers. Mann-Whitney U-test p-value shown for continuous drivers (top row). For categorical drivers, logistic regression odds ratio (OR_adj_ with 95% confidence interval) adjusting for total mutation burden (CNA drivers adjusted for PGA, SNVs adjusted for log_2_ SNVs/Mbp, and genomic rearrangements adjusted for log_2_ GR count). INV = inversion, CTX = translocation.

**Supplementary Figure 16:**
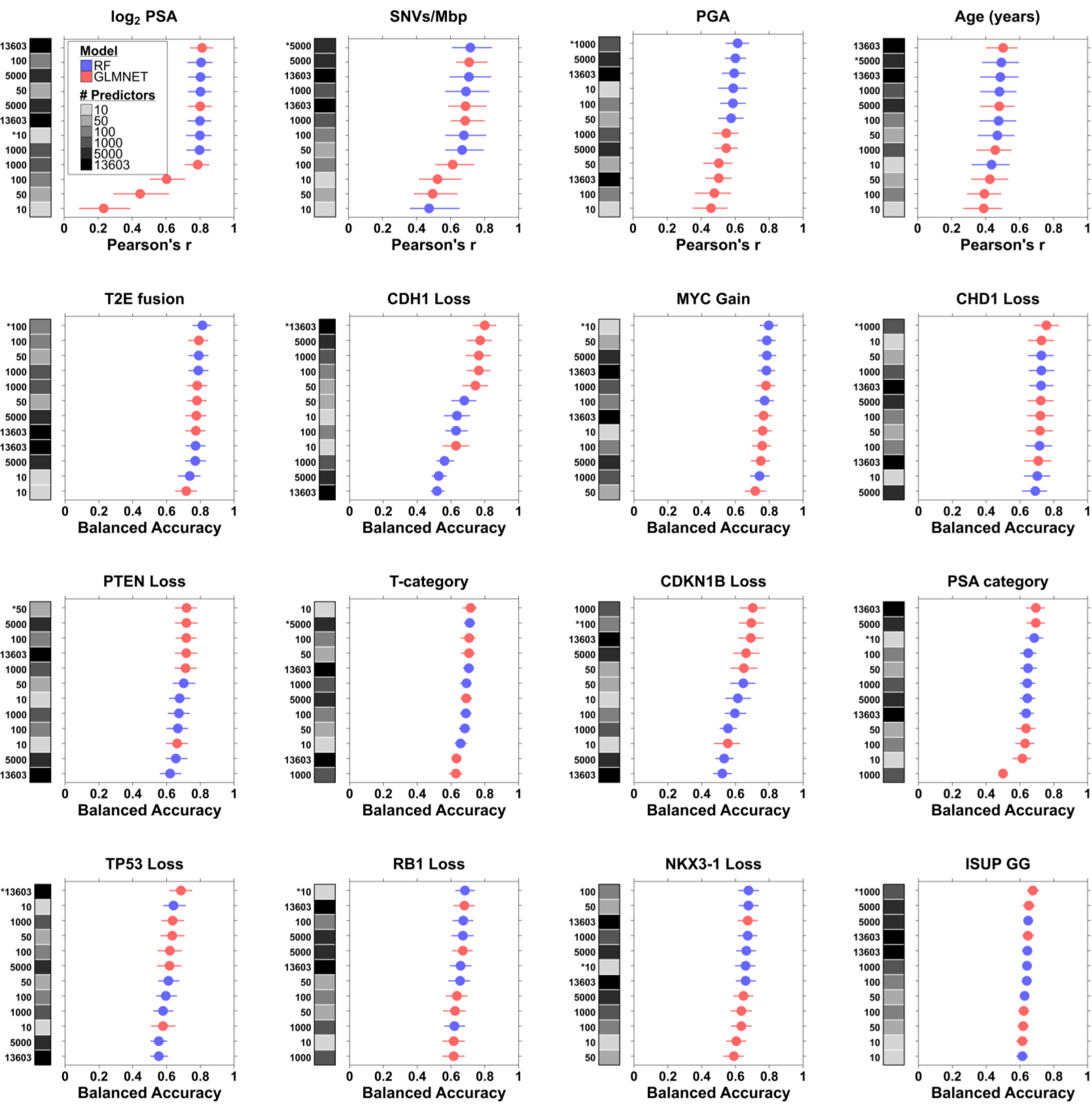
Performance on the test dataset after feature selection. For a given plot, the left black-grey covariate bar indicates how many features were kept after feature selection (see Methods). Pearson’s correlation and balanced accuracy were used for numeric and categorical features, respectively. An asterisk in the black-grey covariate bar indicates the final selected model, which was the smallest model within 0.02 of the best model. Results for random forest are shown in blue and glmnet in red. 95% bootstrap confidence intervals are indicated by the lines beside each point.

**Supplementary Figure 17:**
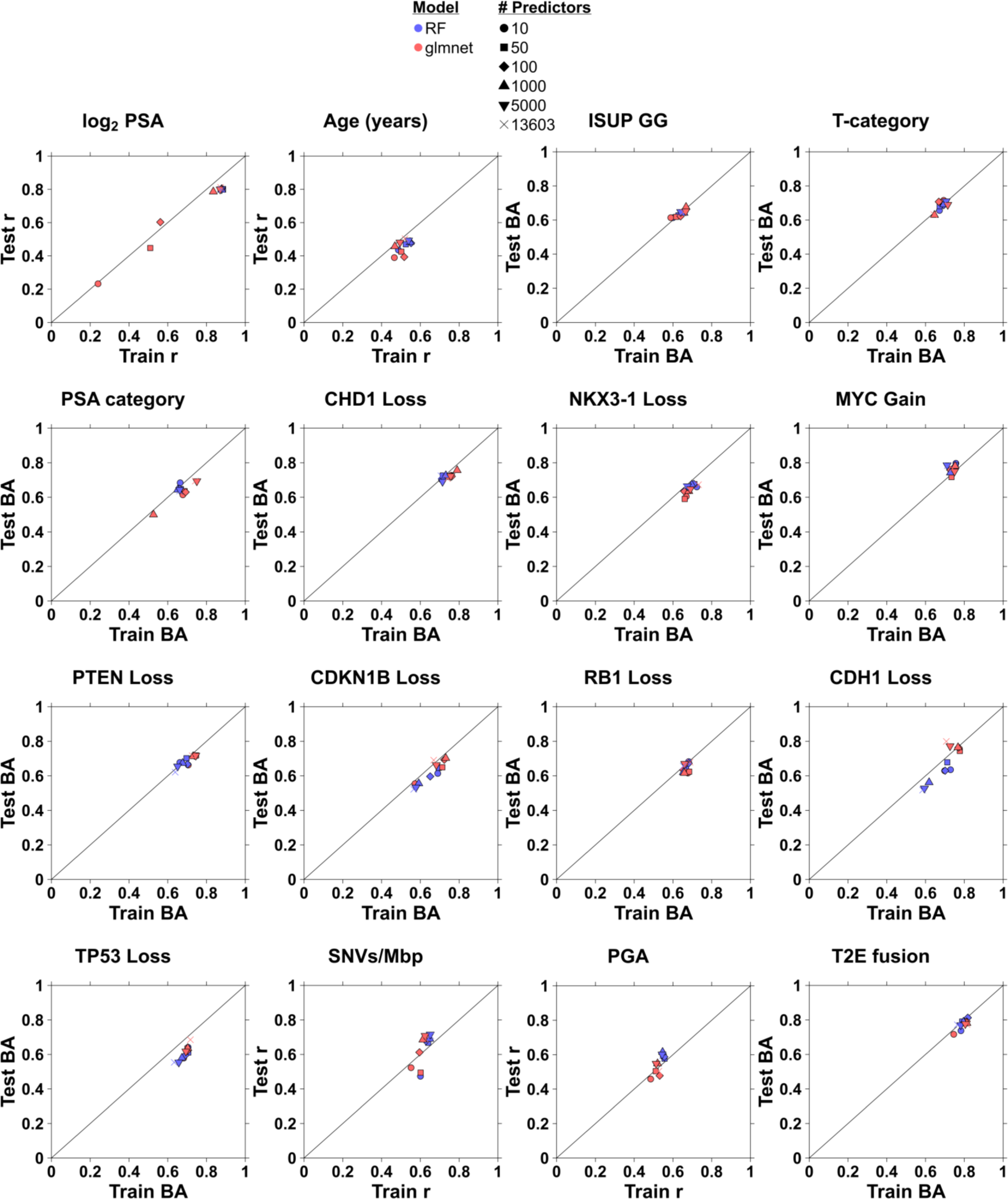
Performance on the training and test datasets for all models. For a given scatterplot, the model performance is shown for the training dataset (x-axis) and test dataset (y-axis) for random forest (RF; blue) and glmnet (red), and different combinations of number of predictors used after feature selection (see legend for shapes of points). Pearson’s correlation (r) and balanced accuracy (BA) were used for numeric and categorical features, respectively.

**Supplementary Figure 18:**
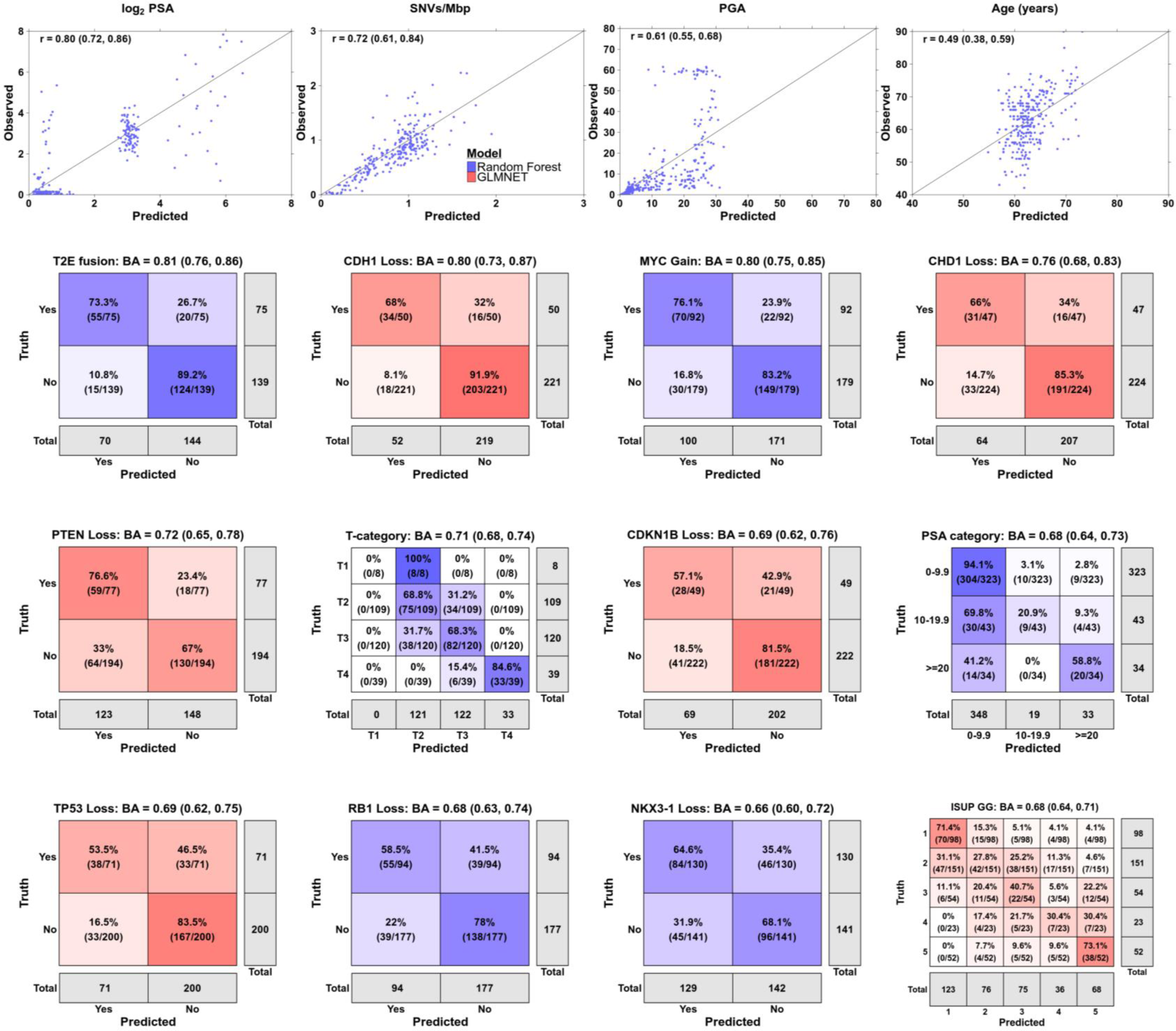
Performance of the final models on the test dataset. Results are shown for the final selected models (see Methods for model section details), using scatterplots for numeric outcomes and confusion matrices for categorical. Results for random forest are shown in blue and glmnet in red. The shading of the confusion matrices is determined by the row-wise percentages, with higher percentages given darker color. Pearson’s correlation is in the upper left corner of scatterplots and balanced accuracy (BA) in the title of the confusion matrix; 95% bootstrap confidence intervals are shown in parentheses.

**Supplementary Figure 19:**
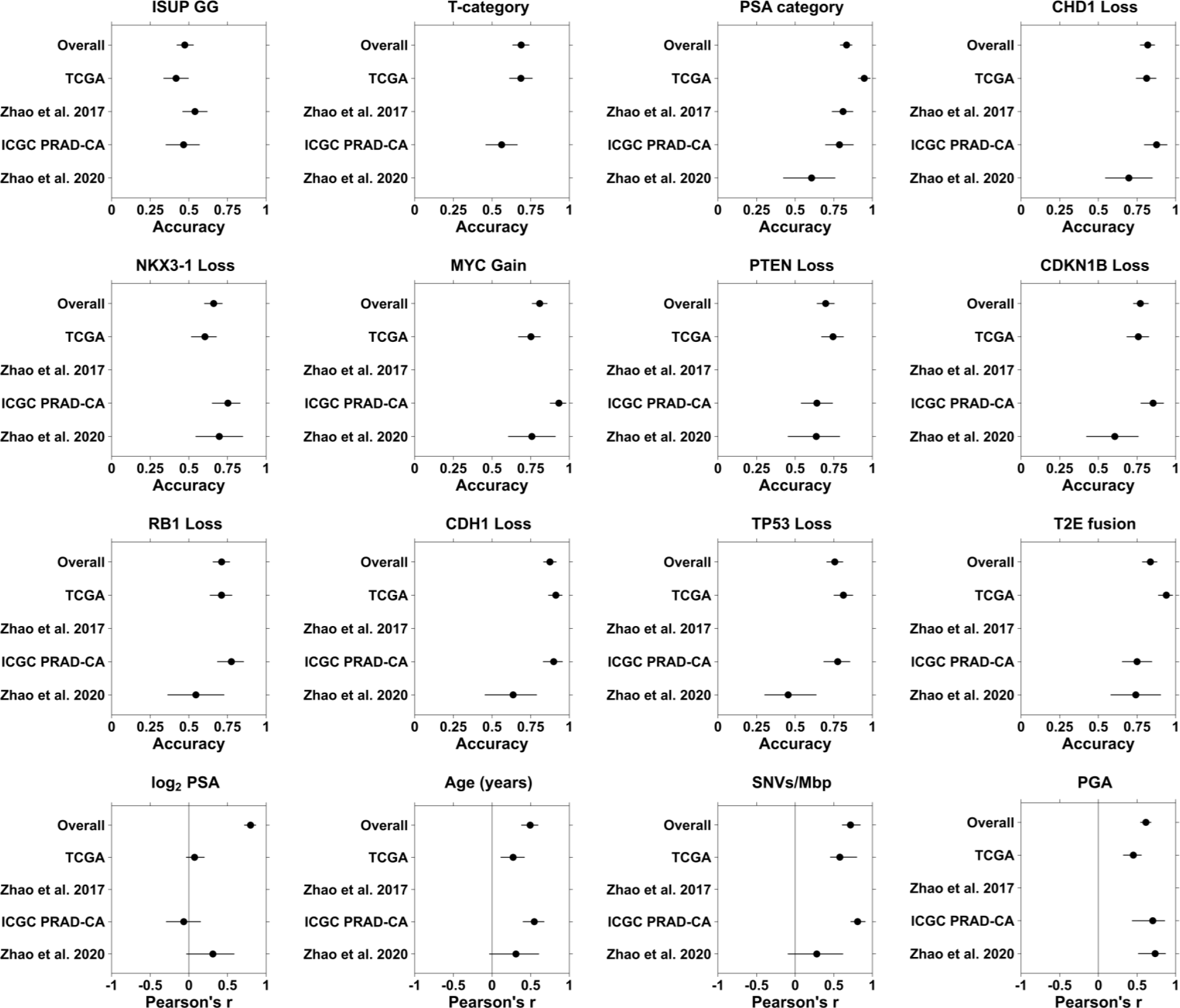
Model performance on the test dataset stratified by cohort. For a given plot, results are shown for the final model when pooling all cohorts together (“Overall”) or when fit separately by cohort as indicated in the y-axis labels. Pearson’s r is used for numeric outcomes and accuracy for categorical outcomes (% of observations correctly classified). Although balanced accuracy (BA) was used to tune and select the final models for categorical outcomes, accuracy is shown here since we were unable to calculate BA for some of the cohorts due to small cell counts resulting in the BA being undefined. 95% bootstrap confidence intervals are indicated by the lines beside each point.

**Supplementary Figure 20:**
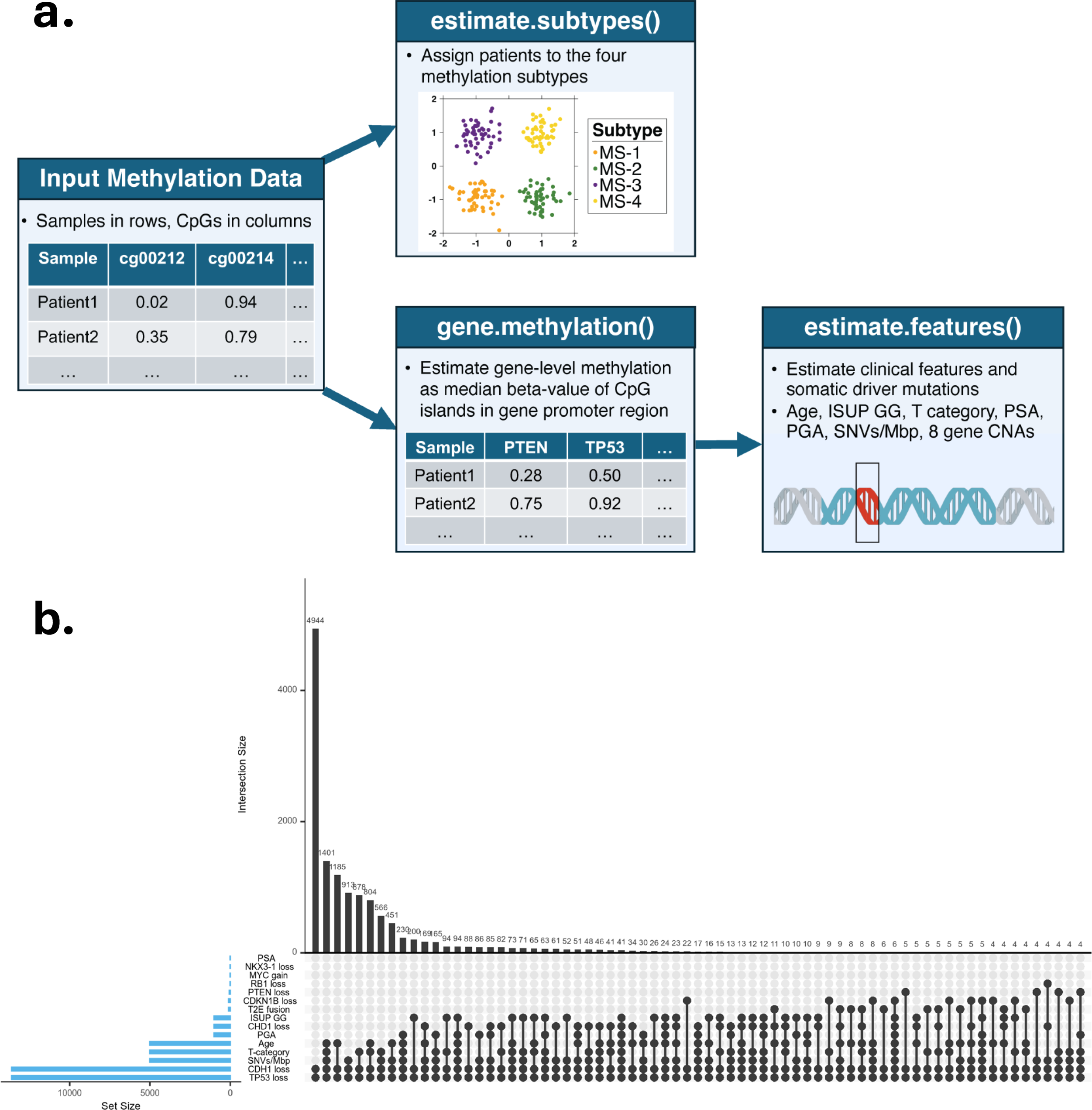
PrCaMethy R package. **a.** User input is a data.frame of methylation CpGs from the Illumina 450K or 850K arrays. Samples are in rows and CpGs in columns. The matrix contains beta-values (continuous proportions ranging from 0 to 1). Using this input dataset, the flowchart demonstrates the workflow of the 3 main R functions. **b.** Overlap of predictors used between the 14 models. Left barplot shows total genes per model. Top barplot shows overlap of genes for various combinations of models. For a given column of the barplot, the black dots with lines indicate which models are represented by that bar. The number above the bar indicates the number of genes shared in common between those models. Combinations that share at least three genes are shown.

**Supplementary Figure 21:**
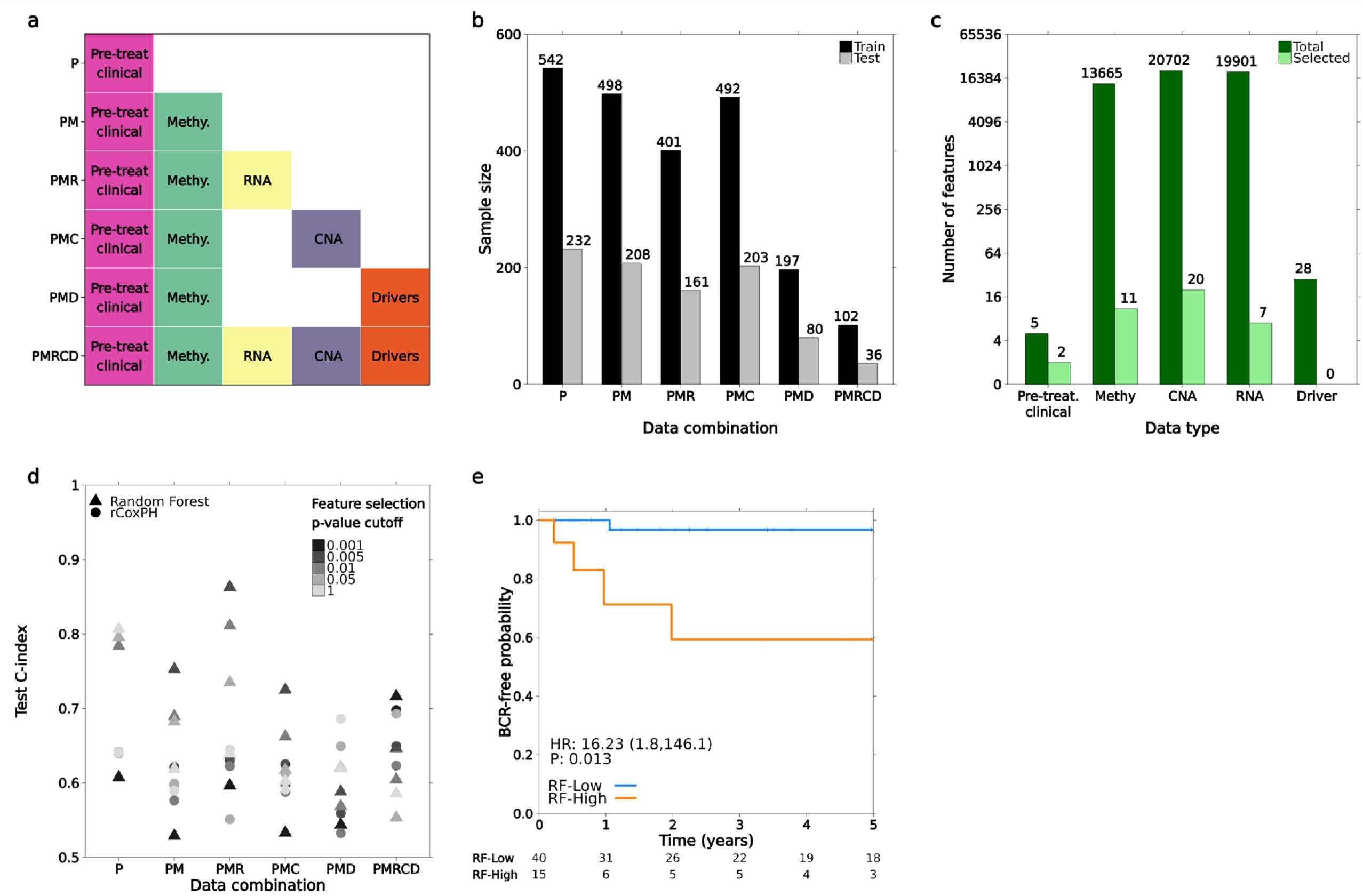
Multi-omic data for predicting time-until-BCR. **a.** Models for predicting time-until-BCR were built using different combinations of data types (Pre-treatment clinical, methylation, RNA, CNA and somatic driver mutations). The acronyms on the y-axis labels indicate the combinations of data types and correspond to the x-axis labels in parts **(b)** and **(d)**. **b.** A total of 774 localized prostate cancer patients with available time-until-BCR data were randomly split into 70% Train (black) and 30% Test datasets (grey). The height of bars indicates the train and test set sample sizes for the different combinations of data types in **(a)**. **c.** The height of bars indicates the total number of features (dark green) and the number of features after performing feature selection (light green) with p-value cutoff 0.005. To perform feature selection, a separate Cox model was fit to each feature treating it as a predictor of time-until-BCR, while adjusting for the pre-treatment clinical features. Then only features with Wald-test p-value < 0.005 were retained. **d.** The following p-value cutoffs were considered for the feature selection in part b: 0.001, 0.005, 0.01, 0.05 and 1 (1 meaning no feature selection, *i.e.* all features were considered in the model). Notice a random forest model with p-value cutoff 0.005 resulted in the highest C-index in the test dataset. **e.** The best random forest model from Figure 4i was used to predict BCR risk score in the test dataset and median-dichotomized into RF-High vs. RF-Low risk. The Kaplan-Meier plot shows the BCR-free probabilities for a subset of 55 patients classified as intermediate risk according to the National Comprehensive Cancer Network (NCCN) risk categories. The hazard ratio (HR), 95% confidence interval, and Wald-test p-value were estimated from a Cox proportional hazards model. The table below the plot gives the number of patients at risk from 0-5 years. Results for all 161 test patients are shown in Figure 4k.

## Supplementary Information

### Association between self-identified race (SIR) and ancestry

To examine relationships between SIR and genetic ancestry with clinico-molecular and epigenomic features, we compiled publicly available SIR labels and inferred genetic ancestry for samples with germline data (**Extended Figure 1a–c**; **Methods**). We then assessed associations between SIR and genetic ancestry and key features. Most effect sizes were small (Cramér’s V for categorical variables; ε² for continuous variables), with values < 0.1 for 16/18 (89%) of genetic-ancestry associations and 12/18 (67%) of SIR associations (**Extended Figure 1d**, **2–3**). Because some datasets are focused on specific clinical contexts (e.g. primary, metastatic tumours), we expected certain associations. However, overall we found limited evidence that genetic ancestry confounds our analyses, particularly for global methylation measures (methylation subtype, CIMP score, DNAm age acceleration) and global mutation-burden measures (SNVs/Mbp, PGA). Further systematic research on how prostate cancer methylation varies by SIR and ancestry is warranted.

### Tumour purity estimates benchmarking

To investigate whether tumour purity affects the methylation subtypes, we compared four tumour purity estimation modalities (DNA, RNA, methylation, and imaging/pathology-based) spanning ten algorithms across the four methylation subtypes and stratified by cohort (**Extended Figure 4a-d**) (1–8). We observed little difference in tumour purity across the four methylation subtypes for methylation (a), RNA (b) and imaging/pathology-based (c) estimates of tumour purity whereas DNA-based estimates showed modest differences (d). Overall, three of four purity-estimation modalities, particularly imaging/pathology-based estimates, showed minimal differences in purity across subtypes.

### Sensitivity analysis for tumour purity

We next performed sensitivity analyses to assess the impact of tumour purity on 1) subtype classification accuracy, 2) stability of subtype calls after including tumour purity in the classifier and 3) association between the methylation subtypes and key clinico-molecular features (**Extended Figures 5, 6**).

First, for each TCGA sample, we defined misclassification as disagreement between the classifier-predicted subtype and the original TCGA-derived subtype reference label. Then, we tested whether the odds of misclassification varied with tumour purity using a logistic generalized additive model (GAM). The main predictor in the model is tumour purity. The GAM framework allowed for a potentially smooth non-linear (or linear) relationship between odds of misclassification and tumour purity. All types of purity estimators showed no significant relationship with subtype classification accuracy (methylation, RNA, imaging pathology and DNA; p > 0.05) (**Extended Figure 5a**).

Next, we added tumour purity as an additional predictor to the subtype classifier to determine whether the tumour purity-adjusted classifications differ from the original unadjusted classifications (**Extended Figure 5b**). There was little difference between the purity-unadjusted and adjusted subtype classifications, and this result held across all types of purity estimators (**Extended Figure 5c**). Consistent with this result, classifications adjusted for different types of purity estimators were all highly correlated (**Extended Figure 5d**, all Matthews correlation coefficients ≥ 0.87), again showing that tumour purity (and type of purity estimator) had little effect on subtype classification.

Lastly, we assessed whether tumour purity confounded the relationship between methylation subtype and nine important clinico-molecular features (**Extended Figure 6a-i**). For each feature, the original estimate of the effect size of MS-4 *vs.* MS-1 was compared with a purity-adjusted version of the effect size using regression to adjust for tumour purity as a covariate. This sensitivity analysis was repeated for six different tumour purity algorithms. For time-to-event outcomes, we compared the log_2_ fold change of the unadjusted/adjusted hazard ratios; binary outcomes used log_2_ fold change of unadjusted/adjusted odds ratios; and continuous outcomes used a standardized difference in linear regression slopes (the difference is interpreted as the number of outcome standard deviations, similar to Cohen’s D). Only 5 of the 54 sensitivity analyses showed evidence for tumour purity confounding (q < 0.05). For each of the nine outcomes, most of the six algorithms showed no significant evidence of tumour-purity confounding. At most, weak evidence was observed for the hypoxia score and T-category (2/6 algorithms) and for ISUP grade (1/6 algorithm). Importantly, for the two algorithms showing some evidence of confounding (EPIC and CLONET), purity-adjusted hypoxia scores remained statistically significant and directionally consistent after stratification by methylation subtype (q < 0.05). Overall, these sensitivity analyses indicate that associations between methylation subtypes and important clinical and molecular features are robust to potential confounding by tumour purity.

In summary, across four purity-estimation modalities spanning ten algorithms, we observed (i) minimal purity differences across methylation subtypes for three modalities (methylation, RNA, and imaging/pathology-based), (ii) no evidence that tumour purity systematically impacts subtype classification accuracy, (iii) strong agreement between unadjusted and purity-adjusted subtype calls, and (iv) little evidence that tumour purity confounds associations between methylation subtype and key clinico-molecular features.

**Extended Figure 1:**
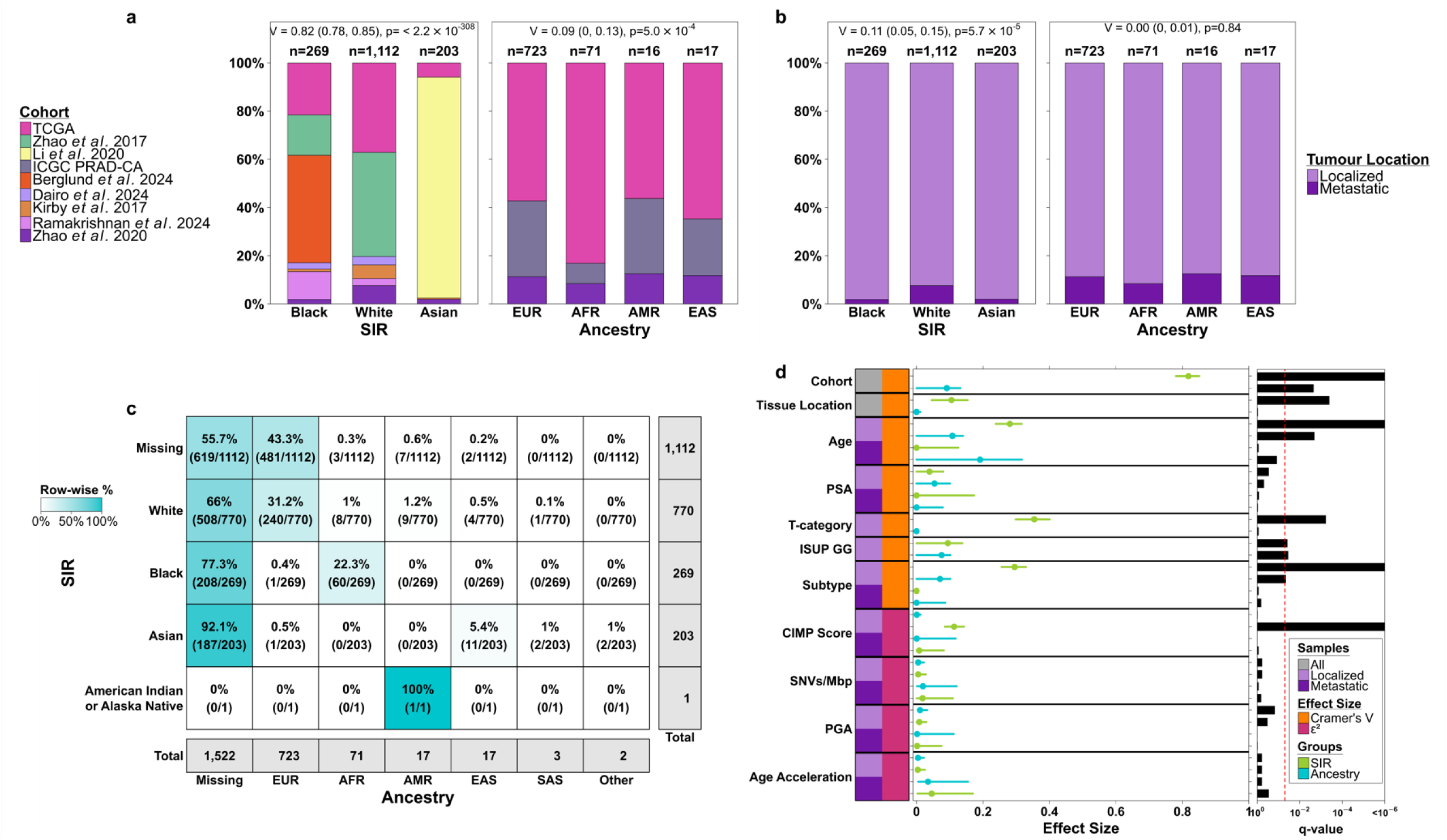
Association between Self-Identified Race (SIR) and genetic ancestry with clinico-molecular features. **a.** Distribution of SIR and ancestry by cohort. **b.** Distribution of SIR and ancestry by tumour location (localized or metastatic). **c.** Contingency table of SIR and ancestry. Cells are colored by row-wise percentages. **d.** Association between SIR and genetic ancestry with clinico-molecular features, stratified by localized and metastatic samples. Bias-corrected Cramér’s V was used for categorical features and epsilon-squared (ε²) for numeric features.

**Extended Figure 2:**
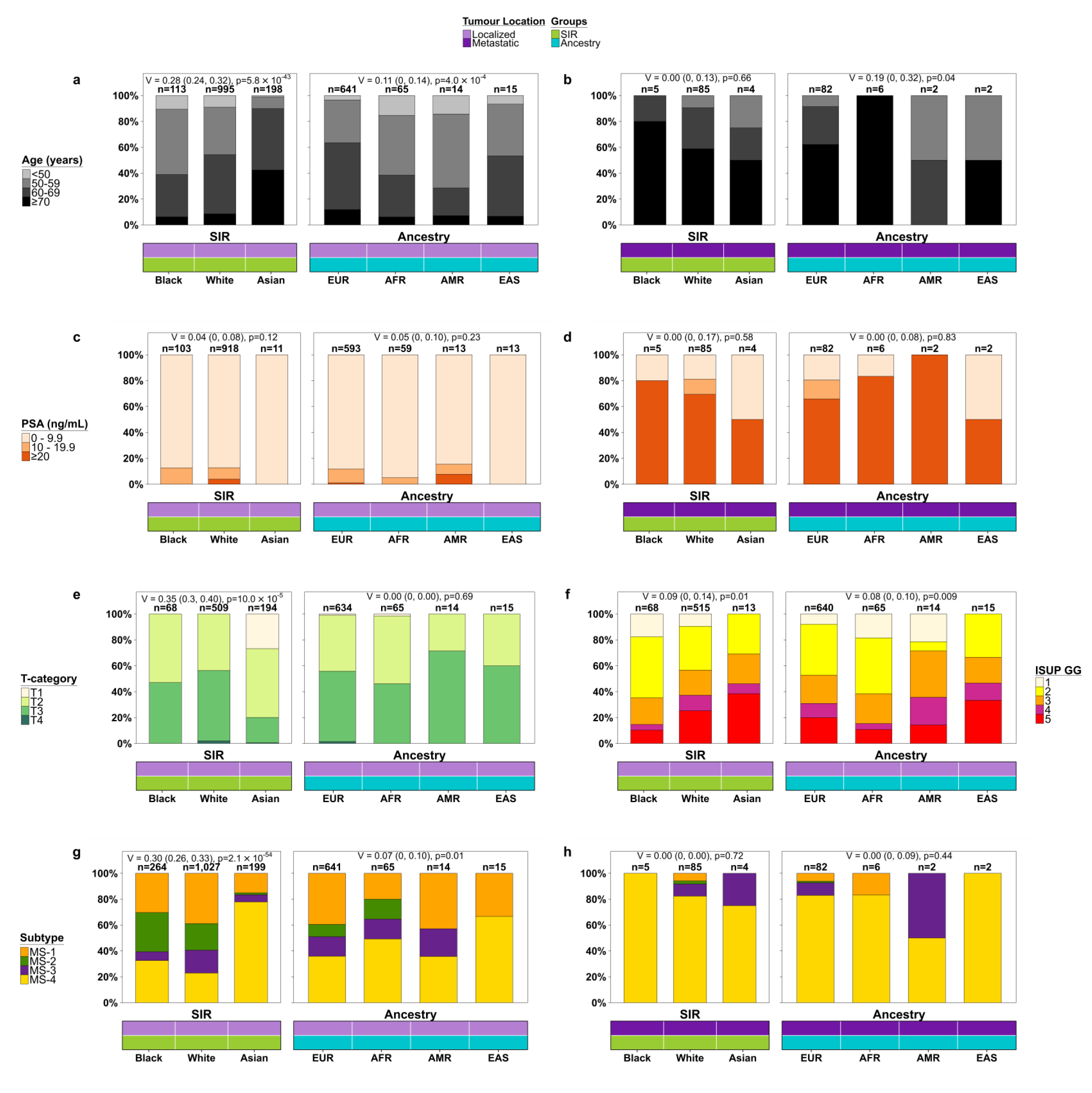
Association between SIR and genetic ancestry with categorical clinico-molecular features. **a-h:** These plots give more detailed visualizations of the relationships between SIR and ancestry with the categorical clinico-molecular features from **Extended Figure 1d**. Results are stratified by localized (left) and metastatic (right) samples. Bias-corrected Cramér’s V effect size with 95% confidence interval and Fisher’s exact test p-value are displayed at the top of each plot.

**Extended Figure 3:**
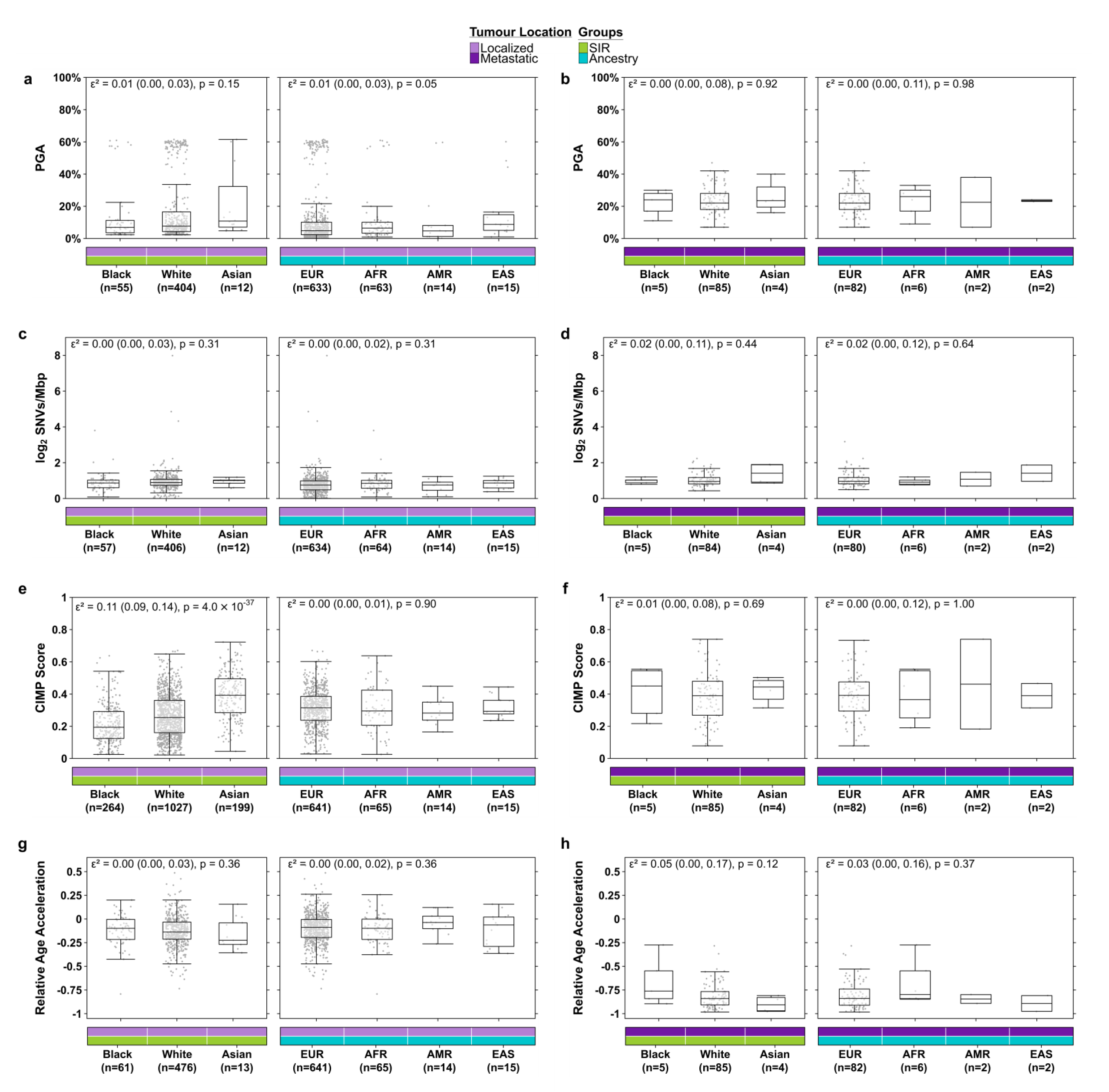
Association between SIR and genetic ancestry with numeric clinico-molecular features. **a-h:** These plots give more detailed visualizations of the relationships between SIR and ancestry with the numeric clinico-molecular features from **Extended Figure 1d**. Results are stratified by localized (left) and metastatic (right) samples. Epsilon-squared (ε²) effect size, 95% confidence interval, and Kruskal-Wallis test p-value are given at the top of each plot.

**Extended Figure 4:**
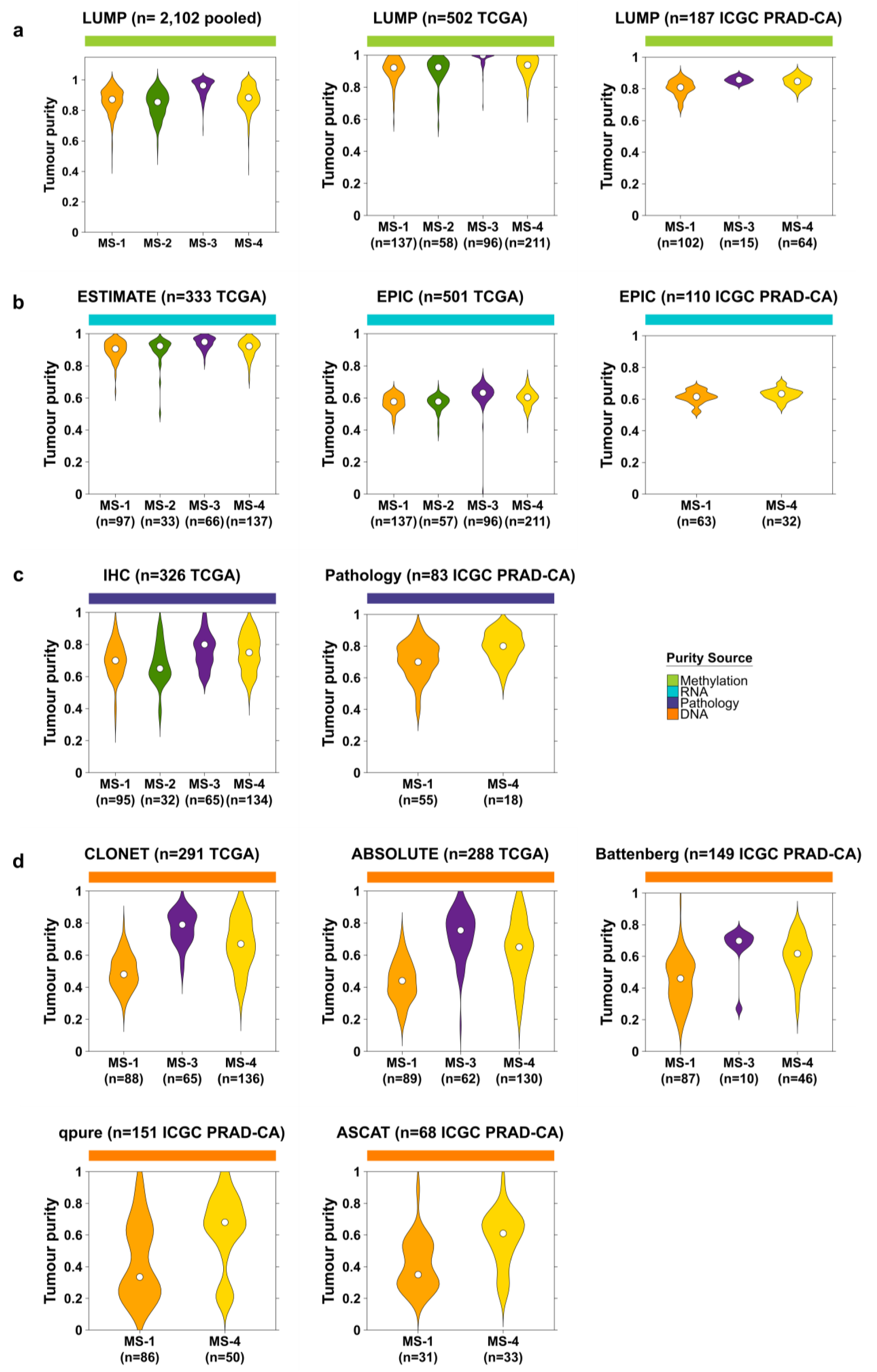
Distribution of tumour purity stratified by methylation subtype, cohort and the methods used to estimate tumour purity: **a.** methylation, **b.** RNA, **c.** imaging/pathology, or **d.** DNA-based methods.

**Extended Figure 5:**
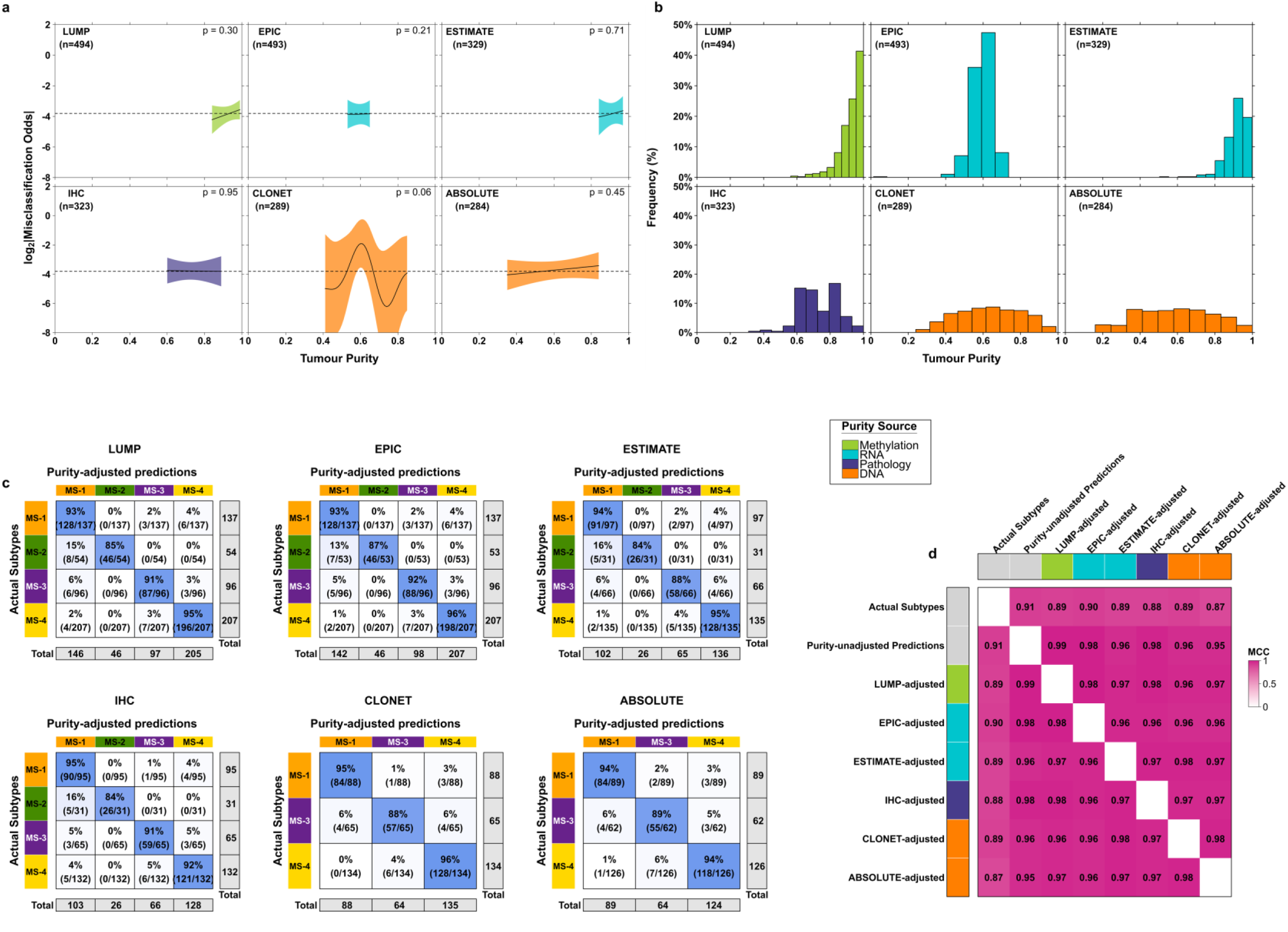
Impact of tumour purity on methylation subtype classification. **a.** A generalized additive model (GAM) was used to estimate the relationship between tumour purity (x-axis) and subtype misclassification (y-axis). Results are stratified by six different types of tumour purity estimators. For a given type of tumour purity, the estimated curve with 95% confidence band is shown for the 10^th^ to 90^th^ percentiles of purity. **b.** Distribution of each type of tumour purity estimator. **c.** Contingency tables of the methylation subtypes derived in the TCGA cohort (y-axis) *vs*. purity-adjusted classifications (x-axis; a new random forest subtype classifier was trained while including tumour purity as a covariate). **d.** Matthews correlation coefficient is shown for pairwise relationships between the original assigned subtypes, purity-unadjusted predicted subtypes, and various types of purity-adjusted predicted subtypes.

**Extended Figure 6:**
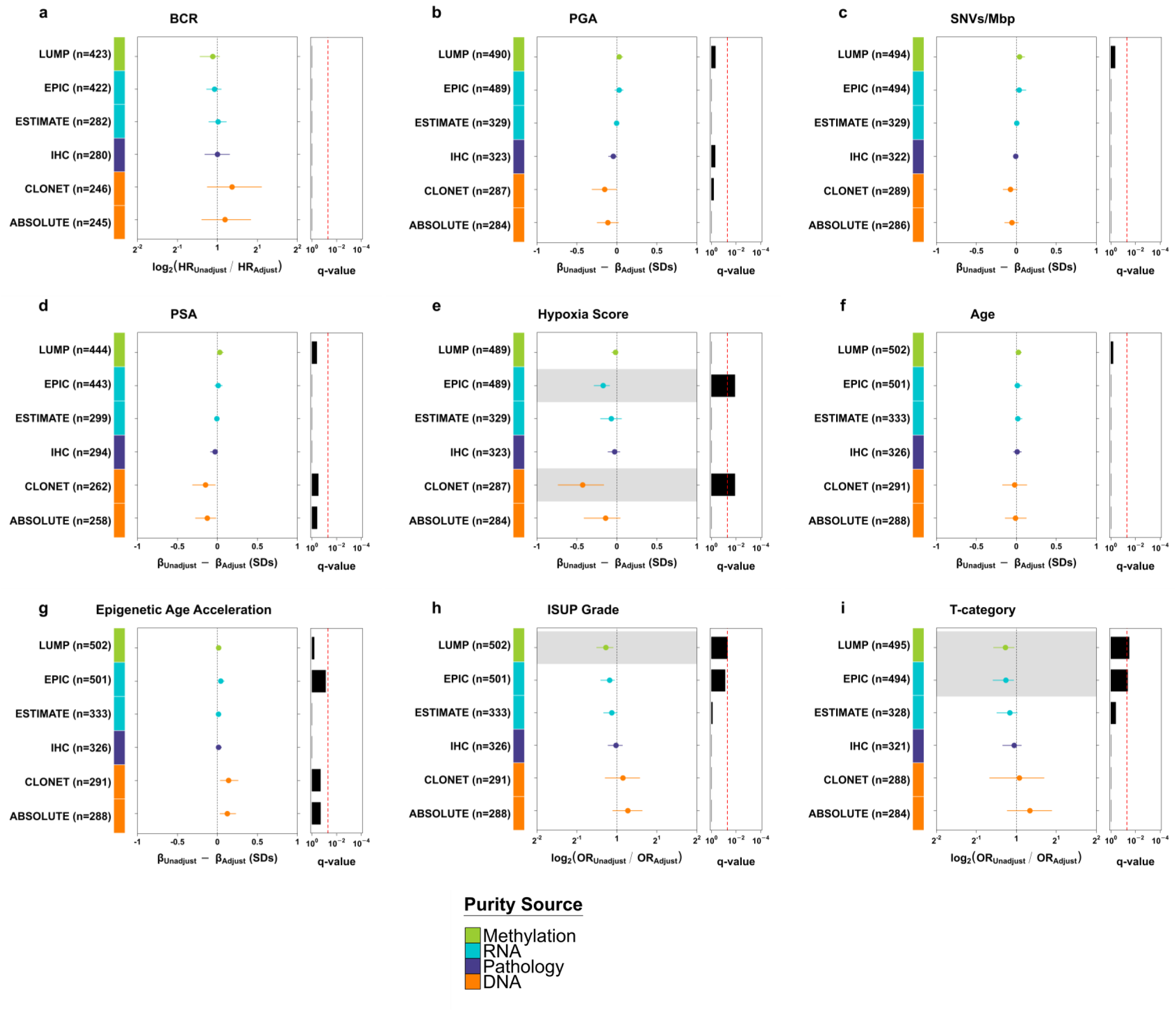
Sensitivity analysis for impact of tumour purity on association between methylation subtypes and key clinico-molecular features. **a-i.** For each feature, the original estimate of the effect size of MS-4 *vs*. MS-1 was compared with a tumour purity-adjusted version of the effect size by using a regression model to adjust for tumour purity as a covariate. This sensitivity analysis was repeated for six different tumour purity estimators. For time-to-event outcomes, we compared the log_2_ fold change of the unadjusted/adjusted hazard ratios; binary outcomes used log_2_ fold change of unadjusted/adjusted odds ratios, and continuous outcomes used a standardized difference in linear regression slopes (the difference is interpreted as the number of outcome standard deviations, similar to Cohen’s D effect size).

